# Parallel Selection for Longevity in Mammals and Birds

**DOI:** 10.64898/2025.12.16.694703

**Authors:** William B. Zhang, Marcus R. Kronforst

**Affiliations:** Department of Pathology, University of Chicago; Department of Ecology & Evolution, University of Chicago

## Abstract

Most studies of aging biology to date have involved the manipulation of short-lived model organisms, while the existing anti-aging mechanisms in naturally occurring long-lived vertebrates have generally remained undiscovered or understudied. The technological advances of the recent “omics revolution” have enabled comparative genomics studies, which have started to unravel genetic signatures of longevity in vertebrates. Building on prior studies and incorporating a novel approach to detecting convergent positive selection, we conducted the first genome-wide survey of positive and purifying selection among hundreds of long-lived mammals and birds, two major vertebrate taxa with notable parallels in their evolutionary history. We discovered an extensive network of shared pathways under purifying selection in both mammals that are exceptionally long-lived for their body size (ELL) and large-bodied long-lived (LLL) birds. In our positive selection survey, we identified 16 genes, involved in eight distinct hallmarks of aging, with concordant signals of positive selection in LLL mammals and LLL birds at neighboring amino acid residues. These included two genes directly involved in cholesterol metabolism, as well as genes whose products clear oxidized metabolites and regulate peroxisomal autophagy. These striking parallels between long-lived mammals and birds, both in broad pathways under purifying selection, as well as in instances of genes under parallel positive selection in LLL mammals and LLL birds, together imply an ancient shared genetic toolkit for longevity, deeply conserved and repeatedly modified to produce longevity in diverse lineages.

**Significance:** While aging is nearly universal in vertebrates, lifespan can vary by several orders of magnitude. Further, extraordinarily long-lived species are dispersed throughout the mammalian and avian phylogenies, indicating repeated evolutionary origins of long life. In the first genome-wide survey of positive and purifying selection in long-lived mammals and birds, we found a striking level of parallelism in the signatures of selection in long-lived species in both clades. These included an extensive network of shared pathways under purifying selection, as well as several novel genes under parallel positive selection. Our results identify multiple novel candidate pathways and genes for further study, and provide evidence for the convergent evolution of extended lifespan.

## 1. Introduction

In vertebrates, aging is a nearly universal degenerative process characterized by time-dependent physiological decline [1, 2]. While the amelioration of its effects would be of immense economic and public health interest to humans, who are already comparatively long-lived at baseline, studies of the biology of aging have traditionally relied on the manipulation of short-lived model organisms for pragmatic reasons [3, 4]. However, the magnitude of lifespan changes observed within species is dwarfed by the magnitude of lifespan variation across species, where even within vertebrates, maximum lifespan can range from just 59 days in the pygmy goby, *Eviota sigillata* [5], to approximately 400 years in the Greenland shark, *Somniosus microcephalus* [6]. Several evolutionary theories of aging have been put forth to explain this vast diversity in naturally-occurring lifespans, proposing that selective pressures modulating lifespan include extrinsic mortality (e.g., limited nutrient supply, predation, and disease) and life history traits such as body mass and age at reproductive maturity [7, 8].

The repeated origin of long lifespans across the tree of life, in multiple distinct lineages, suggests that natural selection has shaped vertebrate genomes to create convergent cellular and molecular mechanisms of longevity [9, 10]. While the genetics of convergent evolution have been studied in specific morphological traits, such as flight membrane formation in flying mammals [11] or pigmentation in butterflies and lizards [12], they are not well-understood in the context of longevity, which is a more abstract, multidimensional, and emergent trait. Previous work has established that purifying selection in certain critical pathways may be required for the development of long lifespans, maintaining existing longevity-enabling functionality by purging deleterious changes [13, 14]. In concert with the prerequisite purifying selection, which preserves existing longevity-enabling pathways with high fidelity, positive Darwinian selection, in which adaptive variation spreads through a population, acts to produce longevity by favoring specific longevity-establishing innovations. These forms of selection, and the pro-longevity mechanisms on which they act, can be readily measured in coding sequences using established methods of comparative genomics [15–18]. With the advent of long-read genomic sequencing technologies and the proliferation of high-quality assemblies [19, 20], comparative approaches identifying cross-species genomic footprints for longevity are now possible. Indeed, recent analyses within groups of mammals [13, 21, 22], rockfish [14, 23], and birds [24, 25] have uncovered several novel longevity-associated genetic signals.

In order to better prioritize candidate mechanisms and genes for further study, it is necessary to first determine which mechanisms of longevity in non-human models are most suitable for translation. Recently, there has been interest in a distinction between public (those which are shared between multiple long-lived species) and private (those which are unique to one species) mechanisms of longevity [26]. Specifically, public mechanisms of longevity may act on highly conserved genetic toolkits, and as a result may be more directly applicable to human biology. Mammals and birds are ideal for identifying these types of convergent longevity mechanisms, as the two vertebrate clades have large variation in lifespan and body size, are geographically widespread, and inhabit diverse ecological niches [9]. Both clades experienced rapid diversification around the Cretaceous-Paleogene boundary approximately 66 million years ago, with placental mammal and neoavian most recent common ancestors (MRCA) estimated to have lived around that time [27, 28] or somewhat earlier [29, 30]. In addition, while humans are ourselves mammals, birds, as amniotes, are also relatively closely related. Due to the parallels and similarities between these two groups, the systematic comparative study of mammals and birds promises to identify novel public longevity mechanisms that are particularly amenable to intervention.

Here, we performed an unbiased, genome-wide survey of protein-coding sequences to identify convergent evolutionary signatures for longevity across hundreds of species of placental mammals and birds. In contrast to previous studies, we separately analyzed two major vertebrate taxa and identified explicit genomic parallels in their longest-lived members. Furthermore, our positive selection analysis used a novel approach: We divided the wider sets of species into 7 mammalian and 9 avian clades, explicitly surveying signals of selection in the longest-lived species *within* each clade to minimize confounding signals due to shared ancestry among long-lived species. We then compared results across clades to discover mechanisms of longevity with support from multiple clades representing independent natural experiments occurring over millions of years. We discovered a striking level of parallelism in pathways under increased constraint in both vertebrate clades, as well as several candidate genes with signatures of evolutionary innovation in both long-lived mammals and long-lived birds.

## 2. Results

### 2.1. Longevity Variation Among Mammals and Birds

First, we characterized the extent of phenotypic variation in our dataset. Among 295 species of mammals and 194 birds with sufficient genetic and phenotypic data in the form of genomic sequences, body mass measurements, and maximum longevity data (Figure 1A), we examined the previously characterized [31] relationship between body size and lifespan (Figure 1B). We observed significant linear associations (*p <* 10*^−^*^28^ for mammals; *p <* 10*^−^*^20^ for birds) between log adult weight and log maximum longevity in both mammals and birds, with birds having generally longer lifespans at a given body size as well as a slightly steeper slope in the fit line, increasing lifespan by a greater extent for an equal increase in body mass. This is consistent with prior studies of the association between extrinsic mortality, and particularly flight ability, on lifespan [8].

**Figure 1:**
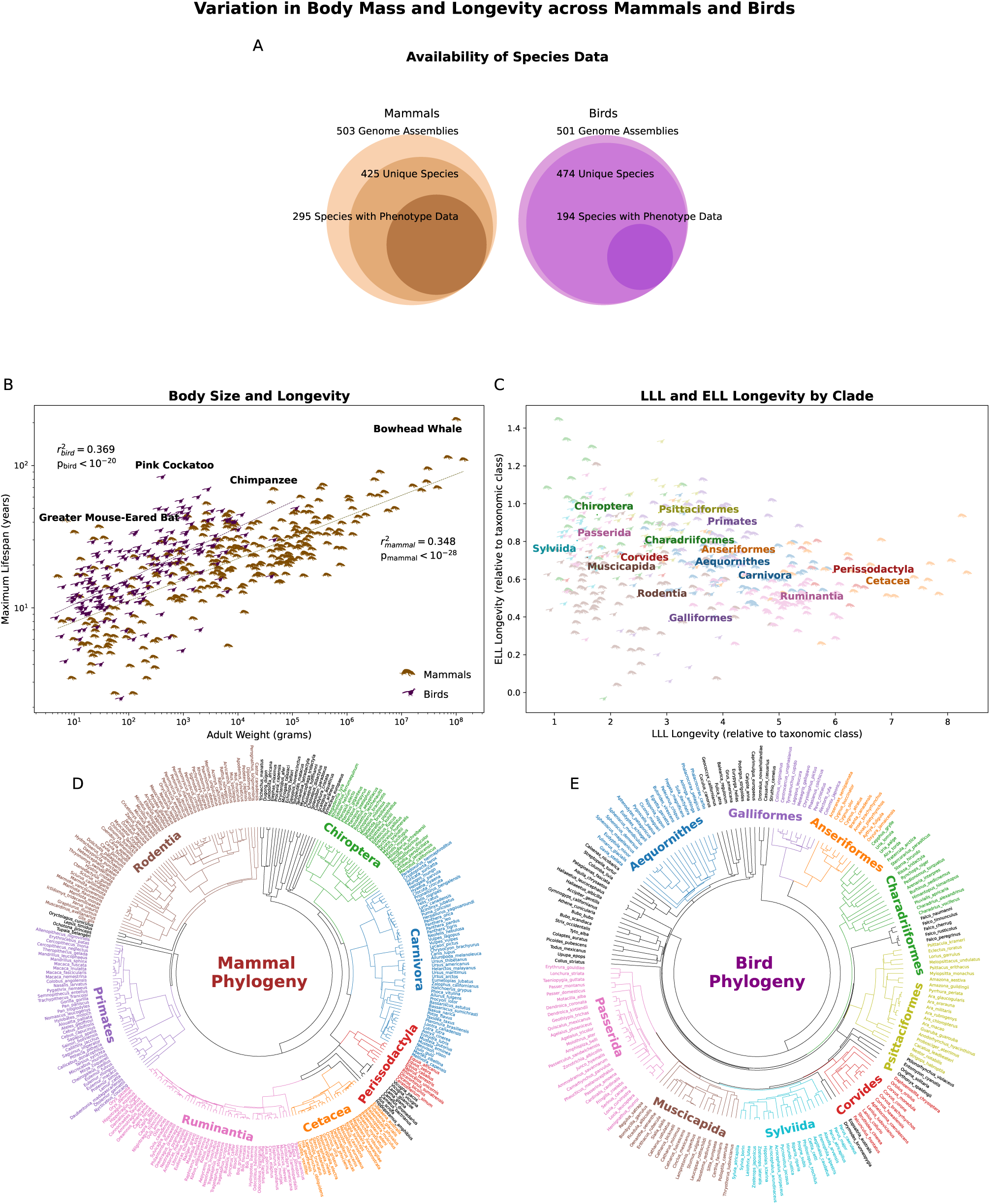
Existing phenotypic variation and ancestral relationships among mammals and birds. (A) Starting from approximately 500 genome assemblies in each taxonomic class, 295 mammalian and 194 avian unique species with available phenotype data were selected. (B) The relationship between body size and longevity within mammals and birds (note the log scales on both axes). (C) Significant variation in LLL and ELL Longevity within and among diverse clades of mammals and birds. (D) A mammalian phylogeny of 295 species with 7 distinct clades highlighted. (E) An avian phylogeny of 194 species with 9 distinct clades highlighted.

We can thus distinguish two distinct forms of longevity, one that varies in a manner correlated with adult weight, and one that is perpendicular to the main association, in which a species may be exceptionally long-lived for its body size. As has been described previously [13], we defined two geometrically orthogonal phenotypes of longevity: large-bodied and long-lived (LLL), and exceptionally long-lived (relative to body size; ELL; Figure 1C). We then selected 7 mammalian and 9 avian clades (Figure 1D–E) from within our full sets of species, and found that while clades have characteristic values of LLL and ELL (e.g. rodents tend to be smaller and thus low in LLL longevity), there was also substantial internal variation in longevity within each clade (Figure 1C), illustrating that lifespan has been repeatedly tuned in several independent mammal and bird clades.

### 2.2. Experimental Overview

After establishing the existing variation across and within mammal and bird clades, we conducted the first large-scale survey of purifying and positive selection in mammals and birds. In order to target genetic sequences with high conserved function across species, our analysis focused on the coding sequences of one-to-one orthologous genes. We did so separately for long-lived species exhibiting high values of the distinct LLL and ELL phenotypes.

In our purifying selection analyses, we first selected 195 mammal and 184 bird species (Tables S1–S25) with sufficient one-to-one orthology to the reference genome to satisfy our method’s technical requirements (Figure 2A). In a manner previously described [13], we computed gene trees from TOGA-aligned coding sequences [32] of species predicted to have one-to-one orthologs for each particular gene (Figure 2B). Then, we used RERConverge [16, 17] to correlate rates of genotype and phenotype (LLL or ELL) evolution for each gene and to identify genes under increased purifying selection (Figure 2C). Lastly, we aggregated sets of genes into pathways to understand the larger biological processes under constraint (Figure 2D). In our positive selection analyses, we developed a new approach in order to detect convergence among independently evolved examples of organismal longevity by separating our species into several clades (Figure 2E). For each gene, species within a clade with one-to-one orthology to the reference were selected, and species in the top quartile of the phenotype (LLL or ELL) were chosen as the foreground (Figure 2F). Positive selection was then assessed in codeML [15] by maximum likelihood comparisons of models with and without positive selection (Figure 2G), producing an array of p-values for each clade and gene within each vertebrate class and longevity phenotype (Figure 2H).

**Figure 2:**
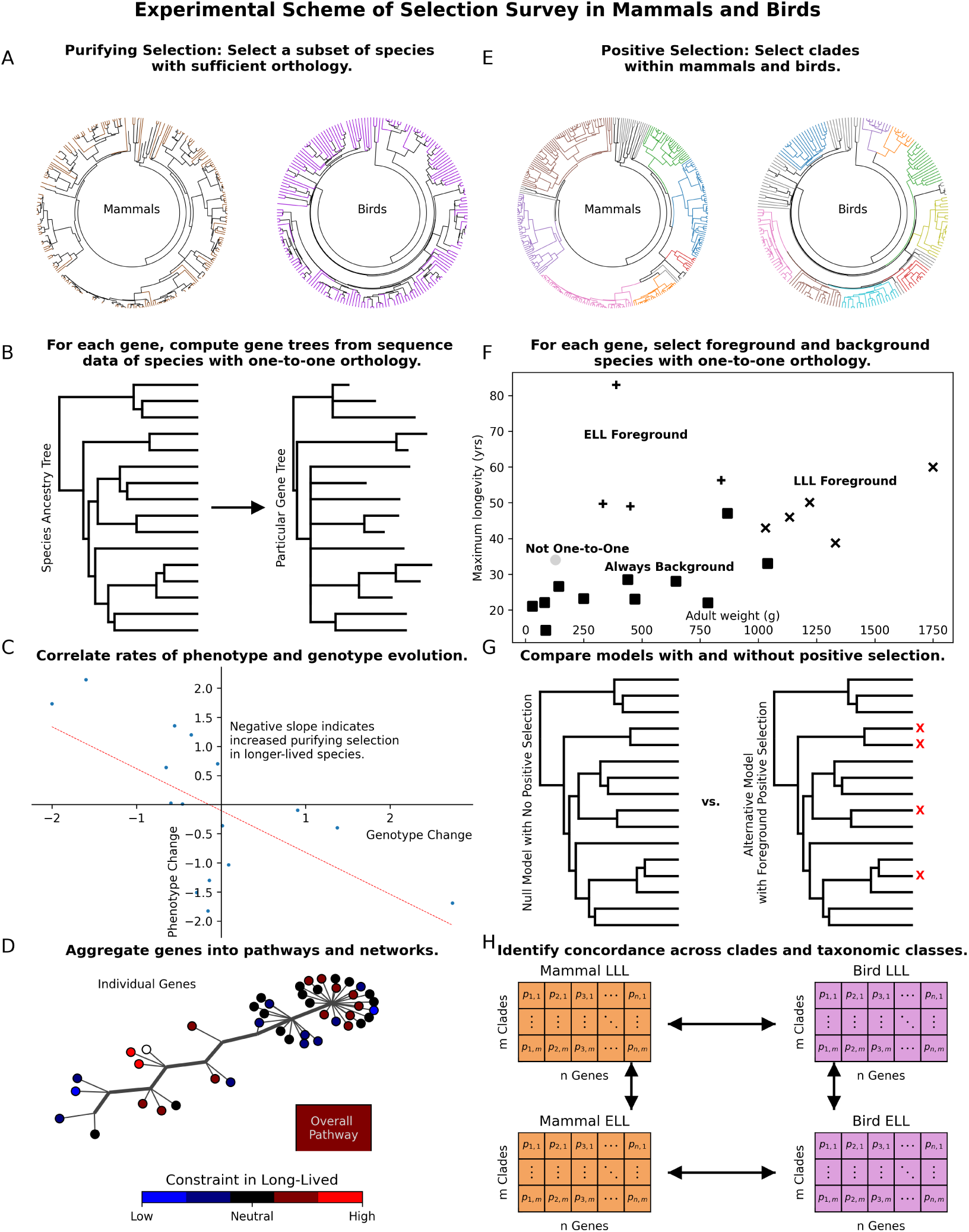
Experimental scheme of a genome-wide survey of purifying and positive selection for two types of longevity in mammals and birds. (A) Starting from approximately 500 genome assemblies in each taxonomic class, 295 mammalian and 194 avian unique species with available phenotype data were selected. (B) Species with insufficient orthology to compute a master gene tree were omitted from the purifying selection analysis. (C) As a basis for determining relative evolutionary rates, gene trees were created from species ancestry topology with branch lengths computed from aligned one-to-one orthologous protein-coding sequences. (D) The relative evolutionary rate of a gene in long-lived species was determined by the Pearson correlation between inferred rates of phenotype change and genotype change in each branch. (E) To understand broader pathways under purifying selection, we aggregated individual genes into higher levels of organization. (F) The positive selection analysis subdivided each taxonomic class into several distinct clades. (G) Within each clade, the top quartile of species in LLL or ELL longevity was selected as the foreground. For each gene, only species with one-to-one orthologs of that gene were included. (H) Using a maximum likelihood test in CodeML, a null model with only neutral and negative selection was compared to an alternative model which allows positive selection in foreground branches. (I) Intra-and inter-class comparisons were made among clade results to determine which candidate genes have the widest support as contributors to species longevity.

### 2.3. Enrichment of Selection Signals

We then analyzed our results for overall enrichment in significant gene-level signals compared to a background of re-shuffled species phenotypes. This allowed us to verify that any signal we observed was due to an actual genotype-phenotype correspondence, and not due to confounding properties of the aligned sequences (e.g. gap content of genetic sequence, Figures S1 and S2).

In our purifying selection results (Figure 3A–D), the null background was largely as expected. The distribution of p-values produced by 100 permulations [33] applied genome-wide closely approximated a uniform distribution, with some reduced density at the lowest p-values. We observed significant enrichment in lower p-values for the mammal LLL (Figure 3A, *p <* 10*^−^*^30^), bird LLL (Figure 3C, *p <* 10*^−^*^60^), and bird ELL (Figure 3D, *p <* 10*^−^*^14^) analyses. For the mammal ELL analysis, lower p-values at the gene level were not significantly enriched (Figure 3B).

**Figure 3:**
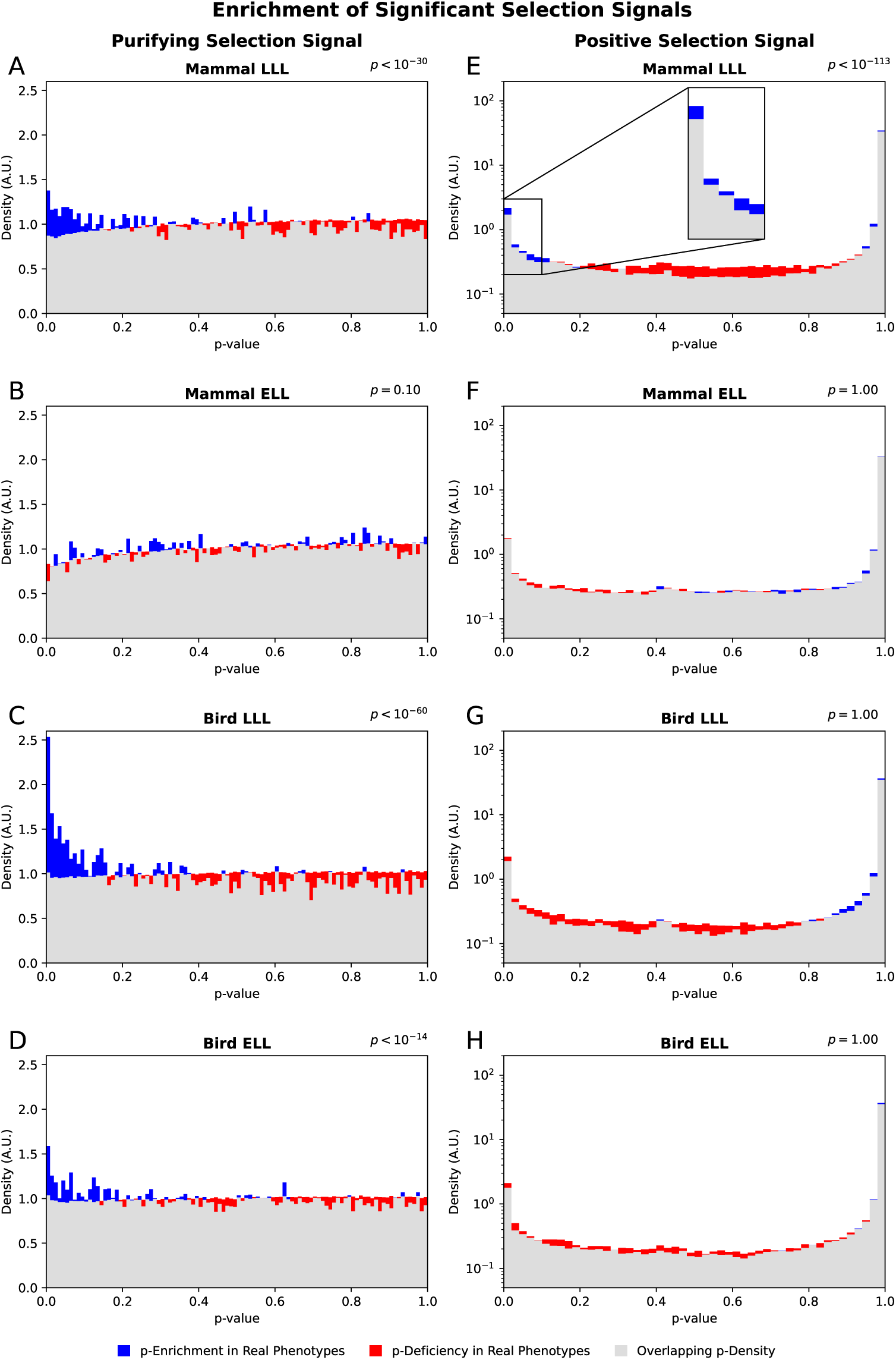
Enrichment of significant p-values in true genotype-phenotype assignments over a null background of analyses using re-shuffled phenotypes: Purifying selection results for (A) LLL mammals, (B) ELL mammals, (C) LLL birds, and (D) ELL birds. Positive selection results for (E) LLL mammals, (F) ELL mammals, (G) LLL birds, and (H) ELL birds. Note that due to the logarithmic scale, the blue and red regions in E–H are not equal in area.

In our positive selection results, the null background deviated substantially from a uniform distribution, with very large excesses of p-values at both extremes of the distribution (Figure 3E–H). Among the positive selection analyses, only the mammal LLL results showed an enrichment in low p-values over background (Figure 3E, *p <* 10*^−^*^113^). Within this LLL mammals analysis, the enrichment of more significant p-values was present in multiple mammal clades, including Carnivora, Cetacea, Chiroptera, Perissodactyla, and Rodentia (Figures S3–S7). On the other hand, Primates and Ruminantia did not show a broadly enriched signal of positive selection over background (Figures S8 and S9). Taken together, these findings indicated a consistent signal of positive selection in mammalian LLL lineages across multiple independent clades.

### 2.4. Purifying Selection Pathways

Despite an overall pattern of excess low p-values, few genes in the purifying selection analyses were significant after correcting for multiple hypothesis testing (data not shown). However, after aggregating gene-level statistical signals into more robust pathway-level results, we formed networks of connectivity that highlighted the broader biological processes involved. We found that LLL mammals showed increased constraint in growth factor signaling and cytokine signaling, two major networks of pathways related to body size and cancer regulation (Figure 4A). ELL Mammals also showed increased constraint in several cancer-associated groups of pathways: namely networks related to the cell cycle and to cancer itself. ELL mammals also demonstrated increased purifying selection in fundamental cellular functions related to genomic and proteomic fidelity: mRNA processing and nucleotide excision repair (Figure 4B). LLL birds showed a similar pattern of constraint to ELL mammals, with one large network composed of pathways relating to mRNA processing, cell cycle, and cancer, along with smaller groups of pathways related to intracellular trafficking, apoptosis, and sumoylation (Figure 4C). ELL birds showed only two connected pathways under significant constraint (Figure 4D), which are related to the the regulation of methylation of lysine 27 on histone 3 (H3K27), and represent critical regulators of organismal development and stem cell self-renewal [34].

**Figure 4:**
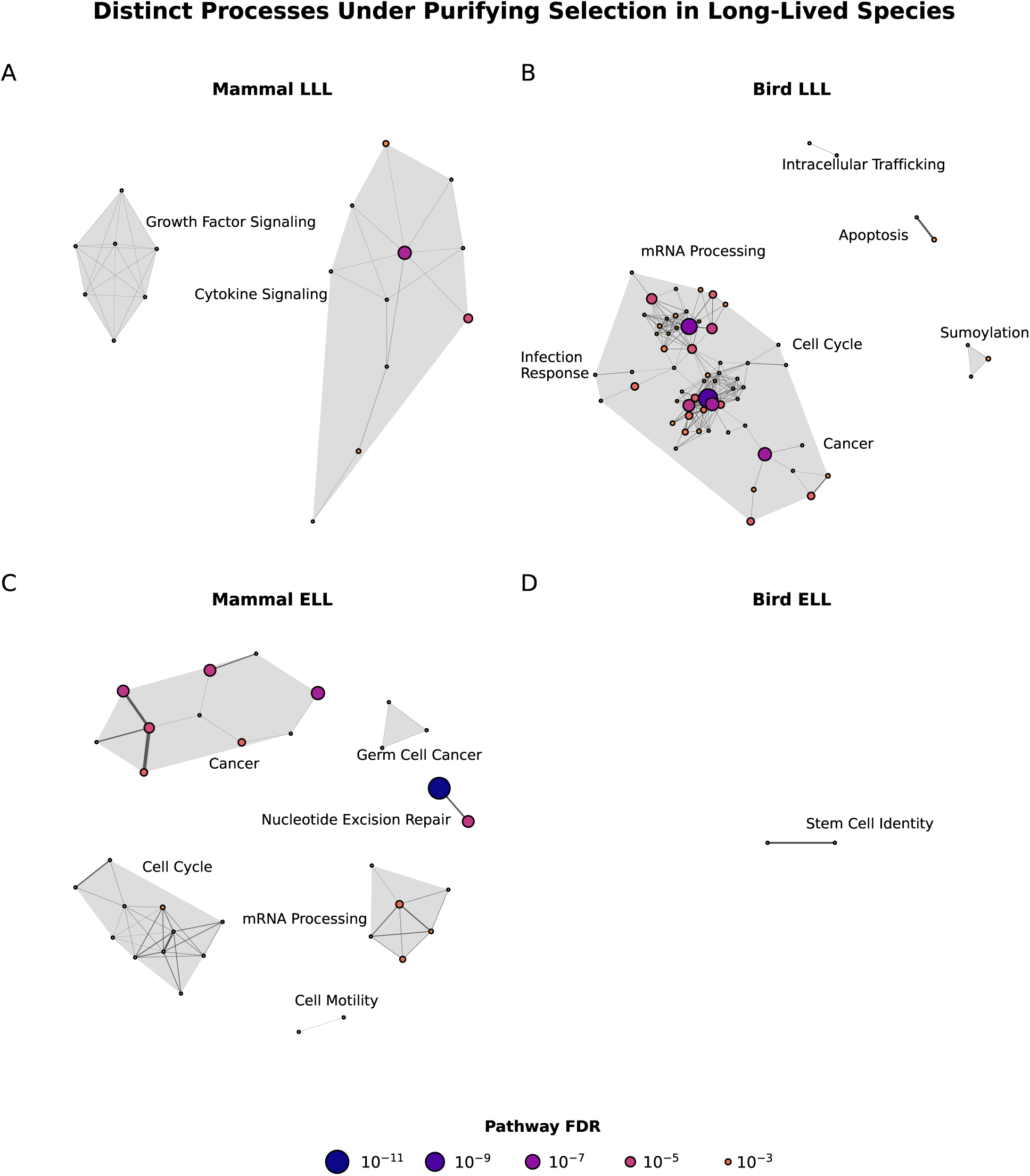
Networks of connected pathways under purifying selection. (A) LLL mammals require fidelity in their cytokine and growth factor signaling to develop longevity. (B) ELL mammal longevity is associated with constraint in pathways related to cancer, cell cycle, mRNA processing, DNA repair, and cell motility. (C) Similarly, LLL bird longevity is associated with constraint in pathways related to mRNA processing, cell cycle, and cancer, as well as unique groupings of pathways related to infection response, sumoylation, apoptosis, and intracellular trafficking. (D) ELL bird longevity is associated with constraint in pathways related to stem cell identity and self-renewal.

### 2.5. Concordance Across Clades

Lastly, we sought to identify and prioritize mechanisms of longevity with broad support from multiple unrelated clades by focusing on the constrained pathways in common between multiple analyses, and the positively selected genes supported by both LLL mammals and LLL birds.

We noted in our purifying selection analyses a significant overlap in the processes under increased constraint in ELL mammals and in LLL birds, and therefore identified a set of pathways with significantly lower rates of sequence evolution in both ELL mammals and LLL birds. These significantly constrained pathways were in the categories of mRNA processing, cell cycle, and cancer (Figure 5A), indicating shared mechanisms of cancer control and accurate information transfer from genome to proteome, maintained in parallel under high fidelity in long-lived members of both groups. Examining the individual constituent genes of the Reactome mRNA splicing pathway [35] demonstrated remarkably widespread parallel constraint at a lower level of genetic and biochemical organization in both ELL mammals (Figures 5B and S10) and LLL birds (Figures 5C and S11).

**Figure 5:**
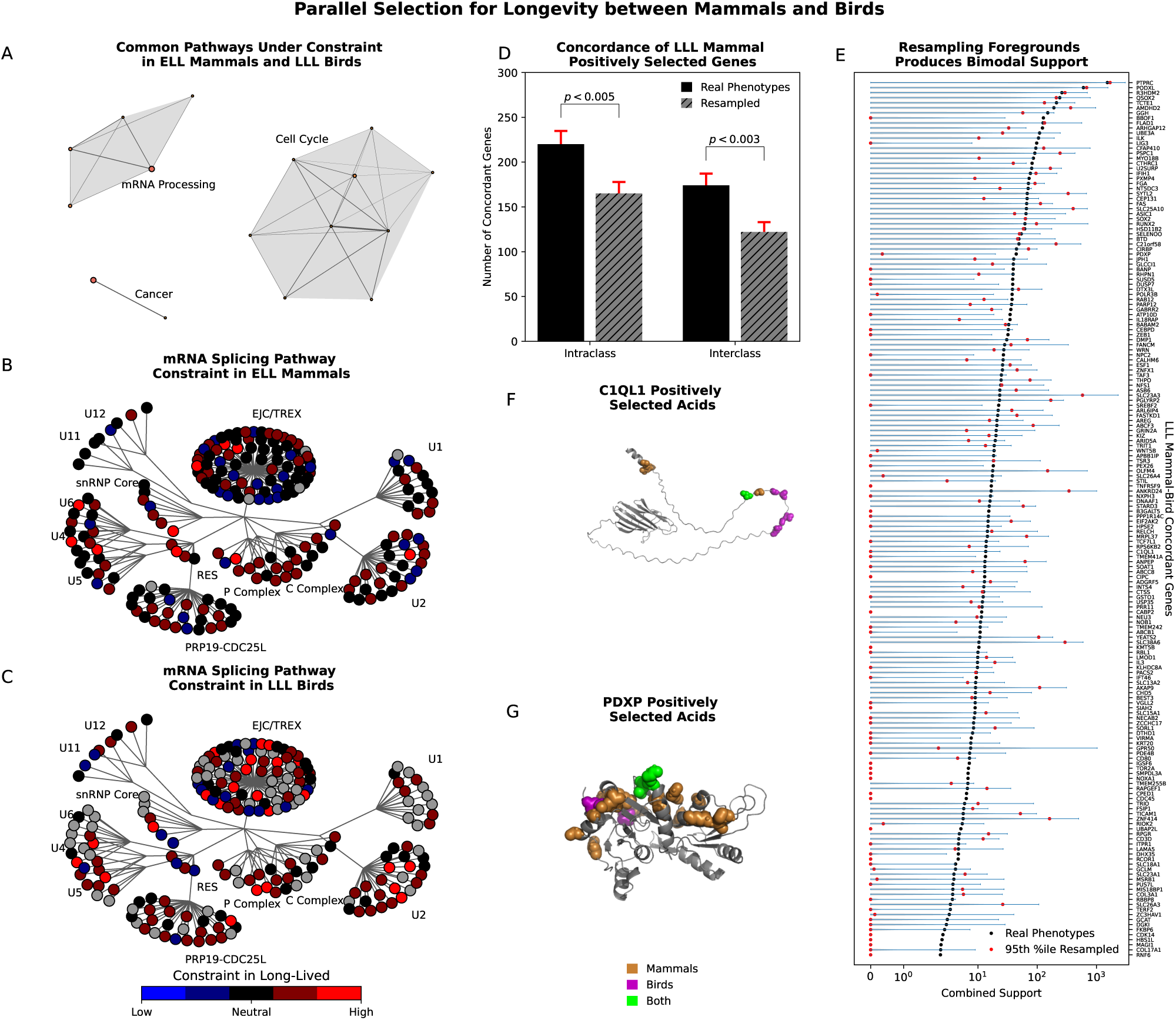
Parallel signals of selection for longevity between mammals and birds. (A) Several connected pathways related to mRNA processing, cell cycle, and cancer are under increased purifying selection in both ELL mammals and LLL birds. (B) Gene-level constraint in ELL mammals for the Reactome mRNA splicing pathway. (C) Gene-level constraint in LLL birds for the Reactome mRNA splicing pathway. (D) There is an increased number of concordant genes over background, both within LLL mammals (intraclass) and between LLL mammals and LLL birds (interclass). (E) Resampling the interclass concordant genes results in a bimodal distribution of 95^th^ percentile support among the resampled runs. (F) Parallel positively selected amino acids in C1QL1 and (G) PDXP.

Next, we examined the 1935 genes with significant signals of positive selection (*FDR <* 0.05) in at least one clade of LLL mammals. To determine which candidate genes were more specifically tied to the longest-lived species and therefore more robustly associated with longevity itself, we defined additional criteria: First, we emphasized genes with support from multiple clades by examining concordance both within LLL mammalian clades and between LLL mammals and LLL birds. Our true phenotypes analysis retained an increased number of significant genes above background in both comparisons (Figure 5D). As the latter is a striking level of parallelism between two major taxa separated by approximately 300 million years of evolution, we focused on the 174 genes with signals of positive selection in both LLL mammals and LLL birds. Second, we performed 100 rounds of phylogenetically-informed resampling of the foregrounded species for our candidate genes, and defined a “combined support” metric (see Methods) to compare statistical support for positive selection in long-lived species to the level of aberrant background signal for positive selection. Interestingly, a bimodal distribution of 95^th^ percentile combined support was observed (Figure 5E), suggesting that some of the mammal-bird concordant genes’ positive selection signals—namely those in the higher mode—are not specific to the true genotype-phenotype pairing. Therefore, we selected the 92 genes in the lower mode, which showed signals of positive selection specific to the selection of long-lived foreground branches.

In our third and final filtering step, we emphasized positive selection at orthologous sites within each protein. To compare across birds and mammals, we aligned the bird reference sequence (*Gallus gallus*) to the mammalian sequences for each gene, and translated all bird amino acid chain numbers to a common mammalian numbering. This necessarily resulted in the loss of bird-specific insertions (relative to the mammalian sequence), but allowed us to compare the locations of orthologous residues. We then classified the 92 lower-mode genes according to the chain length between the closest positively selected mammalian and avian amino acids. We found 2 genes for which an identical amino acid showed a signal of positive selection in both a mammal and a bird clade (Figure 5F and G), and 14 additional genes in which positively selected mammalian and bird residues were within 20 amino acids of each other in the chain (representing acids possibly in or around the same active site [36]), for a total of 16 candidate genes with evidence of positive selection at orthologous sites in both LLL mammals and LLL birds.

These 16 genes—positively selected at orthologous sites in both long-lived mammals and long-lived birds—are novel candidates that have not generally been well-studied in the context of aging, yet the pathways and biological processes in which they are involved are closely tied to known “hallmarks of aging” [2] across multiple biological scales (Table 1 and Figure 6).

**Figure 6:**
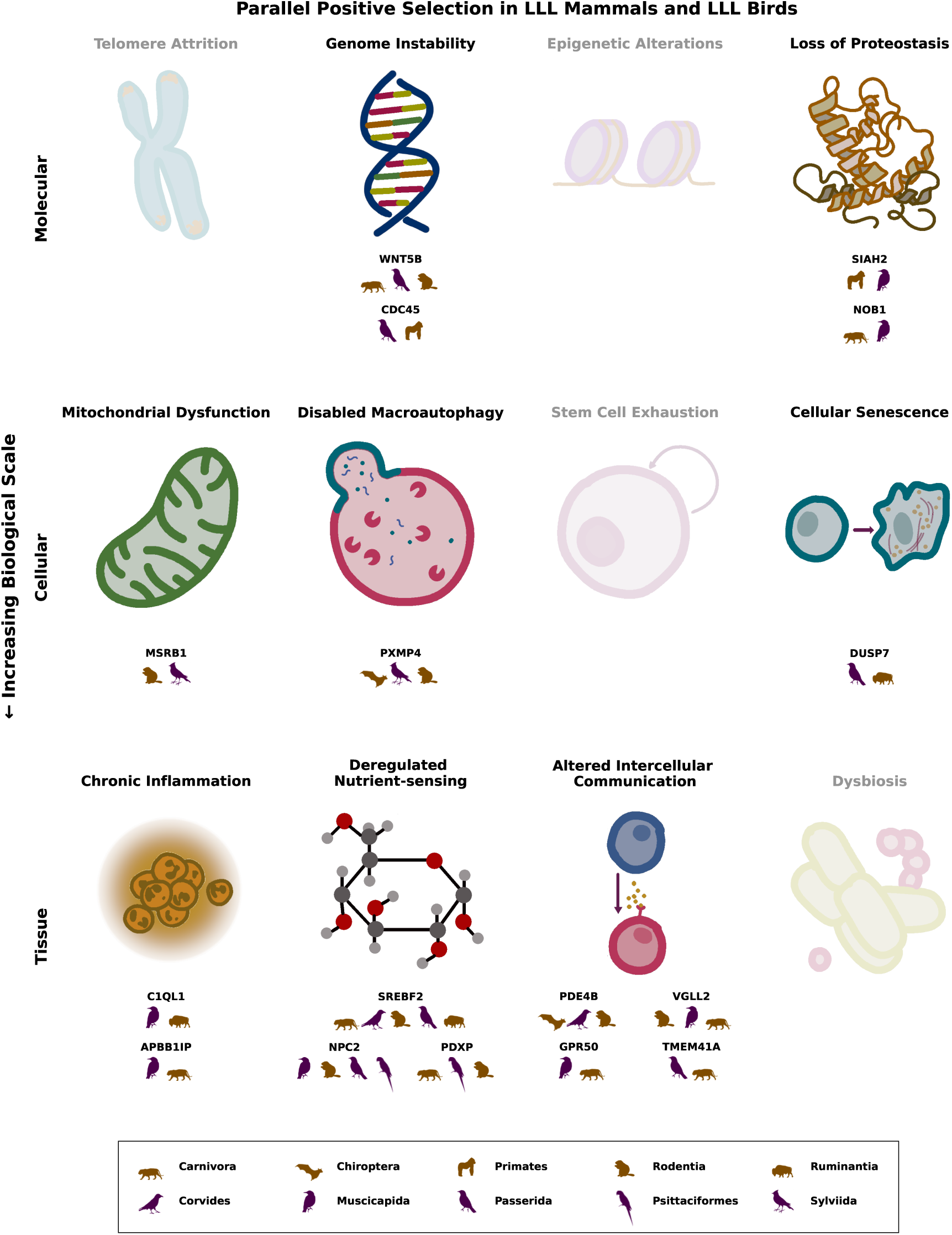
Parallel signals of positive selection and the hallmarks of aging. Genes with shared signals of positive selection between LLL mammals and LLL birds were found to be involved in a number of the hallmarks of aging across the molecular, cellular, and tissue scales.

**Table 1:** Genes positively selected at orthologous sites in both LLL Mammals and LLL Birds.

| Gene Symbol | Functional Connection to Hallmarks of Aging |
| --- | --- |
| C1QL1 | involved in angiogenesis [37] and is related to C1q, which initiates the classic complement pathway of innate immunity [38] |
| PDXP | phosphatase involved in the processing of vitamin B6 to its active form [39] |
| TMEM41A | relatively unstudied transmembrane protein which has been occasionally characterized as a cancer biomarker [40] |
| APBB1IP | interacts with amyloid beta precursors [41] and functions in integrin activation and adhesion of leukocytes [42] |
| CDC45 | component of the eukaryotic replicative helicase CMG [43] |
| DUSP7 | dual-specificity phosphatase known to act on MAPK, one of the key regulators of cellular senescence [44] |
| GPR50 | G-protein coupled receptor with a role in mitophagy [45] |
| MSRB1 | enzyme that reduces an oxidized metabolite back to methionine, countering the effects of oxidative stress and mitochondrial leakage [46] |
| NOB1 | involved in pre-rRNA processing to produce the 18S rRNA of the small ribosomal subunit [47] |
| NPC2 | lysosomal protein involved in cholesterol transport [48, 49] |
| PDE4B | major metabolizer of cyclic AMP, which acts as a second messenger to regulate cellular functions [50] |
| PXMP4 | peroxisomal membrane protein, but has been shown to play a role in lipid metabolism [51] and the regulation of peroxisome autophagy [52] |
| SIAH2 | E3 ligase involved in ubiquitination and proteasome-mediated degradation of various proteins [53, 54] |
| SREBF2 | key transcription factor regulating cholesterol and fatty acid synthesis [55] |
| VGLL2 | transcription cofactor which acts in skeletal muscle development [56] |
| WNT5B | member of the Wnt family, which is canonically implicated in oncogenesis of many tumor types and has been shown to regulate genomic stability *in vitro* [57] |

At the molecular level, we found parallel changes in long-lived birds and mammals in genes related to genome instability and to loss of proteostasis. Interestingly, with regard to proteostasis, we found NOB1, which is involved in the fidelity of protein synthesis, playing a role in the production of a ribosomal subunit, as well as SIAH2, which is involved in marking proteins for turnover and degradation. At the cellular/organelle level, we found evidence of positive selection in genes directly related to mitochondrial function, autophagy, and cellular senescence, including an enzyme that reduces oxidized metabolites back to methionine (MSRB1), a peroxisomal membrane protein that regulates peroxisome autophagy (PXMP4), and a dual-specificity phosphatase known to act on key regulators of cellular senescence (DUSP7). Finally, the majority of our candidate genes were related to mechanisms at the cellular level, involving inflammation, nutrient sensing and processing, and intercellular communication. In particular, we identified two candidate genes in cholesterol metabolism: SREBF2, a transcription factor regulating its synthesis, and NPC2, a protein involved in its export from lysosomes. Notably, our top candidates did not include any genes related to telomere attrition, epigenetic alteration, stem cell exhaustion, or dysbiosis. Taken together, these positively selected genes offer an intriguing glimpse into the evolutionary innovations that have repeatedly given rise to extended lifespans in vertebrates.

## 3. Discussion

In this study, we completed a genome-wide survey of purifying and positive selection in long-lived mammals and birds. We employed a combination of established methods and a novel approach to detect convergent positive selection. Our positive selective strategy emphasized screening for candidate genes within smaller clades of animals to minimize the confounding effects of shared ancestry among long-lived species (e.g. signals unrelated to longevity from consistently long-lived clades such as cetaceans) while conducting a wide-ranging survey of hundreds of species. We subsequently collated results across 7 mammal and 9 bird clades to identify parallel signals in common among multiple clades. We identified striking genetic parallels between long-lived mammals and birds, both in terms of broad pathways under purifying selection in both ELL mammals and LLL birds, as well as in focal examples of genes under parallel positive selection in LLL mammals and LLL birds.

In terms of purifying selection, we identified several foundational processes—including cancer suppression, DNA repair, mRNA processing, and maintaining stem cell identity—that appear to be prerequisites for the development of extended lifespans in different contexts in vertebrates. We have also confirmed, using a larger set of species, previous reports of increased constraint in cancer suppression mechanisms in LLL mammals and DNA repair pathways in ELL mammals [13], and have provided additional comparative genomic evidence for the role of growth factor signaling, such as IGF-1 in model systems and humans [58, 59], in regulating lifespan. Intriguingly, we also found extensively shared networks under parallel constraint in ELL mammals and LLL birds. This implies at least partially convergent strategies for longevity in these two groups of long-lived animals. Compared to mammals, birds as a whole have longer maximum lifespans at a given body size (Figure 1A). From a physiological point of view, it is therefore unsurprising that LLL birds, which are not particularly ELL relative to other birds, but are ELL when considered on the mammalian scale, would share, as a prerequisite for their longevity, a number of pathways under purifying selection with ELL mammals. From an evolutionary perspective, however, it is quite striking for two distinct groups of amniotes, with ancestry separated by 300 million years, to be maintaining the same shared pathways in long-lived members. This implies that there may exist an ancient shared genetic toolkit for longevity, conserved across diverse lineages.

In our positive selection survey, we identified 16 genes, involved in eight distinct hallmarks of aging, with concordant signals of positive selection in LLL mammals and LLL birds at neighboring orthologous amino acid residues. We did so by applying successive rounds of filtering to our initial screen results in order to select the genes with concordant evidence of selection in multiple independent clades. These candidate genes are involved in known aging processes at molecular, cellular, and tissue scales, but have not been extensively studied in the context of aging biology. To move towards implementing evolutionary innovations from other species in humans, it will be important for future studies to validate the longevity-producing potential of these genes in experimental systems and to explore the mechanisms by which they operate.

In recent years, there has been increasing interest in the cross-species regulation of lifespan and the comparative biology of aging. Particularly in mammals, a number of computational and experimental approaches have uncovered mechanisms of longevity in bats, elephants [60], naked mole-rats, and whales [9]. Individual studies have demonstrated a remarkable ability to transpose the pro-longevity effects of a gene from its original species to established model organisms. Naked mole-rat hyaluronan and cGAS, a regulatory enzyme acting on DNA repair, are able to reduce inflammation and increase lifespan in mice and flies, respectively [61, 62]. Bowhead whale CIRBP, an RNA-binding protein, has also been shown to improve homologous recombination repair in human cells [63]. This substantially blurs the line between so-called “private” and “public” mechanisms of longevity, as several studies have demonstrated that pro-longevity innovations in a particular species can often be immediately functional when transposed into another. However, this will not be true in all cases, as some pro-longevity variants may require a particular genetic context that is lacking in the recipient system.

By focusing on convergence between multiple clades and on animals with naturally-occurring longevity, we uncovered more generalizable strategies for longevity [9], allowing us to address fundamental gaps in our understanding of aging. In contrast with more focal traits such as morphological adaptations to flight or marine lifestyles [64], or biochemical adaptation to specific food sources [65], lifespan is not directly encoded *per se* in the genome. It is rather an emergent phenomenon that is determined by interactions between genetics, the environment, and a component of stochastic noise [66]. Nevertheless, species maintain relatively stable characteristic lifespans, and associations between natural lifespan and extrinsic mortality rates indicate that it is actively under the influence of natural selection. Because of its complex nature, however, longer lifespans may convergently evolve through a greater diversity of strategies and mechanisms, resulting in more dispersed signals of selection throughout the genome. By comprehensively surveying the largest set of species available and by doing so in both mammals and birds, we cast a wide net, which has enabled the detection of robust parallels between mammals and birds. Additional studies will allow us to further unravel the intricate web of genetic mechanisms determining longevity.

These results also help to hone our efforts towards translatable mechanisms for improving human healthspans. The hallmarks of aging are a widely recognized framework for understanding aging, and are all significantly associated with aging, but the extent to which each hallmark causes aging in a particular system is variable, and often poorly understood [67]. For example, caloric restriction produces approximately 30–40% lifespan extension in *C. elegans* [68], up to 20% in mice [69], and 10% or less in Rhesus monkeys [70]. Thus, it may be that the lifespan regulatory mechanisms downstream of caloric restriction are already near saturation in longer-lived mammals, and therefore less available for intervention. More generally, different mechanisms are likely to be available for further pro-longevity tuning in humans, who are already long-lived at baseline, than those that work in short-lived laboratory mice. We posit that certain hallmarks of aging are less central to mammalian and avian longevity, while other hallmarks represent lifespan- and healthspan-limiting forms of damage. Continued comparative studies in a variety of species will drive our ability to distinguish which aging mechanisms are most universally active and relevant to human health.

The present study also has limitations that should be noted. While we have been rigorous in our statistical analysis and have selected candidates with maximal support from multiple clades of long-lived animals, experimental evidence is required to establish the longevity effects of any particular pathway or gene. dN/dS-based tests for positive selection are also inherently sensitive to noise, as they are based on relatively rare non-synonymous substitution events [71]. Future work can combine analysis of coding sequence with other genomic elements, such as copy number variation, and integrate data across other high-throughput technologies, such as epigenomics, transcriptomics [72], metabolomics, and proteomics [9] to understand the full biological signaling cascade of long-lived species.

In conclusion, we have presented here the first systematic study of comparative longevity in two diverse vertebrate classes, in which we have uncovered striking parallels between longevity strategies adopted by these amniote lineages separated by 300 million years of evolution. We found a robust network of cell cycle, cancer-suppressing, and mRNA processing pathways under convergent constraint, as well as 16 novel aging-associated genes with concordant signals of positive selection, comprising an extensive set of shared signals of selection between long-lived mammals and birds. These parallel changes hint at deep correspondences in the evolvability of anti-aging mechanisms and in the strategies that may be available to modulate and enhance human longevity and healthspan.

## 4. Methods

### 4.1. Genomic Data

Predictive orthology classifications and summary data on predicted gene inactivation and loss for 503 mammalian and 501 avian genome assemblies were obtained from the TOGA project [32]. As the dataset included multiple genomes for some species, these duplicate genomes were manually inspected and the genomes with newer version numbers and/or higher scaffold N50 were selected for use. Tables S1–S17 show the selected groups of species, with their genome assemblies and phenotypes, used in both purifying and positive selection analyses, while Tables S18–S25 show the groups which did not fit into our clade selections and were therefore used only in the purifying selection analysis.

In order to identify one-to-one orthologous genes, transcripts for each assembly were first matched to the corresponding gene using the “t gene” column in the orthology classifications. Using a table of metadata from Ensembl (https://ensembl.org/, downloaded August 17, 2023), transcript names were also matched to human gene symbols for gene aggregation in pathway-level analysis.

Multiple alignments of coding sequences for 17,434 mammalian and 15,982 avian genes were also downloaded from the same source (https://genome.senckenberg.de/download/TOGA/). Using predictive classifications from TOGA, we selected only genes classified as being intact and one-to-one orthologs with respect to the human (for mammals) or chicken (for birds) reference assemblies.

In addition, in preliminary analyses, we noticed that gaps in the sequence alignments were strongly correlated with a higher chance of a positive selection signal (Figures S1 and S2). This effect persisted over different selections of foreground species, suggesting gap content to be a significant confounding factor, producing false positive signals of positive selection. Therefore, for all analyses, we removed all gapped columns in the sequence alignments to eliminate this source of error.

### 4.2. Phenotype Data

Among species with genome assemblies included in the TOGA multiple alignments, maximum lifespan and mean adult weight data for 295 mammal and 194 bird species were obtained from AnAge (Build 15, released July 3, 2023) [73]. Species with maximum longevity data quality annotated as “low” were excluded from our analysis. In all, our analysis included 237 “acceptable”, 54 “high”, and 4 “questionable” data quality mammalian species and 186 “acceptable”, 3 “high”, and 5 “questionable” data quality bird species.

Least-squares linear regression was used to determine the characteristic relationship between log(weight) and log(lifespan) separately in mammals and birds. Large-bodied longevity (LLL) and exceptional longevity for size (ELL) were defined for each species as the projection of its log(weight) and log(lifespan) onto a unit vector either parallel (LLL) or orthogonal (ELL) to the least-squares line of best fit for the taxonomic class to which that species belongs, either mammals or birds.

### 4.3. Phylogenetic Trees

Organism-level phylogenetic trees for mammals [74] and birds [75] were obtained from the VertLife project (https://vertlife.org/). Species taxonomic names, where different between the VertLife project and the phenotype/genomic data, were manually inspected and harmonized to the VertLife nomenclature.

For initial exploration, phylogeny subsets of our selected 295 mammal and 194 bird species were generated for the distribution of 10,000 trees available in each of four categories: (1) Mammals birth-death node-dated completed trees, (2) Mammals birth-death tip-dated completed trees, (3) Birds Stage2 MayrAll Hackett, and (4) Birds Stage2 MayrAll Ericson. Maximum clade credibility (MCC) mammal and bird trees for our main analyses were also generated as follows: For mammals, the species-level, class-wide “Fossilized birth-death, 5911 species + 76 fossil tips, backbone topology as in Zhou et al. (2013)” tree was obtained from the Vertlife website, and was pruned down to our selected 295 mammalian species. For birds, a composite tree was constructed according to the procedure described in the supplementary information of [75], in which MCC trees for crown clades of birds were grafted onto an MCC backbone tree based on the “Hackett” fossil-calibrated backbone. Briefly, for each crown clade, a sister clade in the backbone tree was identified, outgroups were pruned, and branch lengths were rescaled to match the backbone tree. Three species that were not in the MCC crown clade trees were manually placed using congeners: *Cacatua leadbeateri* using the location of *Cactua sanguinea*, and *Ara militaris* and *Ara rubrogenys* using the locations of *Ara glaucogularis* and *Ara ararauna*.

In order to explore the range of possible phylogenetic histories, we next compared our main mammal and bird trees to two distributions of 10,000 alternate trees using the Robinson-Foulds metric [76]. For mammals, we compared our main tree to 10,000 node-dated trees as well as to 10,000 tip-dated trees, selecting five representative node-dated trees and four representative tip-dated trees for further analysis (Figure S12A). For birds, we compared our main tree to 10,000 “Hackett” backbone trees and 10,000 “Ericson” backbone trees, selecting four “Hackett” backbone trees and five representative “Ericson” backbone trees for further analysis (Figure S12B).

All these alternate trees were selected randomly, with manual inspection to ensure a representative range of Robinson-Foulds distances, and to exclude any trees with topologies identical to already-selected ones. The higher-level topology of the clades was identical to the main trees for most alternate trees selected (Figure S13), but mN3424 and bE0570 did have different clade relationships (Perissodactyla forming a sister clade with Carnivora in mN3424 and Aequornithes forming a sister clade with Charadriiformes in bE0570). Differences between each alternate tree and the main tree used are shown in Figures S14–S31.

### 4.4. Purifying Selection Tests

To assess for purifying selection, the RERConverge package [17] was used to correlate gene-wise rates of sequence evolution with inferred rates of phenotypic (maximum lifespan) change. Pearson linear correlations were used to assess the strength of purifying selection in long-lived lineages for individual genes. As required by the software package, our selected sets of 295 mammal and 194 bird species were reduced to only include species with sufficient one-to-one orthology (30 genes shared across the entire set of species), as predicted by TOGA, to generate a “master gene tree”. By iteratively removing the species with fewest one-to-one orthologs shared with all other remaining species, we selected a final set of 195 mammal and 184 bird species to use in our purifying selection analyses.

For each gene, RERConverge was run on species in which that gene was predicted to be a one-to-one ortholog with respect to the reference (human for mammals and chicken for birds) genome for that class.

To assess purifying selection at the pathway level, gene set assignments compiled from various sources were obtained from MSigDB ( [77], C2: curated gene sets, downloaded March 28, 2024). Pathway enrichment statistics were computed by RERConverge, as described in [13]. For each gene set, the p-values from the Pearson linear correlation analysis for individual genes in the gene set were compared to the same measure for genes included in any pathway annotation using a Wilcoxon Rank-Sum test.

### 4.5. Positive Selection Tests

Both in order to highlight convergent mechanisms of longevity across different groups of birds and mammals and for computational tractability, we selected several clades of mammals and birds for our positive selection analysis. We started by grouping our 295 mammal and 194 bird species by traditional taxonomic orders, then made several modifications to enforce monophyly and to keep clades at a reasonable number of species. In particular, we sometimes grouped multiple orders into one clade (e.g. Aequornithes), split a large clade into lower-level taxa (e.g. Passeriformes into Passida, Muscicapida, Sylviida, and Corvides), or selected only a subgroup of species from a paraphyletic order (e.g. Ruminantia from Artiodactyla). We also required clades to have a minimum of 8 species.

Within each clade, the top quartile of species for “ELL” or “LLL” longevity were designated as foreground species for each respective analysis. Branch-site tests using codeML from the PAML package [15] were then performed on genes that contained at least 8 species, including at least 2 foreground species, that were predicted by TOGA to have that gene as a one-to-one ortholog and were present in the multiple alignment files.

CodeML was run using the following parameters: cleandata = 1 (removes all gaps and ambiguous sequences), seqtype = 1 (specifying codon rather than nucleotide sequence input), aaDist = 0 (equal amino acid distances), CodonFreq = 1 (equilibrium codon frequencies in codon substitution model are calculated from the average nucleotide frequencies), model = 2 (branch model: multiple different omega ratios for each group of branches), clock = 0 (no clock and rates are free to among branches), and NSsites = 2 (M2a model where selection is allowed). Likelihoods of null models with fixed omega = 1 were then compared to alternative models where omega was allowed to possibly be *>*1 (positive selection).

Within each clade and phenotype surveyed, p-values were adjusted using the Benjamini-Hochberg procedure as implemented in scipy [78]. Significant genes with *FDR <* 0.05 at the clade and phenotype level were then selected for further analysis.

To integrate our results across several avian and mammalian clades, we defined a “total support” metric for each gene (Equation 4.1). In this metric, which is a modified version of a harmonic p-value, the negative logarithms of significant FDRs from each clade within a vertebrate class are summed, after which the mammalian and avian sums are multiplied together. The resultant number gives a measure of the total statistical support for positive selection acting on a gene across all clades, while emphasizing convergence by requiring support from at least one mammalian and one avian clade.

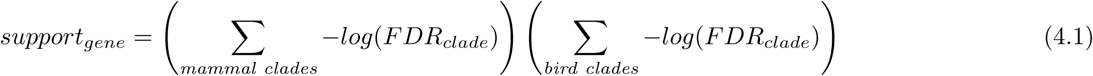

### 4.6. Additional Alternate Tree Analyses

In order to capture the effects of uncertainty in our species trees, we repeated our purifying and positive selection analyses in each of the 9 alternate mammal and 9 alternate bird trees whenever the species topology of an alternate tree differed from our main tree.

For our purifying selection results, all alternate trees were expressly selected to have distinct topologies, so control analyses were run for all 18 alternate trees. To take into account our phylogenetic uncertainty and to summarize the most consistently constrained pathways, we examined the pathways that were constrained in both the main tree and all 9 alternate trees. The results were remarkably consistent, with nearly all connected groups of pathways from the main analysis persisting in all alternate trees (Figure S32).

Since our positive selection results were derived from analyses of variation in smaller clades within mammals or birds, we limited our alternate tree positive selection tests to those in which the topology of a particular smaller clade differed from the main tree topology. We then examined the fraction of genes with significant signals of positive selection (*FDR <* 0.05) from our main analysis that were also significant in each alternate topology. As the mammalian phylogeny is more well-established, only Chiroptera and Rodentia had any topological variation, and only in node-dated trees. By contrast, six of the nine bird clades had topology variations. Overall, the results were also quite concordant across different topologies, with more than 80% of genes generally being concordant between our main analysis and alternate tree analyses (Figure S33).

### 4.7. Additional Statistical Analysis

As a background null distribution of p-values, we performed permulations (permutations informed by phylogenetic simulations [33]) of our phenotypes for both the purifying and positive selection analyses. For the purifying selection tests, 100 rounds of permulation resampling of our foreground species were applied to all 16,785 mammal and 14,489 bird genes tested, resulting in 1,678,500 mammal and 1,448,900 bird p-values for constituting the null distribution for each phenotype (LLL or ELL). For the positive selection analyses, due to heavier computational requirements, each gene and clade was only resampled once. To still obtain a representative sample of p-values, a different permulation was used for each clade-gene pair. Depending on the analysis, some genes in some clades were omitted due to insufficient aligned foreground or background sequences, resulting in a total of 105,730 background values for the mammal LLL analysis, 105,684 for mammal ELL, 102,390 for bird LLL, and 101,768 for mammal ELL.

In the positive selection results, the overwhelming majority of p-values are close to 1 (Figure 3E–H). Therefore, to test the enrichment of the statistically significant lowest p-values, we restricted our testing to the lowest 10% of p-values. This corresponded to a range of values roughly from 0 to 0.5. We then applied the non-parametric Mann-Whitney-U test to the lowest 10% of p-values in each analysis to determine whether the positive selection results using actual phenotypes were enriched in low p-values when compared to a background distribution of permulated p-values.

To group pathways under significant constraint into broader networks representing biological processes, for each analysis we created a graph with each significant pathway as a node, then compared the gene sets among all significant pathways, drawing an edge between two nodes when more than 70% of genes were shared (considering the pathway/node with fewer total genes). We then manually inspected the constituent nodes/pathways in each connected network and selected labels for the biological processes (Tables S26, S27, S28, S29, and S30). There were also some unconnected significant pathways, which met our significance threshold of *FDR <* 0.05 but did not have any edges drawn to other significant pathways. These tended to be more empirically derived and less well-studied (Figures S34, S35, S36 and S37).

To validate candidate genes from our positive selection tests, we performed 100 rounds of permulation resampling of our foreground species. In each round, we used the species phylogeny to randomly simulate phenotype evolution using the “simulatevec” command in RERConverge. This produces distributions of phenotypes that retain phylogenetic relationships among species. We then selected, as in the regular positive selection tests, the species in the highest quartile of simulated phenotypes as foreground species for branch-site tests in PAML using codeML.

### 4.8. Visualizations

The MAFTT version 7 webserver (https://mafft.cbrc.jp/alignment/server/index.html) was used to align the reference bird (*Gallus gallus*) protein sequence to our mammalian multiple alignments for 174 candidate genes with statistical support (*FDR <* 0.05) for positive selection in LLL species from at least one mammalian and one avian clade [79]. Amino acid chain numbers from the supporting clades were then matched with custom python code, and sites within 20 amino acids of each other were selected as genes with possibly convergent selection at orthologous active sites.

Icons of ancestral mammals and birds for Figure 1 were created from illustrations of *Morganucodon* and *Archaeopteryx* from files on Wikimedia Commons uploaded by FunkMonk (Michael B. H.) and UnexpectedDinoLesson, respectively, by defining a color threshold to create a silhouette, and then re-coloring that binary image. Clade representative icons (for Figure 6) were obtained from PhyloPic (www.phylopic.org). A full list of artists/uploaders is provided (Table S31).

## 5. Acknowledgements

We would like to thank members of the Kronforst lab for comments and discussion, the University of Chicago Center for Research Informatics (CRI) for high-performance computing resources, and Dr. Amy Z. Xu for assistance with illustration and comments on the manuscript.

## 5.1. Author Contributions

W.B.Z. and M.R.K. designed the research project; W.B.Z. performed the computational experiments and analyzed the data; W.B.Z. and M.R.K. wrote the manuscript.

## 5.4. Data Sharing Plans

As described in the methods, this project used publicly available data: orthology predictions and multiple alignments from the TOGA project, phylogenetic trees from VertLife, and phenotype data from AnAge. Intermediate PAML and RERConverge result files, processed multiple alignments, and python projects used for analysis will be shared on Dryad prior to final publication of the manuscript.

## 7. Supplementary Information

**Table S1:**
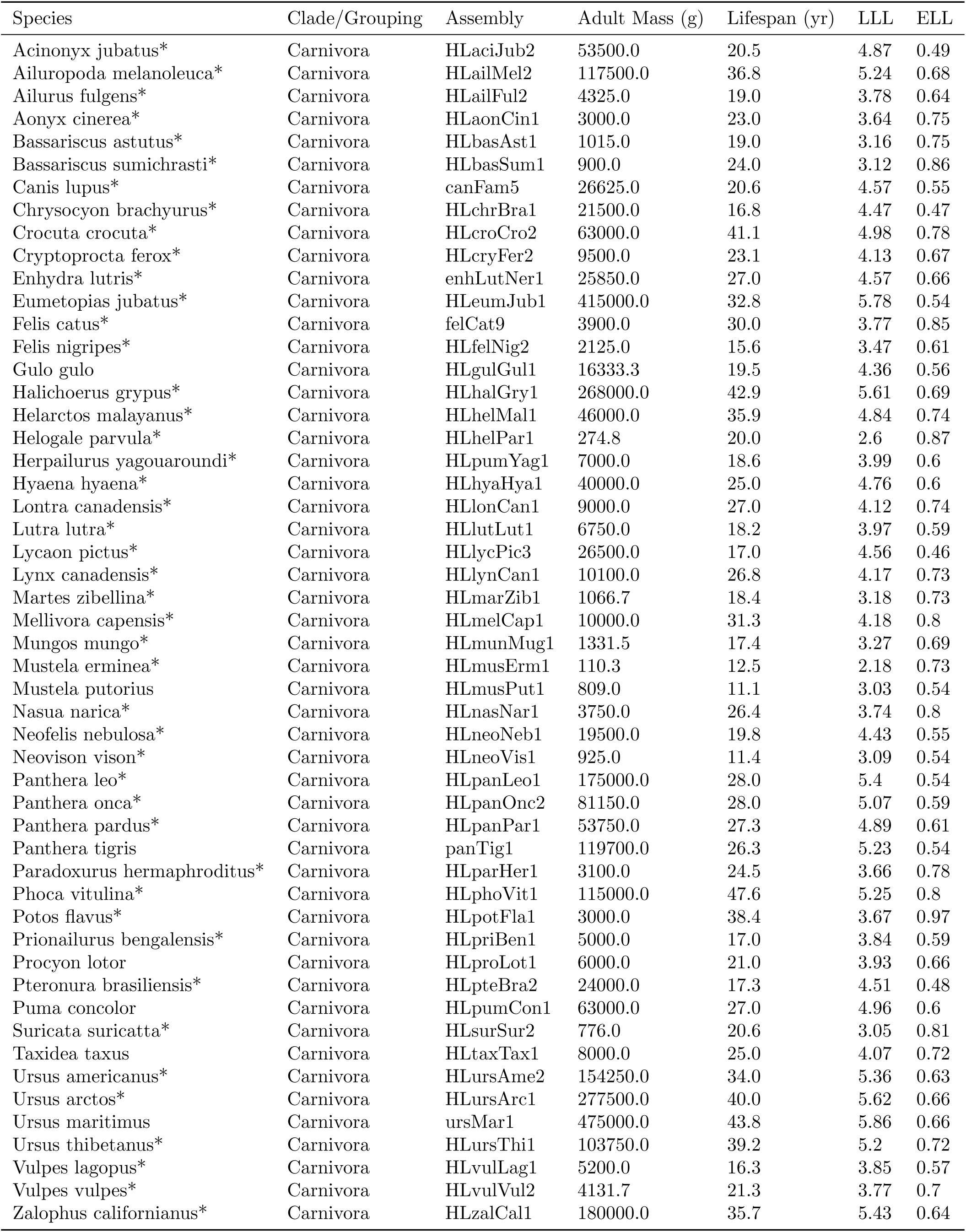
Phenotypes and genome assemblies used for Carnivora. *\* indicates species used in main purifying selection analyses*.

**Table S2:**
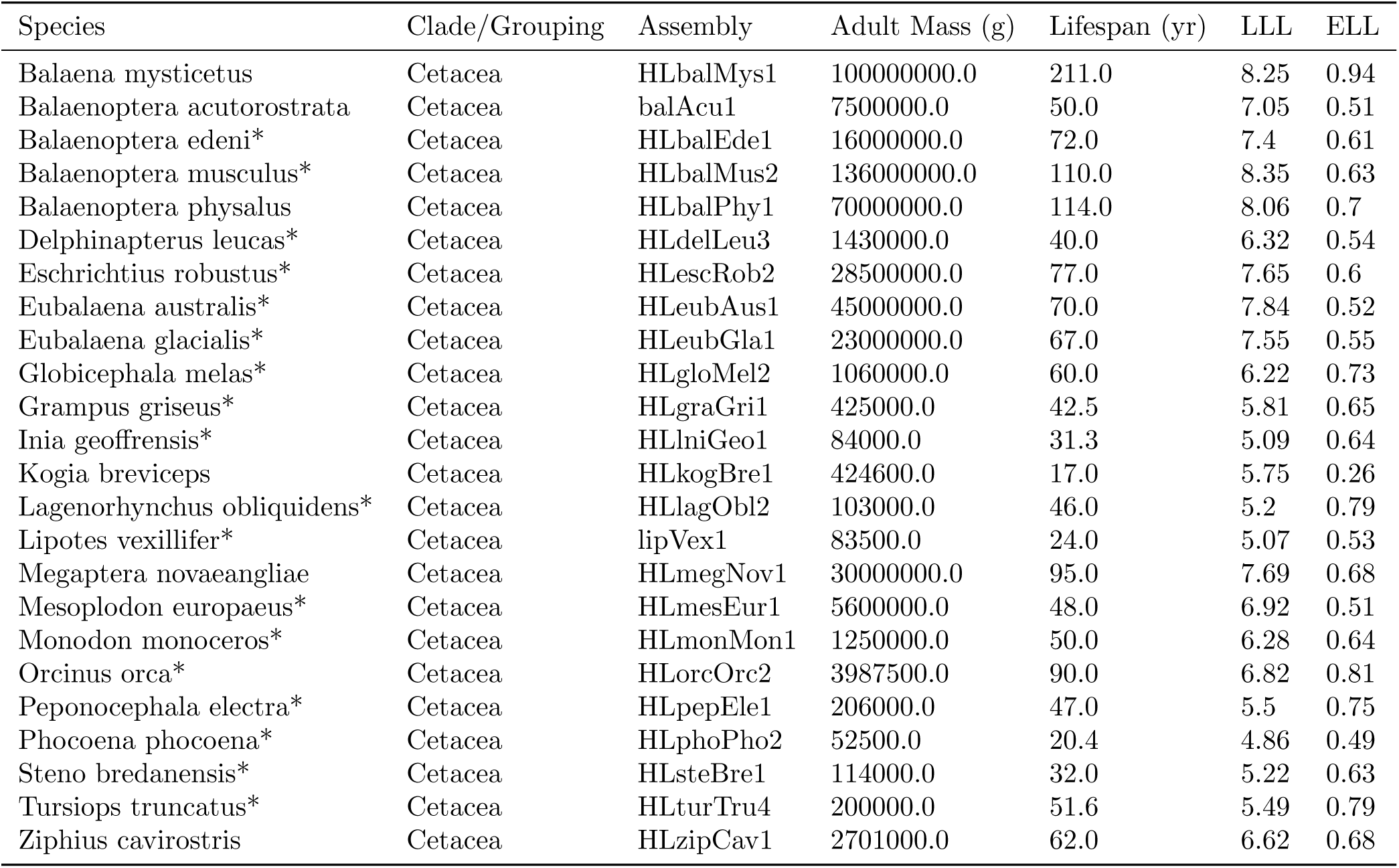
Phenotypes and genome assemblies used for Cetacea. *\* indicates species used in main purifying selection analyses.*

| Species | Clade/Grouping | Assembly | Adult Mass (g) | Lifespan (yr) | LLL | ELL |
| --- | --- | --- | --- | --- | --- | --- |
| Balaena mysticetus | Cetacea | HLbalMys1 | 100000000.0 | 211.0 | 8.25 | 0.94 |
| Balaenoptera acutorostrata | Cetacea | balAcu1 | 7500000.0 | 50.0 | 7.05 | 0.51 |
| Balaenoptera edeni* | Cetacea | HLbalEde1 | 16000000.0 | 72.0 | 7.4 | 0.61 |
| Balaenoptera musculus* | Cetacea | HLbalMus2 | 136000000.0 | 110.0 | 8.35 | 0.63 |
| Balaenoptera physalus | Cetacea | HLbalPhy1 | 70000000.0 | 114.0 | 8.06 | 0.7 |
| Delphinapterus leucas* | Cetacea | HLdelLeu3 | 1430000.0 | 40.0 | 6.32 | 0.54 |
| Eschrichtius robustus* | Cetacea | HLescRob2 | 28500000.0 | 77.0 | 7.65 | 0.6 |
| Eubalaena australis* | Cetacea | HLeubAus1 | 45000000.0 | 70.0 | 7.84 | 0.52 |
| Eubalaena glacialis* | Cetacea | HLeubGla1 | 23000000.0 | 67.0 | 7.55 | 0.55 |
| Globicephala melas* | Cetacea | HLgloMel2 | 1060000.0 | 60.0 | 6.22 | 0.73 |
| Grampus griseus* | Cetacea | HLgraGri1 | 425000.0 | 42.5 | 5.81 | 0.65 |
| Inia geoffrensis* | Cetacea | HLlniGeo1 | 84000.0 | 31.3 | 5.09 | 0.64 |
| Kogia breviceps | Cetacea | HLkogBre1 | 424600.0 | 17.0 | 5.75 | 0.26 |
| Lagenorhynchus obliquidens* | Cetacea | HLlagObl2 | 103000.0 | 46.0 | 5.2 | 0.79 |
| Lipotes vexillifer* | Cetacea | lipVex1 | 83500.0 | 24.0 | 5.07 | 0.53 |
| Megaptera novaeangliae | Cetacea | HLmegNov1 | 30000000.0 | 95.0 | 7.69 | 0.68 |
| Mesoplodon europaeus* | Cetacea | HLmesEur1 | 5600000.0 | 48.0 | 6.92 | 0.51 |
| Monodon monoceros* | Cetacea | HLmonMon1 | 1250000.0 | 50.0 | 6.28 | 0.64 |
| Orcinus orca* | Cetacea | HLorcOrc2 | 3987500.0 | 90.0 | 6.82 | 0.81 |
| Peponocephala electra* | Cetacea | HLpepEle1 | 206000.0 | 47.0 | 5.5 | 0.75 |
| Phocoena phocoena* | Cetacea | HLphoPho2 | 52500.0 | 20.4 | 4.86 | 0.49 |
| Steno bredanensis* | Cetacea | HLsteBre1 | 114000.0 | 32.0 | 5.22 | 0.63 |
| Tursiops truncatus* | Cetacea | HLturTru4 | 200000.0 | 51.6 | 5.49 | 0.79 |
| Ziphius cavirostris | Cetacea | HLzipCav1 | 2701000.0 | 62.0 | 6.62 | 0.68 |

**Table S3:** Phenotypes and genome assemblies used for Chiroptera. *\* indicates species used in main purifying selection analyses*.

| Species | Clade/Grouping | Assembly | Adult Mass (g) | Lifespan (yr) | LLL | ELL |
| --- | --- | --- | --- | --- | --- | --- |
| Antrozous pallidus | Chiroptera | HLantPal1 | 20.8 | 14.8 | 1.48 | 0.93 |
| Artibeus jamaicensis* | Chiroptera | HLartJam2 | 42.2 | 19.2 | 1.8 | 0.99 |
| Carollia perspicillata | Chiroptera | HLcarPer3 | 15.0 | 17.0 | 1.34 | 1.01 |
| Cynopterus brachyotis | Chiroptera | HLcynBra1 | 45.0 | 10.1 | 1.78 | 0.71 |
| Eidolon helvum | Chiroptera | HLeidHel2 | 306.2 | 21.8 | 2.66 | 0.9 |
| Eptesicus fuscus* | Chiroptera | HLeptFus2 | 23.0 | 19.0 | 1.53 | 1.03 |
| Macrotus californicus | Chiroptera | HLmacCal1 | 11.6 | 17.0 | 1.23 | 1.03 |
| Megaderma lyra* | Chiroptera | HLmegLyr2 | 38.6 | 14.0 | 1.74 | 0.86 |
| Miniopterus schreibersii* | Chiroptera | HLminSch1 | 13.0 | 22.0 | 1.3 | 1.13 |
| Molossus molossus* | Chiroptera | HLmolMol2 | 14.0 | 5.6 | 1.24 | 0.54 |
| Myotis brandtii | Chiroptera | myoBra1 | 7.0 | 41.0 | 1.07 | 1.45 |
| Myotis lucifugus | Chiroptera | myoLuc2 | 10.0 | 34.0 | 1.21 | 1.34 |
| Myotis myotis* | Chiroptera | HLmyoMyo6 | 28.6 | 37.1 | 1.67 | 1.3 |
| Noctilio leporinus* | Chiroptera | HLnocLep1 | 50.0 | 11.8 | 1.84 | 0.77 |
| Nycticeius humeralis | Chiroptera | HLnycHum2 | 9.8 | 6.0 | 1.09 | 0.6 |
| Phyllostomus discolor* | Chiroptera | HLphyDis3 | 43.4 | 9.0 | 1.76 | 0.66 |
| Pipistrellus kuhlii* | Chiroptera | HLpipKuh2 | 6.0 | 8.0 | 0.9 | 0.76 |
| Pipistrellus pipistrellus* | Chiroptera | HLpipPip2 | 5.0 | 16.6 | 0.87 | 1.08 |
| Pteropus alecto | Chiroptera | pteAle1 | 672.1 | 20.3 | 2.99 | 0.81 |
| Pteropus giganteus* | Chiroptera | HLpteGig1 | 1175.0 | 44.0 | 3.28 | 1.1 |
| Pteropus pselaphon* | Chiroptera | HLptePse1 | 321.5 | 16.7 | 2.66 | 0.78 |
| Pteropus vampyrus | Chiroptera | HLpteVam2 | 872.0 | 20.9 | 3.1 | 0.8 |
| Rhinolophus ferrumequinum* | Chiroptera | HLrhiFer5 | 22.9 | 30.5 | 1.56 | 1.23 |
| Rousettus aegyptiacus* | Chiroptera | HLrouAeg4 | 125.0 | 22.9 | 2.27 | 0.98 |
| Rousettus leschenaultii* | Chiroptera | HLrouLes1 | 108.2 | 14.0 | 2.18 | 0.78 |
| Tadarida brasiliensis | Chiroptera | HLtadBra1 | 12.5 | 12.0 | 1.24 | 0.88 |

**Table S4:** Phenotypes and genome assemblies used for Perissodactyla. *\* indicates species used in main purifying selection analyses*.

| Species | Clade/Grouping | Assembly | Adult Mass (g) | Lifespan (yr) | LLL | ELL |
| --- | --- | --- | --- | --- | --- | --- |
| Ceratotherium simum* | Perissodactyla | cerSim1 | 2175000.0 | 45.0 | 6.51 | 0.56 |
| Dicerorhinus sumatrensis* | Perissodactyla | HLdicSum1 | 1260000.0 | 32.8 | 6.26 | 0.46 |
| Diceros bicornis* | Perissodactyla | HLdicBic1 | 1100000.0 | 57.0 | 6.23 | 0.71 |
| Equus africanus* | Perissodactyla | HLequAsiAsi2 | 164998.0 | 47.0 | 5.41 | 0.76 |
| Equus caballus* | Perissodactyla | equCab3 | 300000.0 | 57.0 | 5.68 | 0.8 |
| Equus zebra* | Perissodactyla | HLequZeb1 | 296000.0 | 33.2 | 5.64 | 0.57 |
| Rhinoceros unicornis* | Perissodactyla | HLrhiUni1 | 1602330.0 | 43.5 | 6.38 | 0.56 |
| Tapirus terrestris* | Perissodactyla | HLtapTer2 | 250000.0 | 39.6 | 5.57 | 0.66 |

**Table S5:**
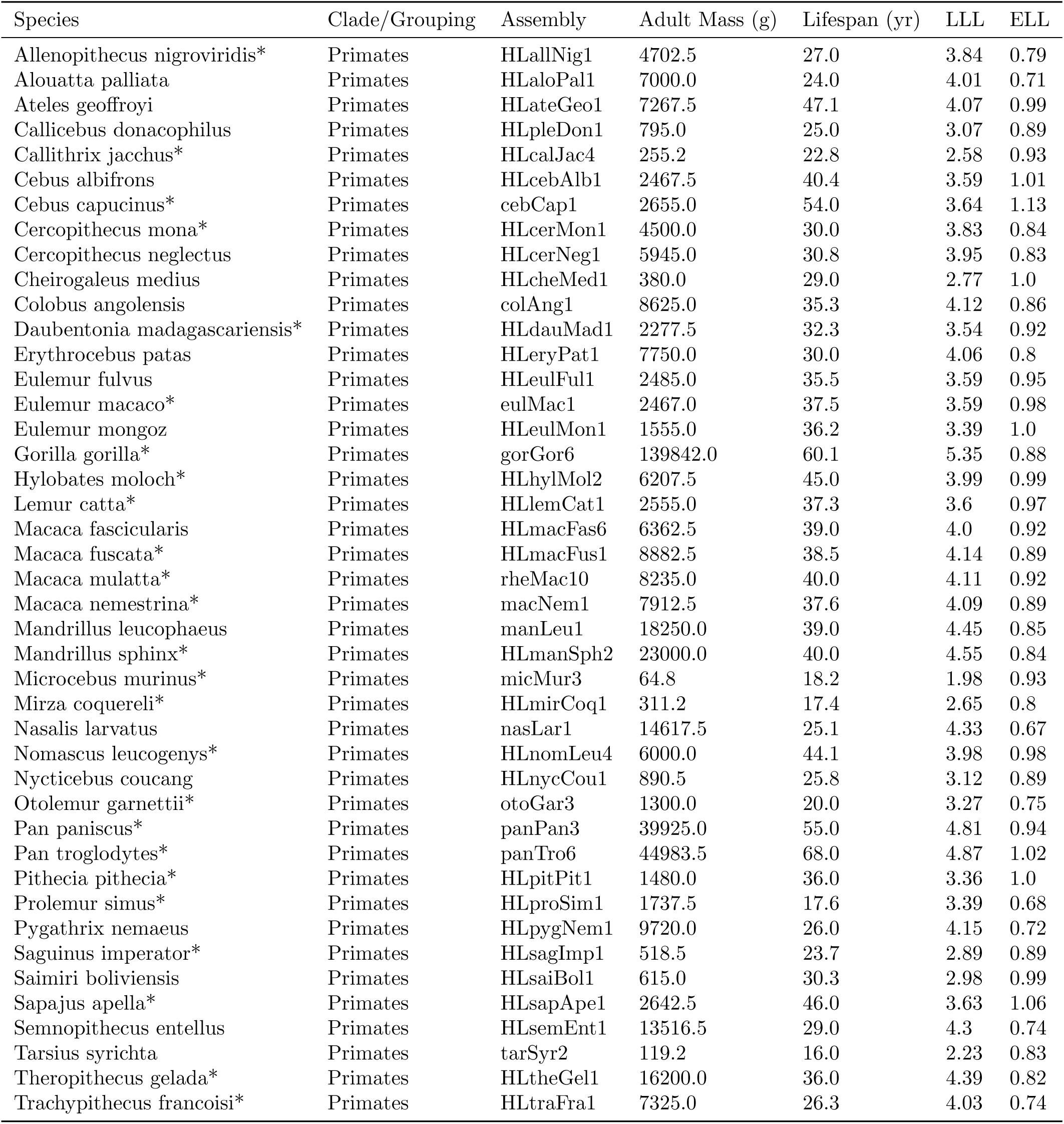
Phenotypes and genome assemblies used for Primates. *\* indicates species used in main purifying selection analyses*.

**Table S6:**
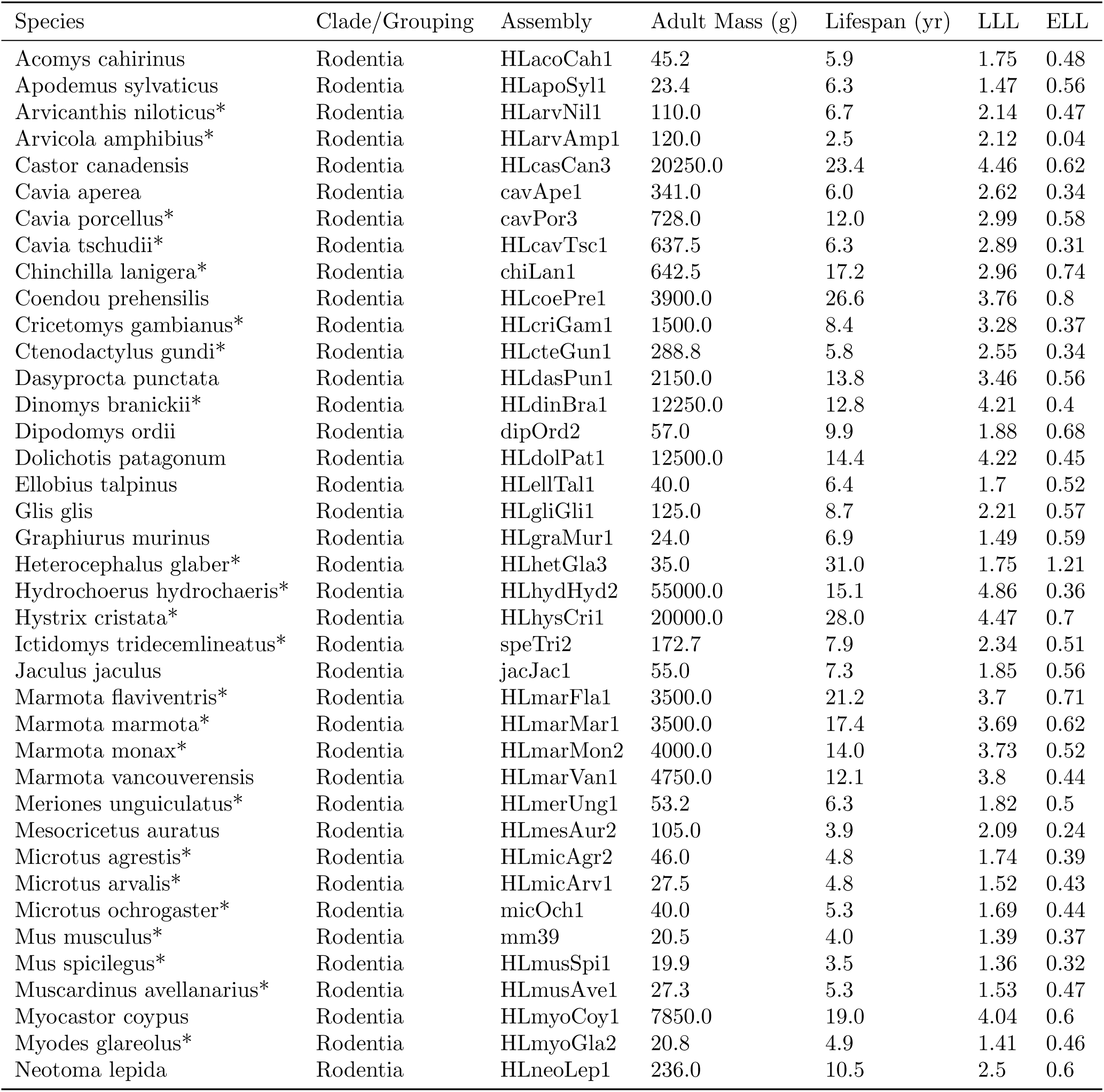
Phenotypes and genome assemblies used for Rodentia (part 1). *\* indicates species used in main purifying selection analyses*.

**Table S7:** Phenotypes and genome assemblies used for Rodentia (part 2). *\* indicates species used in main purifying selection analyses*.

| Species | Clade/Grouping | Assembly | Adult Mass (g) | Lifespan (yr) | LLL | ELL |
| --- | --- | --- | --- | --- | --- | --- |
| Octodon degus* | Rodentia | octDeg1 | 235.0 | 14.0 | 2.51 | 0.73 |
| Onychomys torridus* | Rodentia | HLonyTor1 | 22.0 | 4.6 | 1.43 | 0.43 |
| Pachyuromys duprasi* | Rodentia | HLpacDup1 | 40.0 | 8.3 | 1.72 | 0.63 |
| Pedetes capensis* | Rodentia | HLpedCap1 | 3500.0 | 20.0 | 3.7 | 0.68 |
| Perognathus longimembris | Rodentia | HLperLonPac1 | 8.0 | 8.3 | 1.03 | 0.75 |
| Peromyscus californicus* | Rodentia | HLperCal2 | 42.0 | 5.5 | 1.71 | 0.45 |
| Peromyscus crinitus | Rodentia | HLperCri1 | 16.5 | 7.7 | 1.33 | 0.67 |
| Peromyscus eremicus | Rodentia | HLperEre1 | 25.0 | 7.4 | 1.51 | 0.62 |
| Peromyscus leucopus* | Rodentia | HLperLeu1 | 23.0 | 7.9 | 1.48 | 0.65 |
| Peromyscus maniculatus* | Rodentia | HLperManBai2 | 20.5 | 8.3 | 1.43 | 0.68 |
| Peromyscus polionotus* | Rodentia | HLperPol1 | 14.0 | 5.5 | 1.24 | 0.54 |
| Psammomys obesus* | Rodentia | HLpsaObe1 | 212.0 | 3.2 | 2.38 | 0.1 |
| Rattus norvegicus* | Rodentia | HLratNor7 | 300.0 | 3.8 | 2.54 | 0.15 |
| Rattus rattus* | Rodentia | HLratRat7 | 200.0 | 4.2 | 2.37 | 0.22 |
| Rhombomys opimus* | Rodentia | HLrhoOpi1 | 285.0 | 4.5 | 2.52 | 0.23 |
| Sciurus carolinensis* | Rodentia | HLsciCar1 | 533.0 | 23.6 | 2.9 | 0.89 |
| Sciurus vulgaris* | Rodentia | HLsciVul1 | 600.0 | 14.8 | 2.92 | 0.68 |
| Sigmodon hispidus* | Rodentia | HLsigHis1 | 185.0 | 5.2 | 2.35 | 0.32 |
| Thryonomys swinderianus | Rodentia | HLthrSwi1 | 6600.0 | 5.4 | 3.89 | 0.07 |
| Xerus inauris | Rodentia | HLxerIna1 | 588.0 | 11.5 | 2.9 | 0.58 |
| Zapus hudsonius | Rodentia | HLzapHud1 | 18.0 | 5.6 | 1.35 | 0.52 |

**Table S8:**
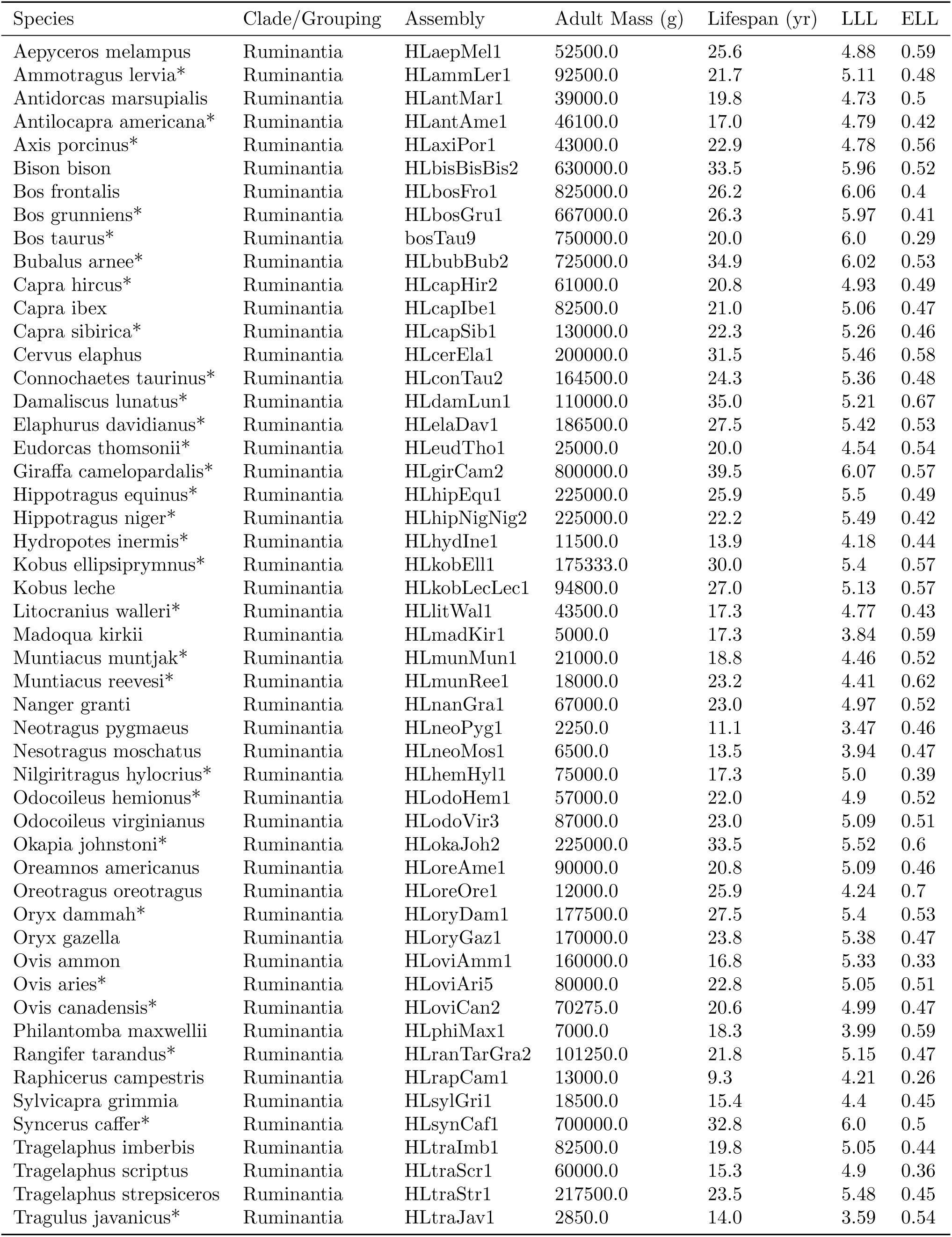
Phenotypes and genome assemblies used for Ruminantia. *\* indicates species used in main purifying selection analyses*.

**Table S9:**
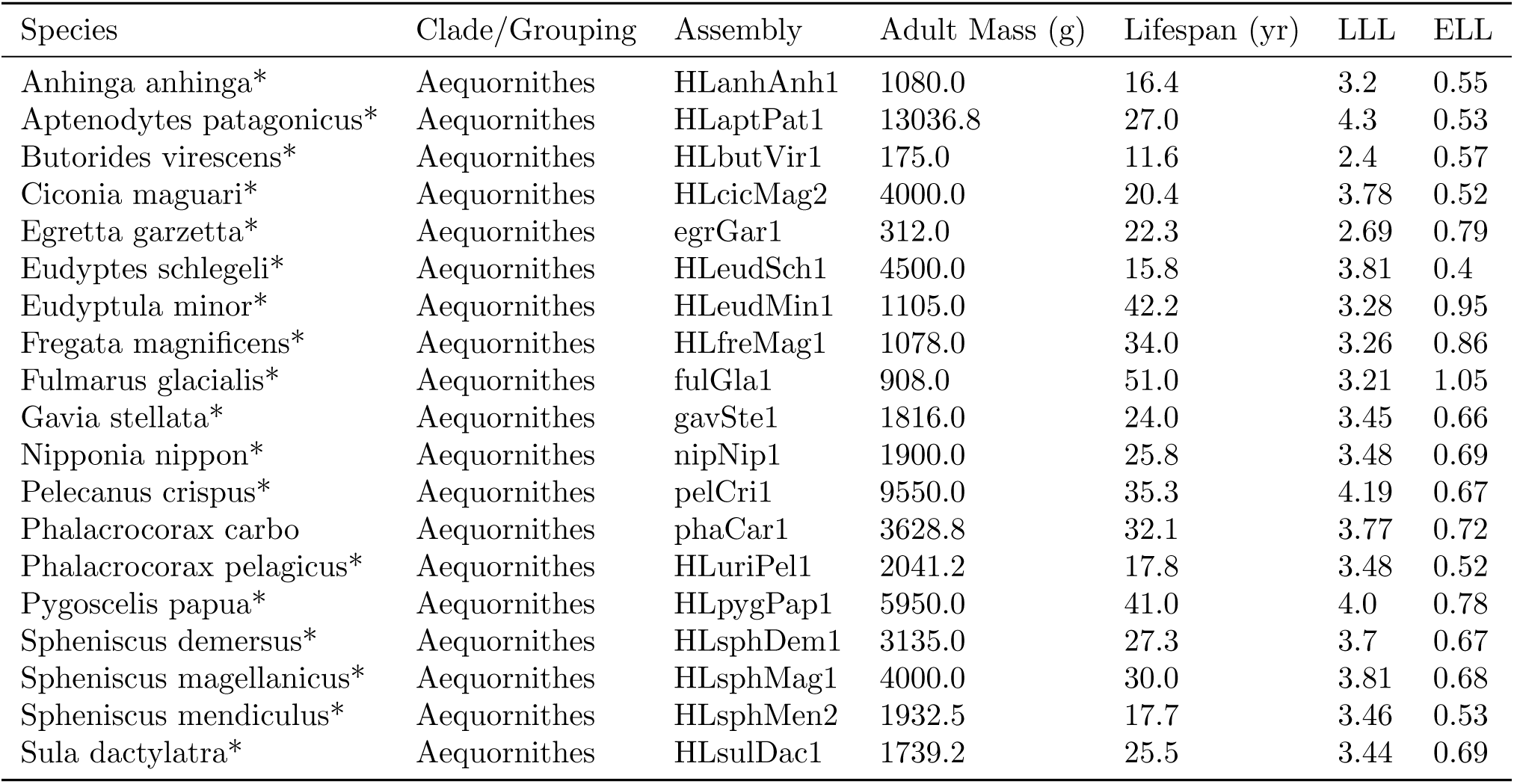
Phenotypes and genome assemblies used for Aequornithes. *\* indicates species used in main purifying selection analyses*.

**Table S10:** Phenotypes and genome assemblies used for Anseriformes. *\* indicates species used in main purifying selection analyses*.

| Species | Clade/Grouping | Assembly | Adult Mass (g) | Lifespan (yr) | LLL | ELL |
| --- | --- | --- | --- | --- | --- | --- |
| Anas platyrhynchos* | Anseriformes | HLanaPla1 | 1048.1 | 29.1 | 3.23 | 0.79 |
| Anser brachyrhynchus* | Anseriformes | HLansBra1 | 2645.0 | 40.9 | 3.65 | 0.85 |
| Anseranas semipalmata* | Anseriformes | HLansSem1 | 2071.0 | 24.8 | 3.51 | 0.66 |
| Aythya fuligula* | Anseriformes | HLaytFul1 | 701.5 | 45.2 | 3.09 | 1.02 |
| Branta canadensis* | Anseriformes | HLbraCan1 | 3200.0 | 42.0 | 3.74 | 0.85 |
| Cygnus atratus* | Anseriformes | HLcygAtr1 | 5650.0 | 24.7 | 3.94 | 0.57 |
| Cygnus buccinator* | Anseriformes | HLcygBuc1 | 13000.0 | 32.5 | 4.32 | 0.61 |
| Cygnus olor* | Anseriformes | HLcygOlo1 | 8300.0 | 40.0 | 4.14 | 0.74 |
| Oxyura jamaicensis* | Anseriformes | HLoxyJam1 | 542.0 | 20.0 | 2.92 | 0.7 |

**Table S11:**
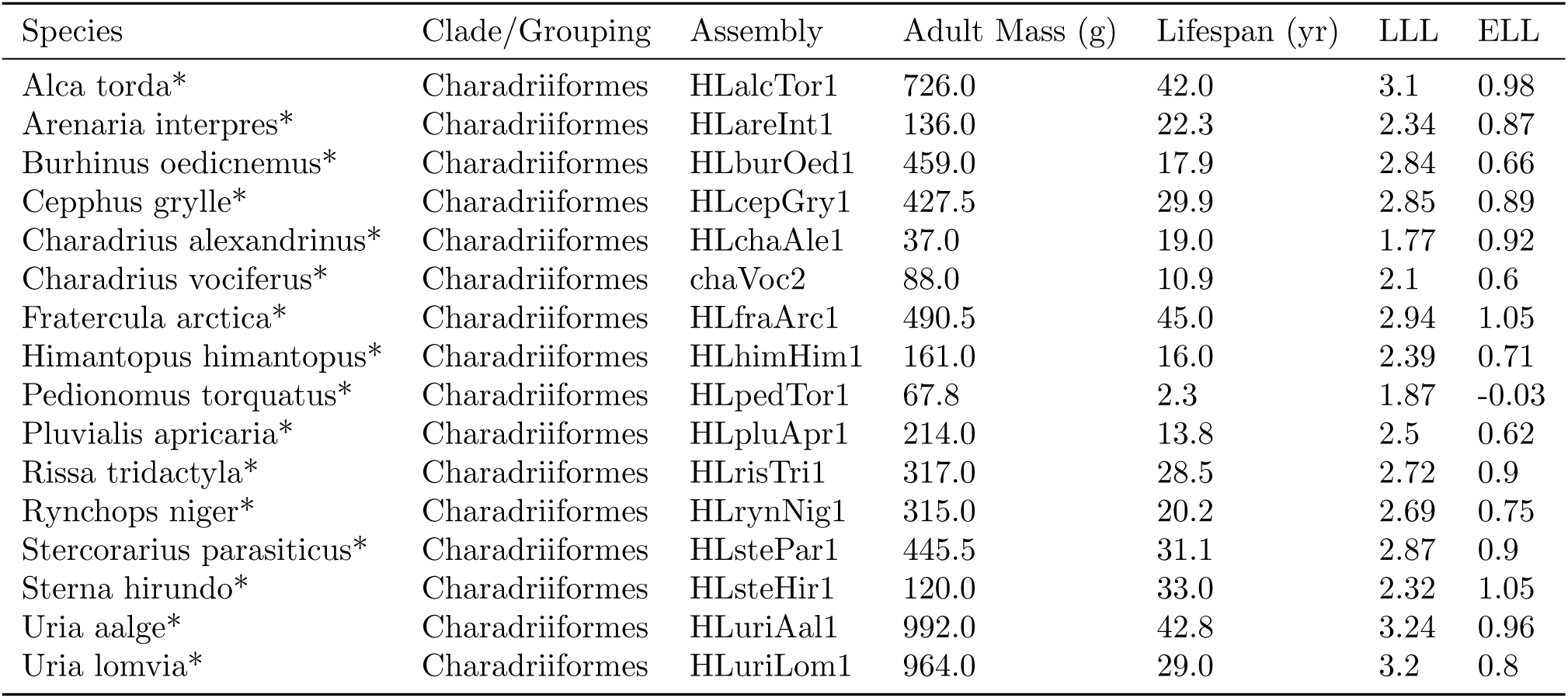
Phenotypes and genome assemblies used for Charadriiformes. *\* indicates species used in main purifying selection analyses*.

**Table S12:**
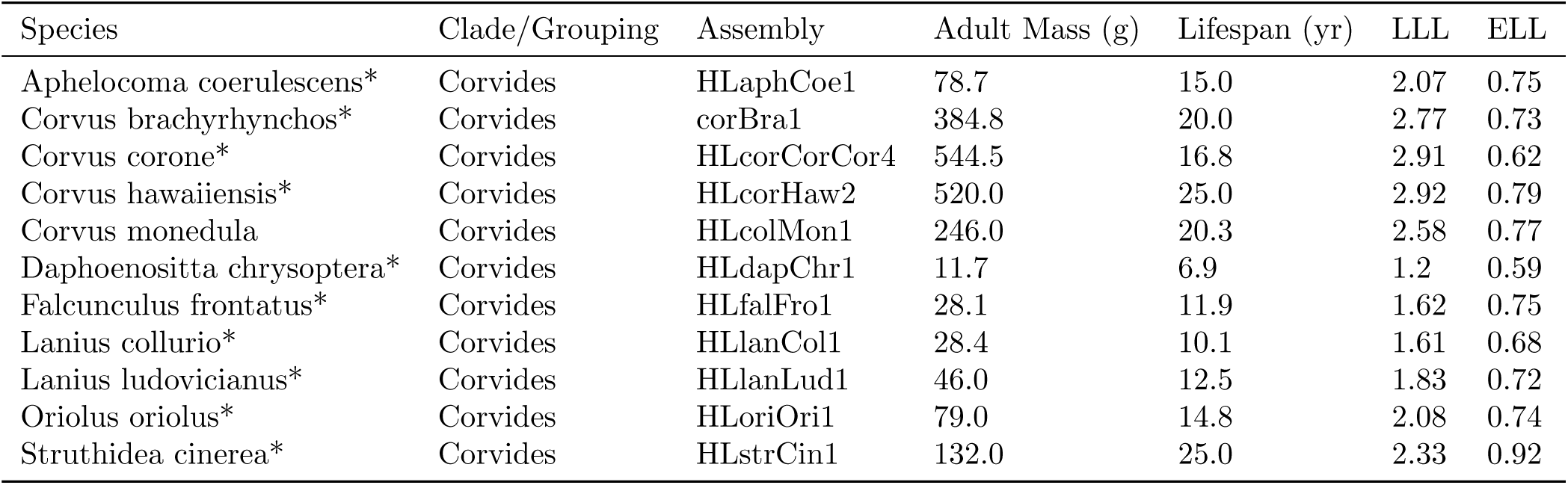
Phenotypes and genome assemblies used for Corvides. *\* indicates species used in main purifying selection analyses*.

**Table S13:** Phenotypes and genome assemblies used for Galliformes. *\* indicates species used in main purifying selection analyses*.

| Species | Clade/Grouping | Assembly | Adult Mass (g) | Lifespan (yr) | LLL | ELL |
| --- | --- | --- | --- | --- | --- | --- |
| Alectoris rufa* | Galliformes | HLalcRuf1 | 528.0 | 6.2 | 2.82 | 0.2 |
| Centrocercus urophasianus* | Galliformes | HLcenUro1 | 1929.5 | 7.0 | 3.38 | 0.13 |
| Chrysolophus pictus* | Galliformes | HLchrPic1 | 625.0 | 13.4 | 2.95 | 0.51 |
| Colinus virginianus* | Galliformes | HLcolVir2 | 194.0 | 6.4 | 2.39 | 0.31 |
| Coturnix japonica* | Galliformes | HLcotJap2 | 115.0 | 6.0 | 2.17 | 0.33 |
| Lagopus leucura* | Galliformes | HLlagLeu1 | 326.0 | 15.0 | 2.68 | 0.62 |
| Meleagris gallopavo* | Galliformes | HLmelGal6 | 6050.0 | 13.0 | 3.92 | 0.29 |
| Pavo cristatus | Galliformes | HLpavCri1 | 4200.0 | 23.2 | 3.81 | 0.57 |
| Phasianus colchicus* | Galliformes | HLphaCol2 | 1094.8 | 27.0 | 3.24 | 0.76 |
| Tympanuchus cupido* | Galliformes | HLtymCup1 | 908.0 | 5.0 | 3.04 | 0.06 |

**Table S14:** Phenotypes and genome assemblies used for Muscicapida. *\* indicates species used in main purifying selection analyses*.

| Species | Clade/Grouping | Assembly | Adult Mass (g) | Lifespan (yr) | LLL | ELL |
| --- | --- | --- | --- | --- | --- | --- |
| Bombycilla garrulus* | Muscicapida | HLbomGar1 | 54.5 | 13.5 | 1.91 | 0.74 |
| Catharus bicknelli* | Muscicapida | HLcatBic1 | 27.5 | 11.9 | 1.61 | 0.75 |
| Catharus fuscescens* | Muscicapida | HLcatFus1 | 28.8 | 10.1 | 1.61 | 0.67 |
| Catharus minimus* | Muscicapida | HLcatMin1 | 24.8 | 7.3 | 1.53 | 0.55 |
| Catharus ustulatus* | Muscicapida | HLcatUst1 | 27.9 | 11.0 | 1.61 | 0.71 |
| Certhia familiaris* | Muscicapida | HLcerFam1 | 9.1 | 8.2 | 1.11 | 0.69 |
| Cinclus mexicanus* | Muscicapida | HLcinMex1 | 50.2 | 8.1 | 1.83 | 0.53 |
| Erithacus rubecula* | Muscicapida | HLeriRub1 | 17.3 | 19.3 | 1.45 | 1.0 |
| Ficedula albicollis* | Muscicapida | ficAlb2 | 12.7 | 9.8 | 1.26 | 0.74 |
| Lamprotornis superbus* | Muscicapida | HLlamSup1 | 64.9 | 25.0 | 2.03 | 0.98 |
| Leucopsar rothschildi* | Muscicapida | HLleuRot1 | 95.0 | 25.0 | 2.19 | 0.95 |
| Oenanthe oenanthe* | Muscicapida | HLoenOen1 | 25.0 | 10.1 | 1.55 | 0.69 |
| Polioptila caerulea* | Muscicapida | HLpolCae1 | 6.5 | 4.2 | 0.91 | 0.44 |
| Regulus satrapa* | Muscicapida | HLregSat1 | 5.5 | 6.3 | 0.87 | 0.63 |
| Sialia sialis* | Muscicapida | HLsiaSia1 | 30.5 | 10.5 | 1.64 | 0.69 |
| Sitta europaea* | Muscicapida | HLsitEur1 | 20.4 | 12.9 | 1.49 | 0.81 |
| Sturnus vulgaris* | Muscicapida | HLstuVul1 | 74.0 | 22.9 | 2.08 | 0.94 |
| Thryothorus ludovicianus* | Muscicapida | HLthrLud1 | 17.5 | 9.2 | 1.39 | 0.68 |
| Toxostoma redivivum* | Muscicapida | HLtoxRed1 | 84.4 | 6.9 | 2.04 | 0.41 |

**Table S15:** Phenotypes and genome assemblies used for Passerida. *\* indicates species used in main purifying selection analyses*.

| Species | Clade/Grouping | Assembly | Adult Mass (g) | Lifespan (yr) | LLL | ELL |
| --- | --- | --- | --- | --- | --- | --- |
| Agelaius phoeniceus* | Passerida | HLagePho2 | 56.7 | 20.0 | 1.96 | 0.9 |
| Agelaius tricolor* | Passerida | HLageTri1 | 56.5 | 13.2 | 1.92 | 0.73 |
| Ammodramus savannarum* | Passerida | HLammSavPra1 | 17.6 | 9.1 | 1.4 | 0.68 |
| Amphispiza belli* | Passerida | HLartBel1 | 17.1 | 9.1 | 1.38 | 0.68 |
| Camarhynchus parvulus* | Passerida | HLcamPar1 | 13.2 | 15.0 | 1.31 | 0.91 |
| Cardinalis cardinalis* | Passerida | HLcarCar1 | 42.6 | 28.5 | 1.86 | 1.08 |
| Dendroica coronata* | Passerida | HLsetCorCor2 | 12.0 | 10.0 | 1.24 | 0.75 |
| Dendroica kirtlandii* | Passerida | HLsetKir1 | 15.5 | 9.0 | 1.34 | 0.68 |
| Erythrura gouldiae* | Passerida | HLeryGou1 | 12.0 | 6.0 | 1.2 | 0.53 |
| Fringilla coelebs* | Passerida | HLfriCoeCoe1 | 20.7 | 29.0 | 1.55 | 1.15 |
| Geothlypis trichas* | Passerida | HLgeoTri1 | 9.5 | 11.5 | 1.15 | 0.83 |
| Hemignathus munroi* | Passerida | HLhemWil1 | 28.0 | 13.0 | 1.62 | 0.78 |
| Junco hyemalis* | Passerida | HLjunHye1 | 19.0 | 11.3 | 1.45 | 0.76 |
| Lonchura striata* | Passerida | HLlonStr2 | 12.3 | 10.0 | 1.25 | 0.75 |
| Loxia curvirostra* | Passerida | HLloxCur1 | 39.5 | 16.1 | 1.79 | 0.84 |
| Loxia leucoptera* | Passerida | HLloxLeu1 | 28.9 | 4.0 | 1.54 | 0.28 |
| Molothrus ater* | Passerida | HLmolAte1 | 38.1 | 16.9 | 1.77 | 0.87 |
| Motacilla alba* | Passerida | HLmotAlbAlb2 | 24.1 | 13.7 | 1.56 | 0.82 |
| Passer domesticus* | Passerida | HLpasDom1 | 25.3 | 23.0 | 1.62 | 1.04 |
| Passer montanus* | Passerida | HLpasMon1 | 20.5 | 13.1 | 1.49 | 0.82 |
| Passerculus sandwichensis* | Passerida | HLpasSanAla1 | 20.2 | 18.0 | 1.51 | 0.95 |
| Passerina amoena* | Passerida | HLpasAmo1 | 15.2 | 9.9 | 1.34 | 0.72 |
| Pheucticus melanocephalus* | Passerida | HLpheMel1 | 42.0 | 25.0 | 1.85 | 1.02 |
| Quiscalus mexicanus* | Passerida | HLquiMex1 | 168.7 | 12.5 | 2.39 | 0.6 |
| Serinus canaria* | Passerida | HLserCan1 | 24.3 | 24.0 | 1.61 | 1.06 |
| Taeniopygia guttata* | Passerida | HLtaeGut4 | 12.0 | 12.0 | 1.25 | 0.83 |
| Zonotrichia albicollis* | Passerida | zonAlb1 | 21.1 | 14.9 | 1.51 | 0.87 |

**Table S16:**
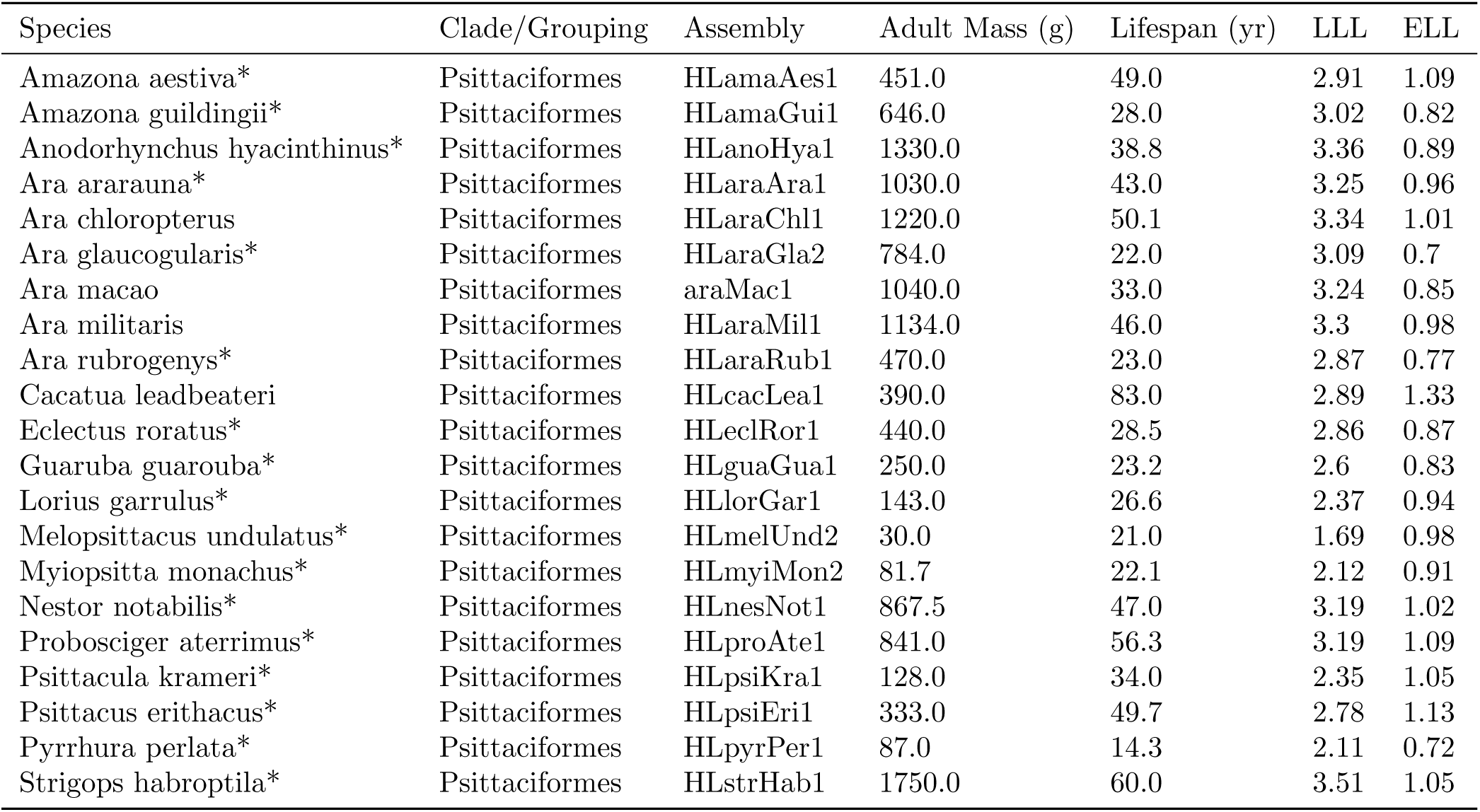
Phenotypes and genome assemblies used for Psittaciformes. *\* indicates species used in main purifying selection analyses*.

**Table S17:** Phenotypes and genome assemblies used for Sylviida. *\* indicates species used in main purifying selection analyses*.

| Species | Clade/Grouping | Assembly | Adult Mass (g) | Lifespan (yr) | LLL | ELL |
| --- | --- | --- | --- | --- | --- | --- |
| Acrocephalus arundinaceus* | Sylviida | HLacrAru1 | 30.0 | 10.1 | 1.63 | 0.67 |
| Acrocephalus scirpaceus* | Sylviida | HLacrSciSci1 | 12.3 | 16.9 | 1.29 | 0.97 |
| Aegithalos caudatus* | Sylviida | HLaegCau1 | 8.6 | 11.1 | 1.11 | 0.82 |
| Alauda arvensis* | Sylviida | HLalaArv1 | 37.5 | 10.1 | 1.73 | 0.65 |
| Cettia cetti* | Sylviida | HLcetCet1 | 13.3 | 9.3 | 1.28 | 0.71 |
| Eremophila alpestris* | Sylviida | HLereAlp1 | 33.5 | 8.0 | 1.66 | 0.56 |
| Hippolais icterina* | Sylviida | HLhipIct1 | 13.2 | 10.8 | 1.29 | 0.77 |
| Hirundo rustica* | Sylviida | HLhirRus2 | 18.3 | 16.0 | 1.46 | 0.91 |
| Leiothrix lutea* | Sylviida | HLleiLut1 | 21.5 | 15.0 | 1.52 | 0.87 |
| Parus atricapillus* | Sylviida | HLpoeAtr1 | 10.4 | 12.4 | 1.19 | 0.85 |
| Parus caeruleus* | Sylviida | HLcyaCae1 | 10.3 | 14.6 | 1.2 | 0.92 |
| Parus major* | Sylviida | HLparMaj1 | 17.9 | 15.4 | 1.44 | 0.9 |
| Phylloscopus trochilus* | Sylviida | HLphyTro1 | 8.7 | 11.8 | 1.11 | 0.85 |
| Progne subis* | Sylviida | HLproSubSub1 | 45.0 | 13.8 | 1.83 | 0.77 |
| Pycnonotus jocosus* | Sylviida | HLpycJoc1 | 27.7 | 11.1 | 1.6 | 0.72 |
| Riparia riparia | Sylviida | HLripRip1 | 12.7 | 10.1 | 1.26 | 0.75 |
| Sylvia atricapilla* | Sylviida | HLsylAtr1 | 16.7 | 13.8 | 1.41 | 0.86 |
| Sylvia borin* | Sylviida | HLsylBor1 | 24.6 | 24.0 | 1.61 | 1.06 |
| Zosterops japonicus* | Sylviida | HLzosJap1 | 11.2 | 5.1 | 1.16 | 0.47 |
| Zosterops lateralis* | Sylviida | HLzosLat1 | 12.7 | 18.7 | 1.31 | 1.01 |

**Table S18:**
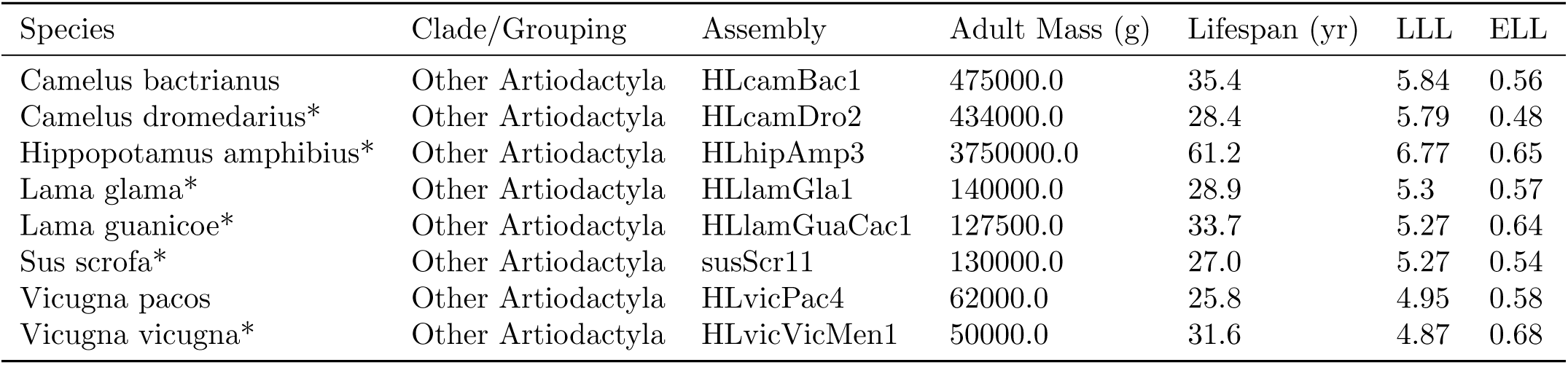
Phenotypes and genome assemblies used for other Artiodactyla. *\* indicates species used in main purifying selection analyses*.

**Table S19:**
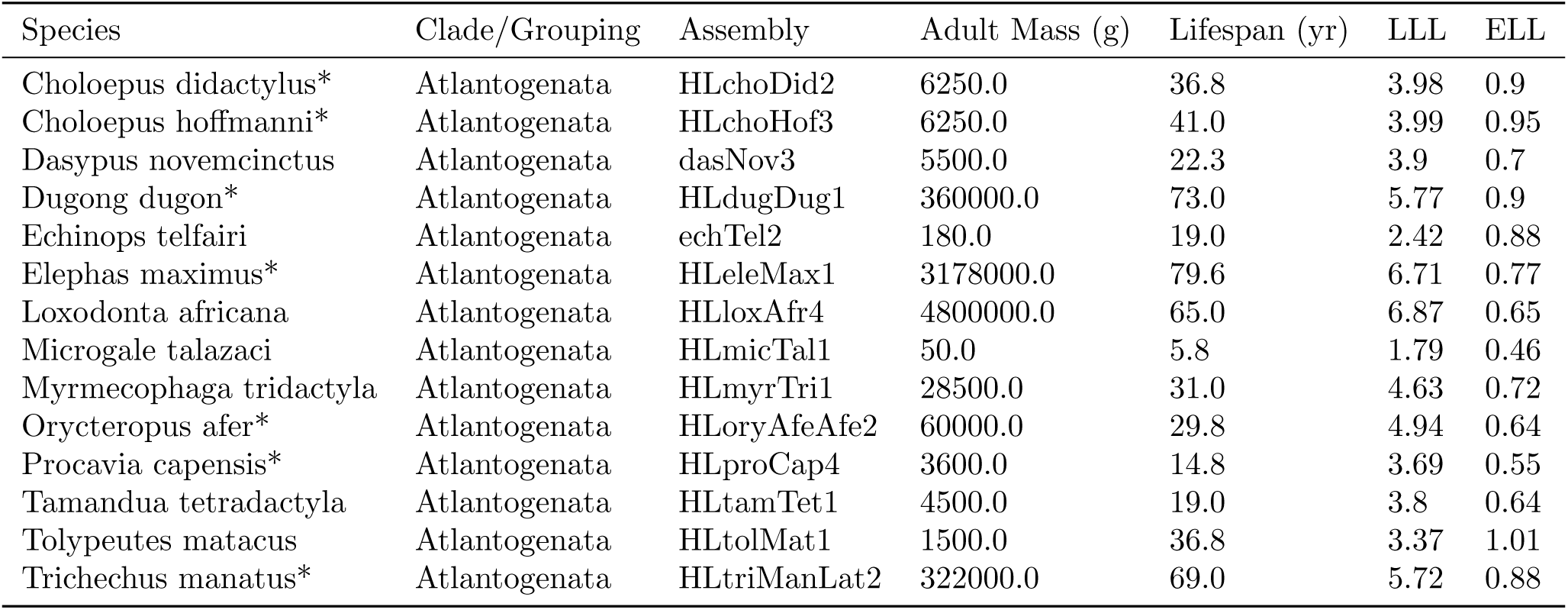
Phenotypes and genome assemblies used for Atlantogenata. *\* indicates species used in main purifying selection analyses*.

**Table S20:**
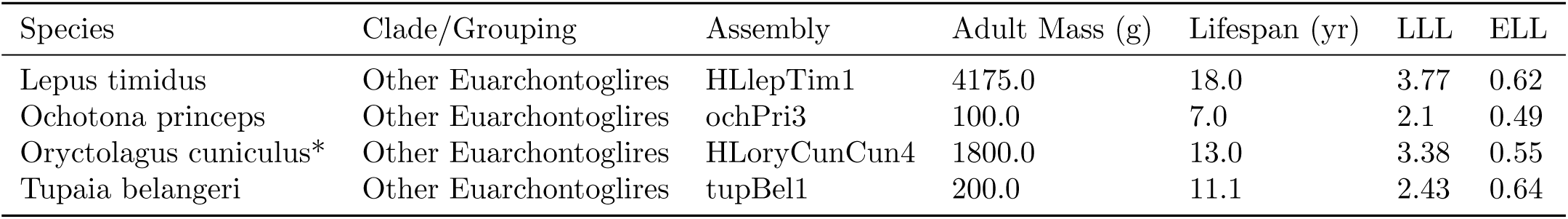
Phenotypes and genome assemblies used for other Euarchontoglires. *\* indicates species used in main purifying selection analyses*.

| Species | Clade/Grouping | Assembly | Adult Mass (g) | Lifespan (yr) | LLL | ELL |
| --- | --- | --- | --- | --- | --- | --- |
| Lepus timidus | Other Euarchontoglires | HLlepTim1 | 4175.0 | 18.0 | 3.77 | 0.62 |
| Ochotona princeps | Other Euarchontoglires | ochPri3 | 100.0 | 7.0 | 2.1 | 0.49 |
| Oryctolagus cuniculus* | Other Euarchontoglires | HLoryCunCun4 | 1800.0 | 13.0 | 3.38 | 0.55 |
| Tupaia belangeri | Other Euarchontoglires | tupBel1 | 200.0 | 11.1 | 2.43 | 0.64 |

**Table S21:** Phenotypes and genome assemblies used for Eulipotyphla. *\* indicates species used in main purifying selection analyses*.

| Species | Clade/Grouping | Assembly | Adult Mass (g) | Lifespan (yr) | LLL | ELL |
| --- | --- | --- | --- | --- | --- | --- |
| Condylura cristata | Eulipotyphla | conCri1 | 55.3 | 2.5 | 1.78 | 0.1 |
| Erinaceus europaeus* | Eulipotyphla | eriEur2 | 750.0 | 11.7 | 3.0 | 0.57 |
| Scalopus aquaticus* | Eulipotyphla | HLscaAqu1 | 90.0 | 6.2 | 2.05 | 0.45 |
| Solenodon paradoxus* | Eulipotyphla | HLsolPar1 | 1000.0 | 12.1 | 3.13 | 0.56 |
| Sorex araneus | Eulipotyphla | sorAra2 | 9.0 | 3.2 | 1.02 | 0.34 |

**Table S22:** Phenotypes and genome assemblies used for Falco. *\* indicates species used in main purifying selection analyses.*

| Species | Clade/Grouping | Assembly | Adult Mass (g) | Lifespan (yr) | LLL | ELL |
| --- | --- | --- | --- | --- | --- | --- |
| Falco cherrug* | Falco | falChe1 | 966.0 | 15.9 | 3.15 | 0.55 |
| Falco naumanni* | Falco | HLfalNau1 | 152.5 | 10.9 | 2.33 | 0.55 |
| Falco peregrinus* | Falco | falPer1 | 840.0 | 25.0 | 3.13 | 0.75 |
| Falco rusticolus* | Falco | HLfalRus1 | 1406.5 | 13.5 | 3.3 | 0.44 |
| Falco tinnunculus* | Falco | HLfalTin1 | 184.0 | 23.8 | 2.47 | 0.87 |

**Table S23:**
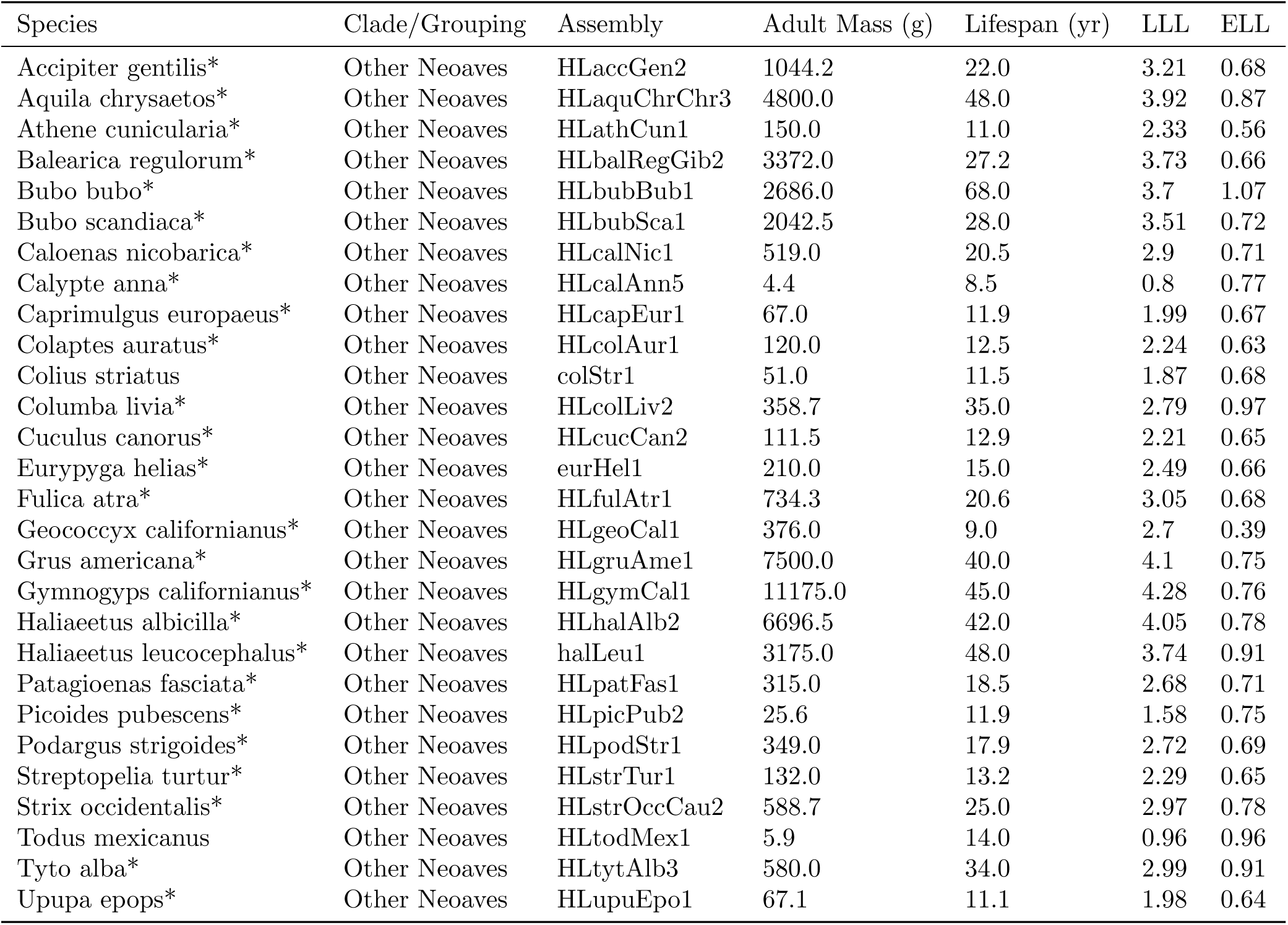
Phenotypes and genome assemblies used for other Neoaves. *\* indicates species used in main purifying selection analyses*.

**Table S24:** Phenotypes and genome assemblies used for Paleognathae. *\* indicates species used in main purifying selection analyses*.

| Species | Clade/Grouping | Assembly | Adult Mass (g) | Lifespan (yr) | LLL | ELL |
| --- | --- | --- | --- | --- | --- | --- |
| Casuarius casuarius* | Palaeognathae | HLcasCas2 | 44000.0 | 30.0 | 4.83 | 0.47 |
| Dromaius novaehollandiae* | Palaeognathae | HLdroNov2 | 38925.0 | 16.6 | 4.73 | 0.23 |
| Struthio camelus* | Palaeognathae | strCam1 | 111000.0 | 50.0 | 5.27 | 0.6 |

**Table S25:** Phenotypes and genome assemblies used for other Passeriformes. *\* indicates species used in main purifying selection analyses*.

| Species | Clade/Grouping | Assembly | Adult Mass (g) | Lifespan (yr) | LLL | ELL |
| --- | --- | --- | --- | --- | --- | --- |
| Drymodes brunneopygia* | Other Passeriformes | HLdryBru1 | 34.0 | 9.7 | 1.68 | 0.64 |
| Entomyzon cyanotis* | Other Passeriformes | HLentCya1 | 101.4 | 9.1 | 2.14 | 0.51 |
| Eopsaltria australis* | Other Passeriformes | HLeopAus2 | 20.2 | 14.7 | 1.49 | 0.87 |
| Origma solitaria* | Other Passeriformes | HLoriSol1 | 14.4 | 7.0 | 1.29 | 0.58 |
| Orthonyx spaldingii* | Other Passeriformes | HLortSpa1 | 145.1 | 16.5 | 2.34 | 0.73 |
| Ptilonorhynchus violaceus* | Other Passeriformes | HLptiVio1 | 213.0 | 25.1 | 2.54 | 0.88 |

**Figure S1:**
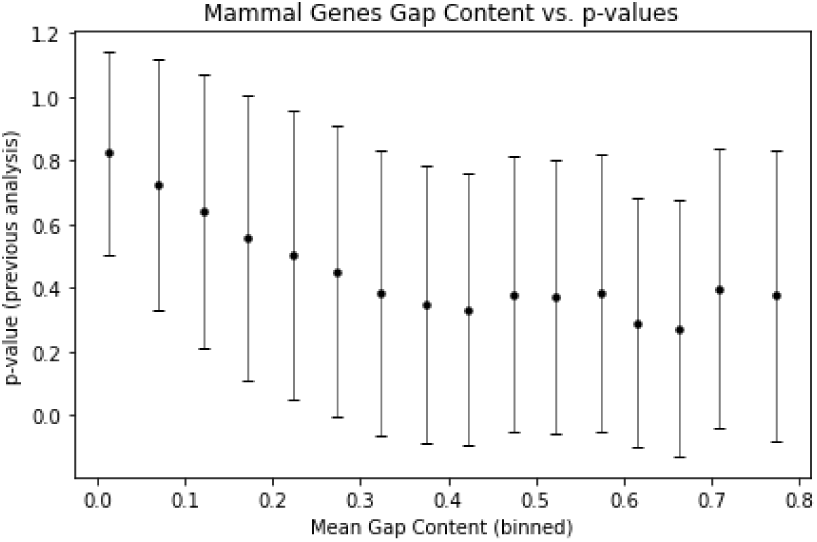
Decreasing mean p-values in genes with higher proportional gap content in mammals.

**Figure S2:**
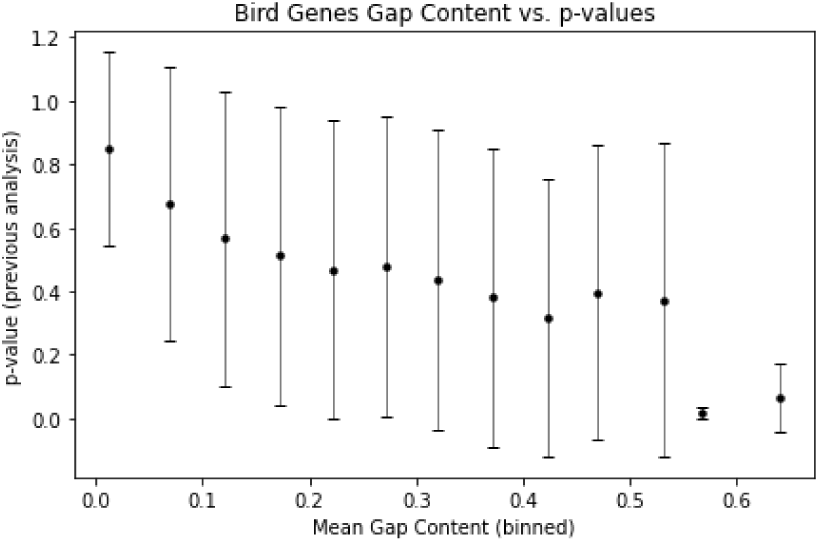
Decreasing mean p-values in genes with higher proportional gap content in bird.

**Figure S3:**
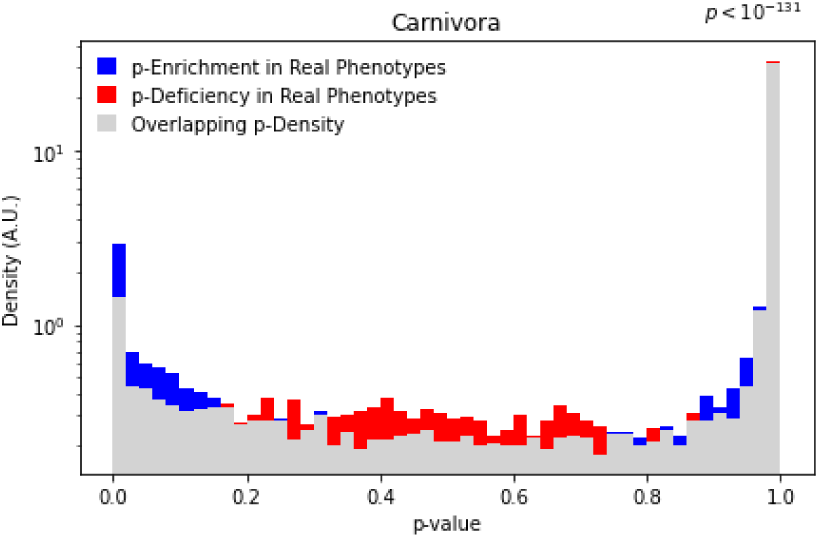
Distribution of p-values in the LLL Carnivora positive selection analysis, for the actual phenotypes compared to randomly permulated phenotypes.

**Figure S4:**
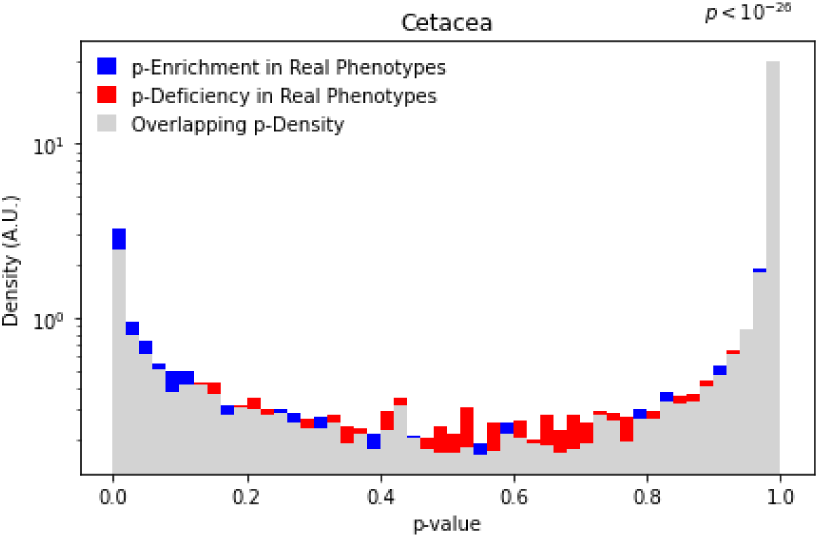
Distribution of p-values in the LLL Cetacea positive selection analysis, for the actual phenotypes compared to randomly permulated phenotypes.

**Figure S5:**
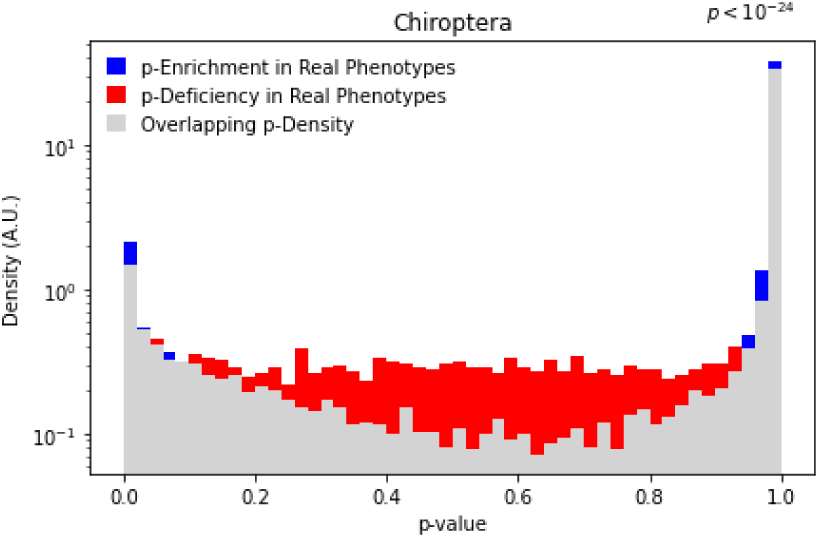
Distribution of p-values in the LLL Chiroptera positive selection analysis, for the actual phenotypes compared to randomly permulated phenotypes.

**Figure S6:**
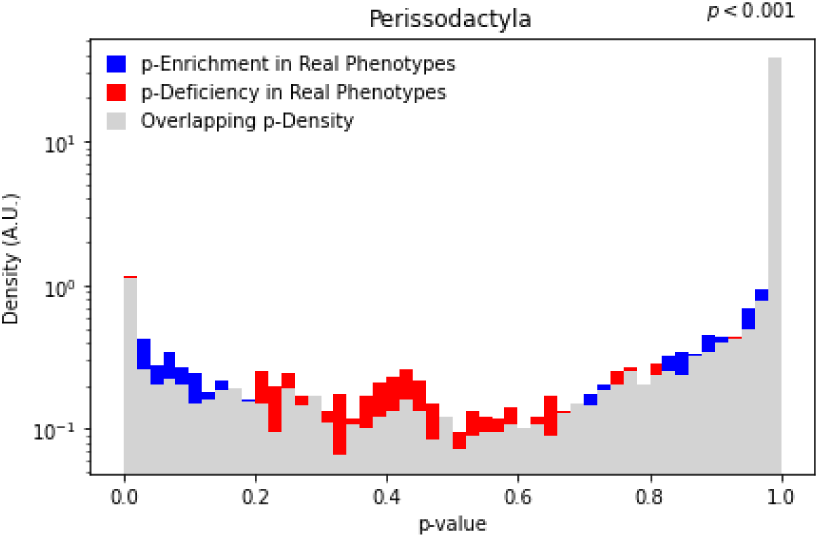
Distribution of p-values in the LLL Perissodactyla positive selection analysis, for the actual phenotypes compared to randomly permulated phenotypes.

**Figure S7:**
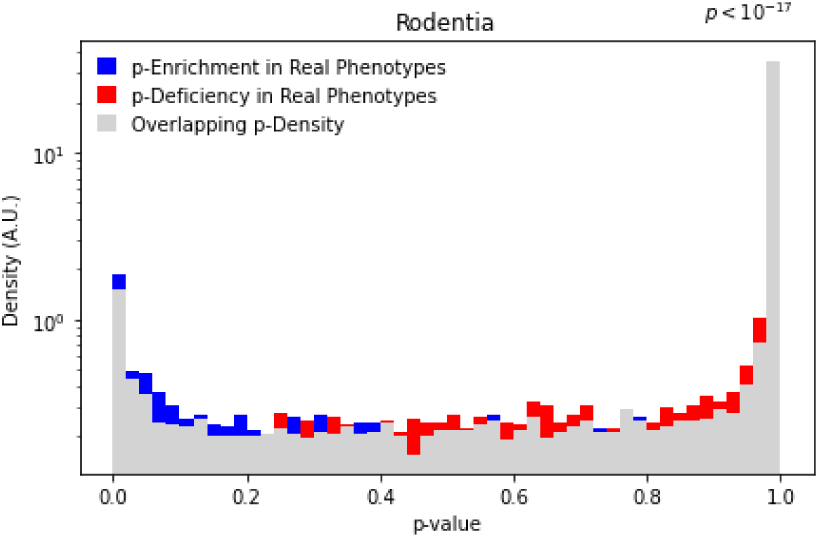
Distribution of p-values in the LLL Rodentia positive selection analysis, for the actual phenotypes compared to randomly permulated phenotypes.

**Figure S8:**
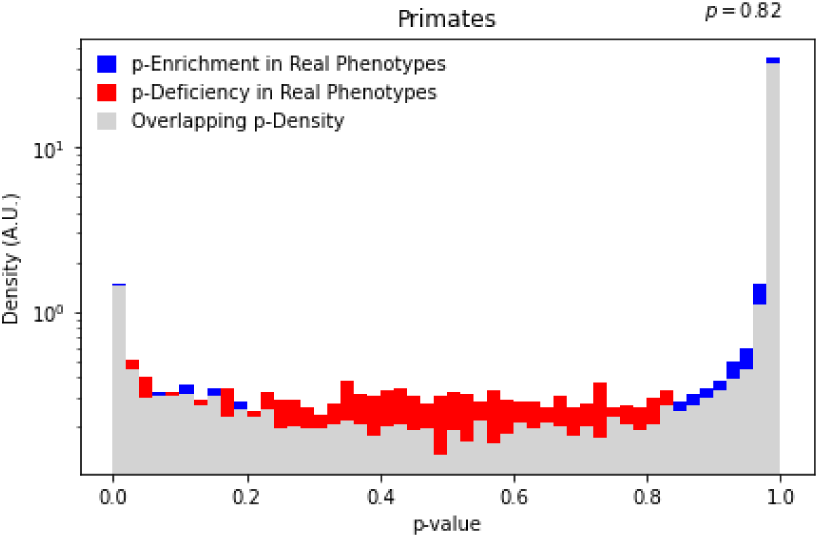
Distribution of p-values in the LLL Primates positive selection analysis, for the actual phenotypes compared to randomly permulated phenotypes.

**Figure S9:**
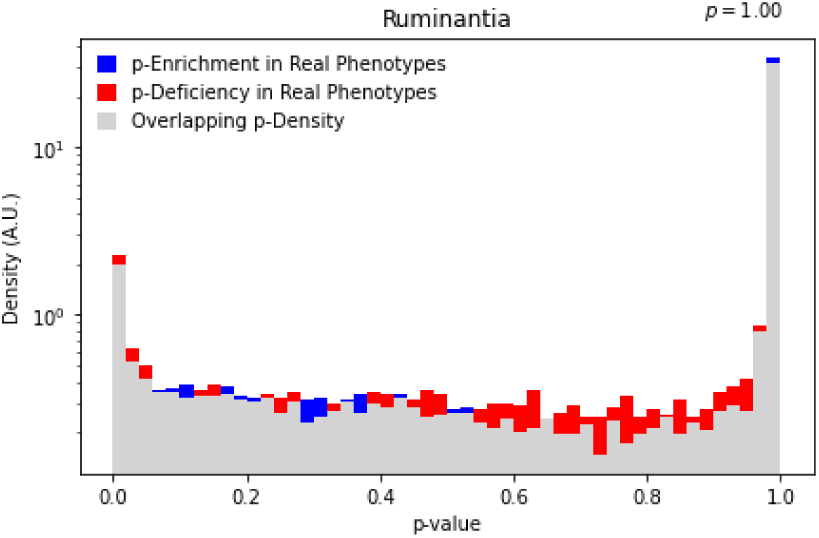
Distribution of p-values in the LLL Ruminantia positive selection analysis, for the actual phenotypes compared to randomly permulated phenotypes.

**Figure S10:**
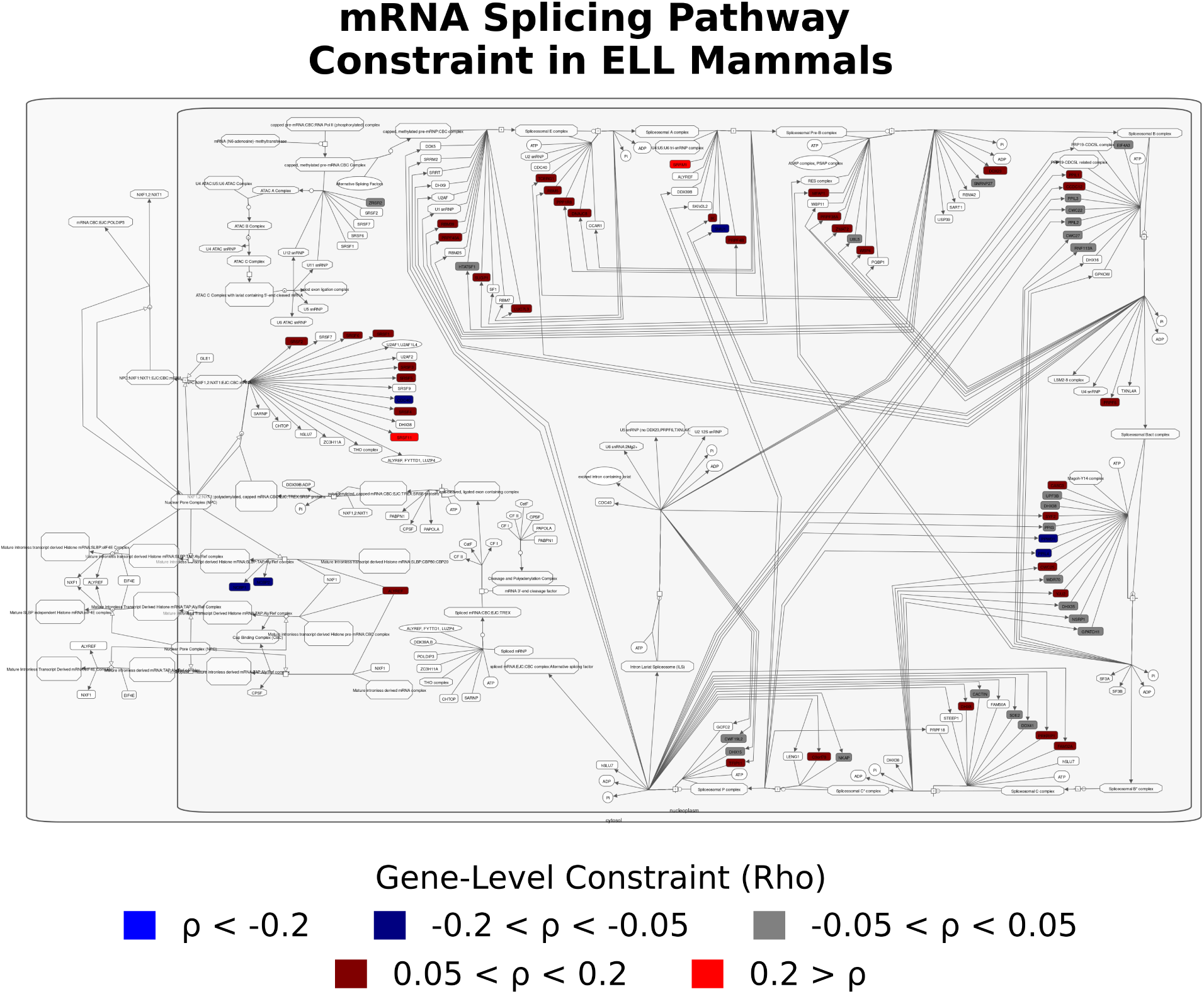
Gene level constraint for individual genes in the Reactome mRNA Splicing pathway annotation in ELL mammals. These are highlighted using custom python code, working from a modification of Reactome’s pathway browser [35] format as exported from the web-based Newt Editor [80] in an XML-based format (NWT 0.3). Complexes of multiple genes or other elements, or single genes not included in our gene set, are left white.

**Figure S11:**
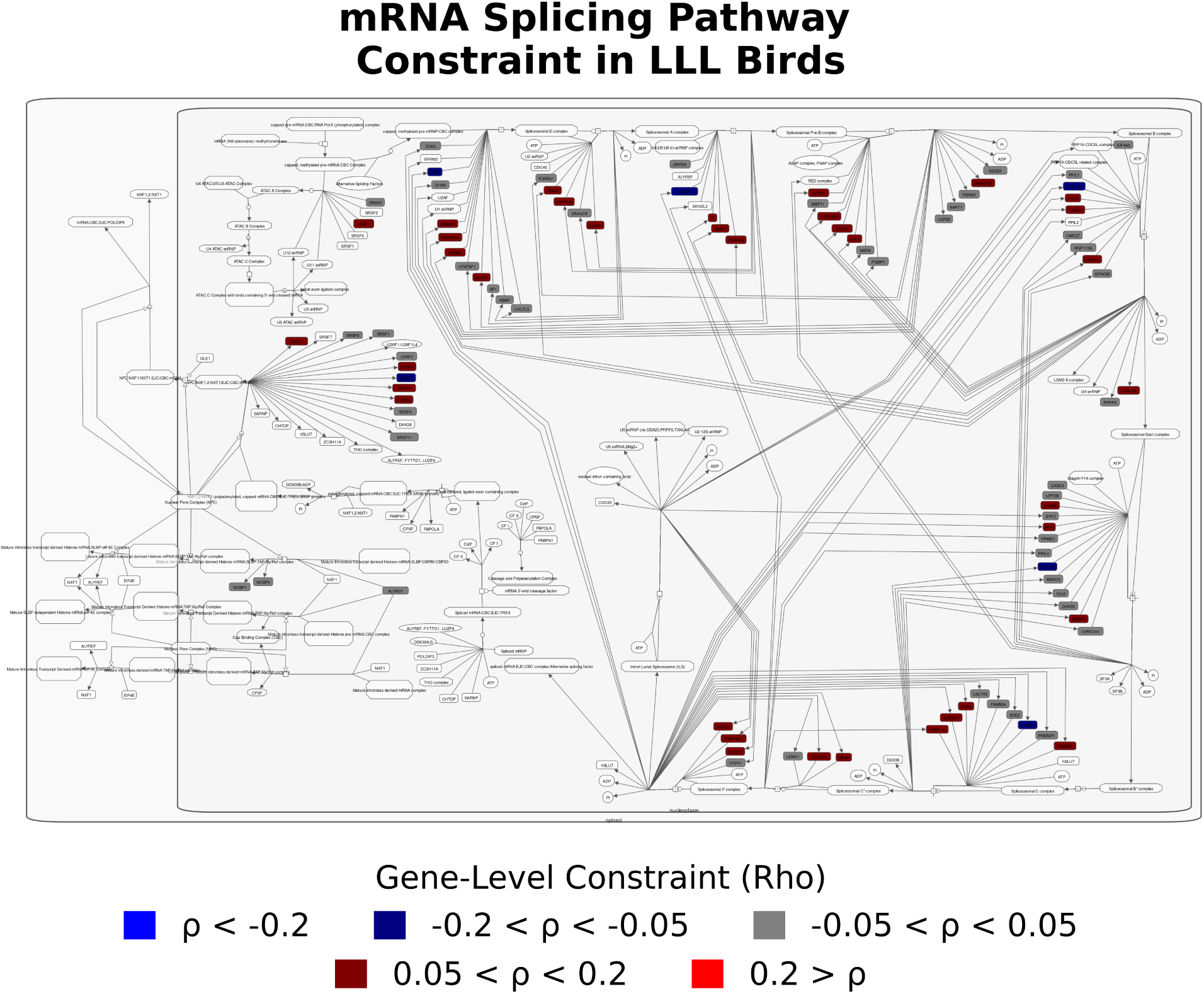
Gene level constraint for individual genes in the Reactome mRNA Splicing pathway annotation in LLL birds. These are highlighted using custom python code, working from a modification of Reactome’s pathway browser [35] format as exported from the web-based Newt Editor [80] in an XML-based format (NWT 0.3). Complexes of multiple genes or other elements, or single genes not included in our gene set, are left white.

**Figure S12:**
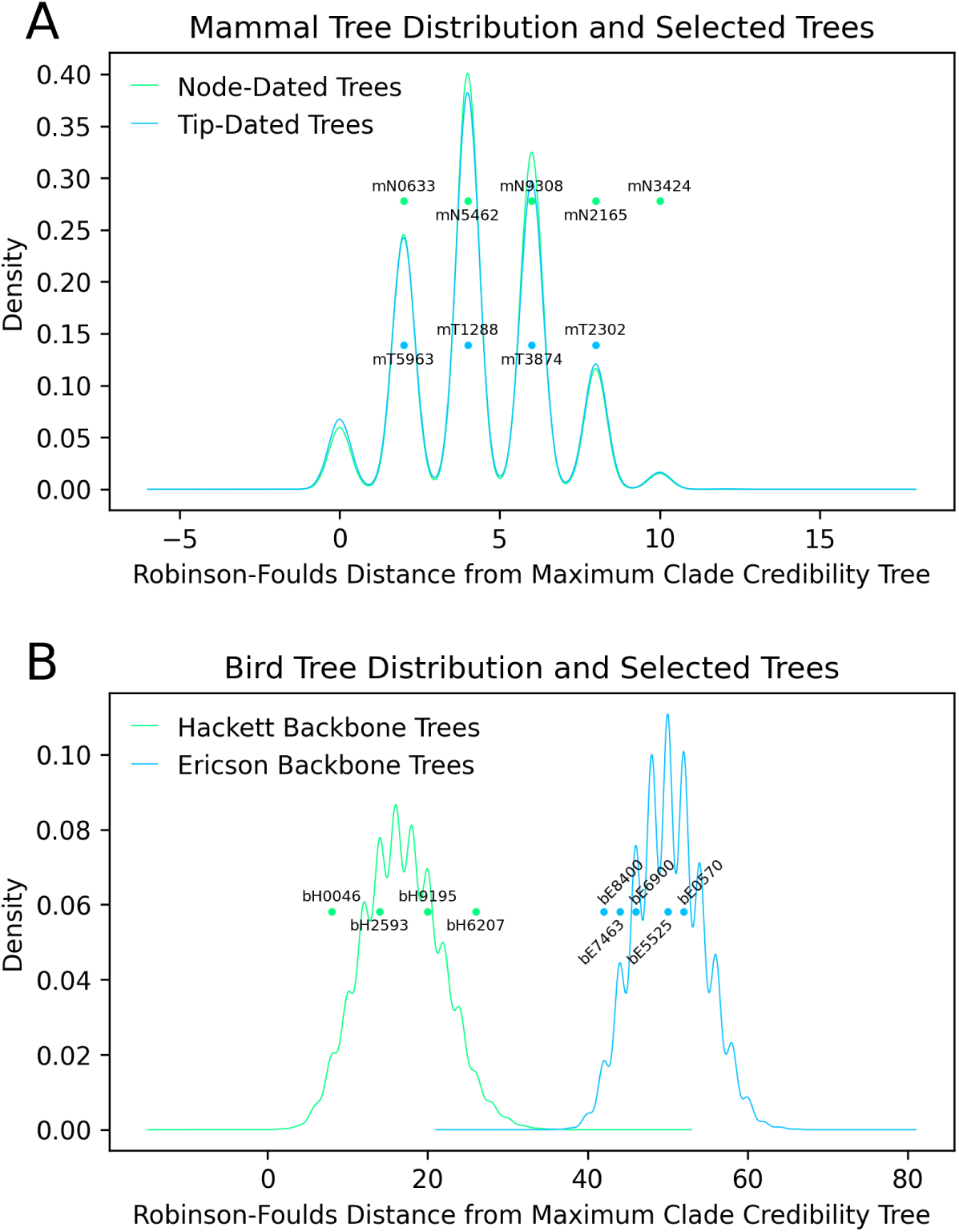
**Distributions of Robinson-Foulds distances between the main tree used in our analyses and 20,000 alternate trees** *for (A) mammals (10,000 node-dated trees and 10,000 tip-dated trees) and (B) birds (10,000 Hackett backbone trees and 10,000 Ericson backbone trees).*

**Figure S13:**
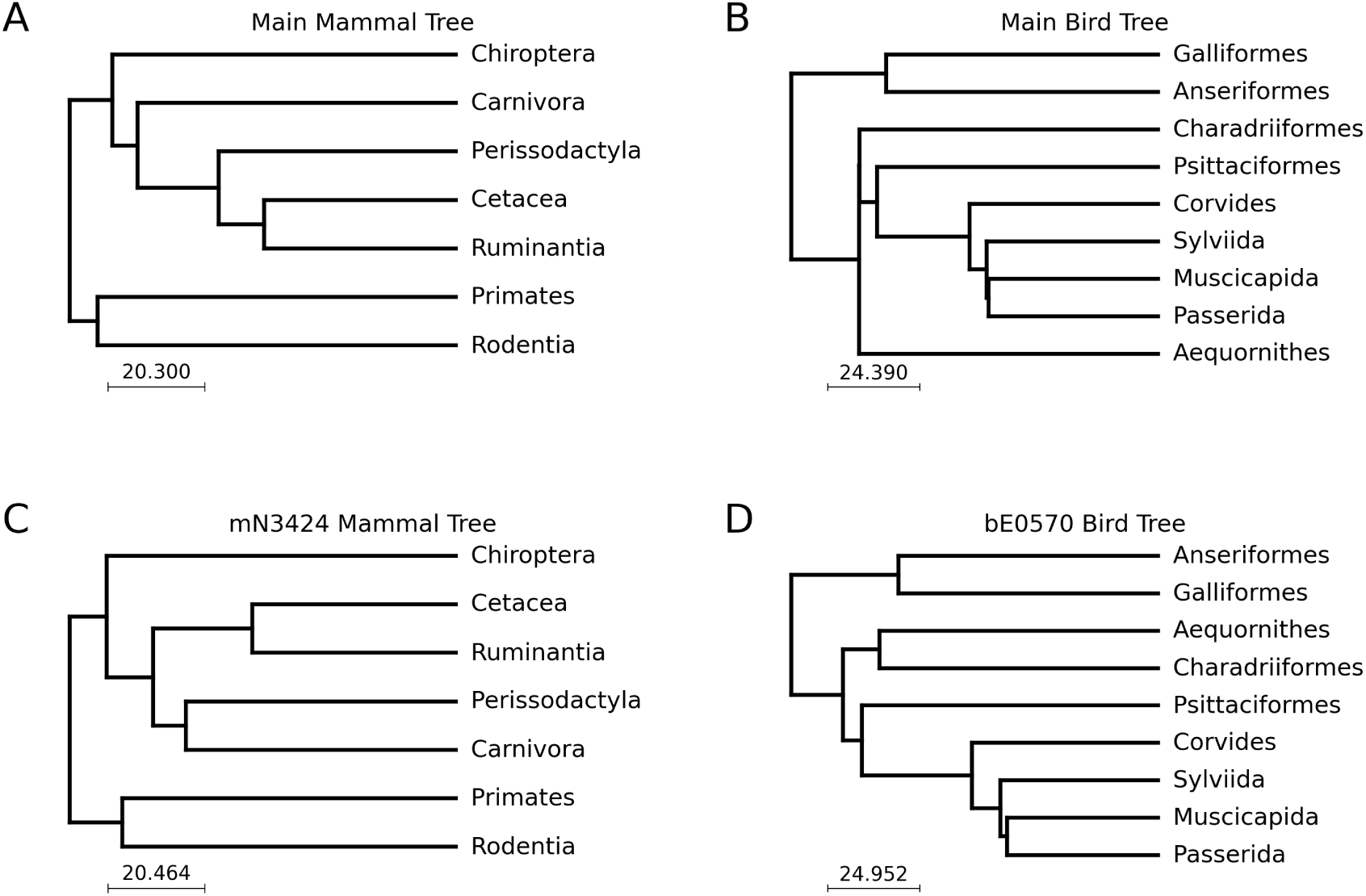
Ancestral relationships among clades selected for positive selection tests. (A) All selected mammal trees, except for mN3424, have the same clade-level topology as the main mammal tree: mT1288, mT2302, mT3874, mT5963, mN0633, mN2165, mN5462, and mN9308. (B) All selected “Hackett” backbone trees have the same clade-level topology as the main bird tree: bH0046, bH2593, bH6207, and bH9195. (C) mN3424 has an alternative phylogeny where Carnivora and Perissodactyla are sister clades. (D) All selected “Ericson” backbone trees share the same clade-level topology as bE0570: bE5525, bE6900, bE7463, and bE8400. In contrast to the main bird tree, Charadriiformes and Aequornithes are sister clades.

**Figure S14:**
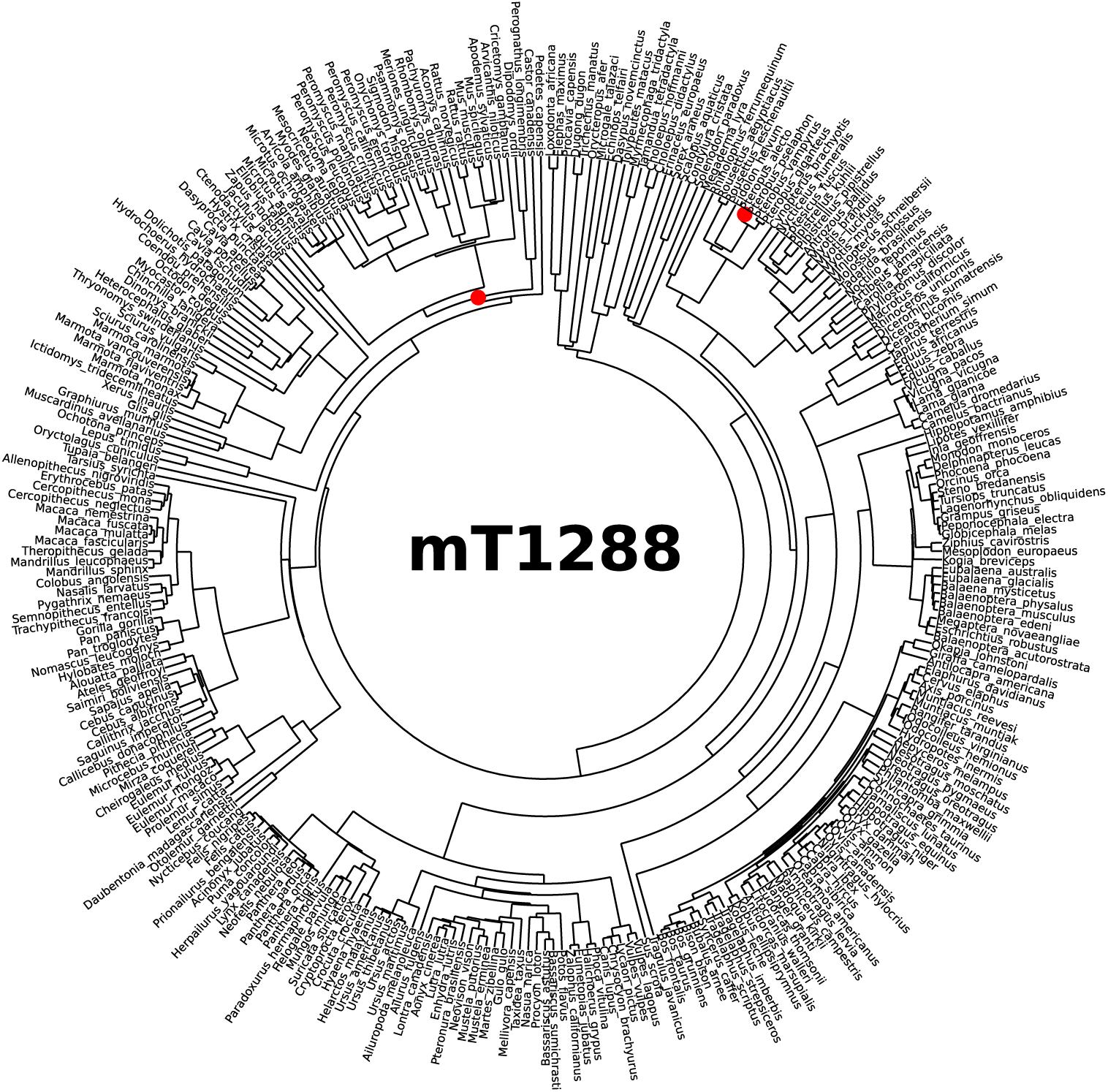
Mammal tip-dated tree 1288 (mT1288), with nodes involved in topology differences from the main tree highlighted in red.

**Figure S15:**
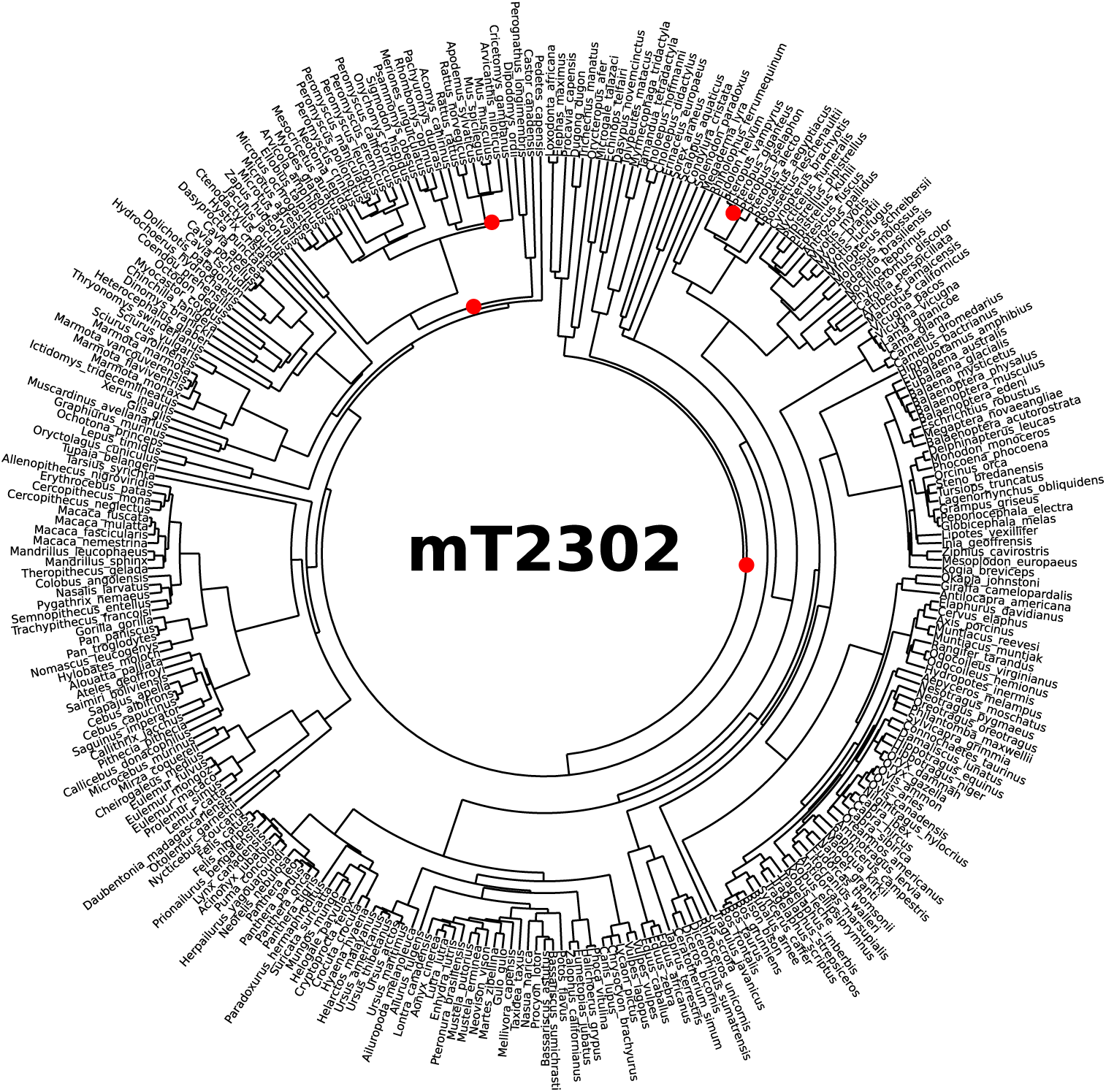
Mammal tip-dated tree 2302 (mT2302), with nodes involved in topology differences from the main tree highlighted in red.

**Figure S16:**
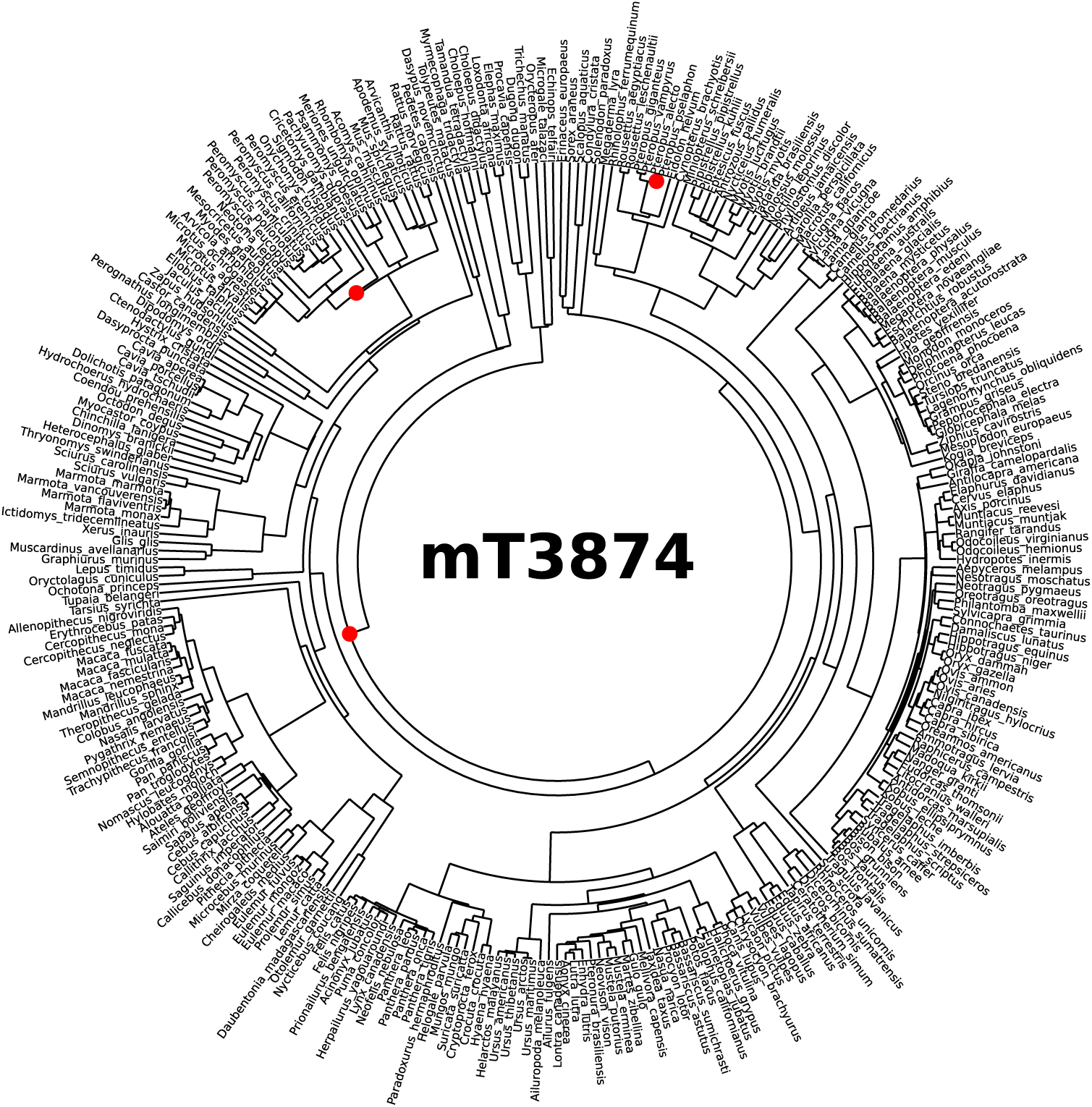
Mammal tip-dated tree 3874 (mT3874), with nodes involved in topology differences from the main tree highlighted in red.

**Figure S17:**
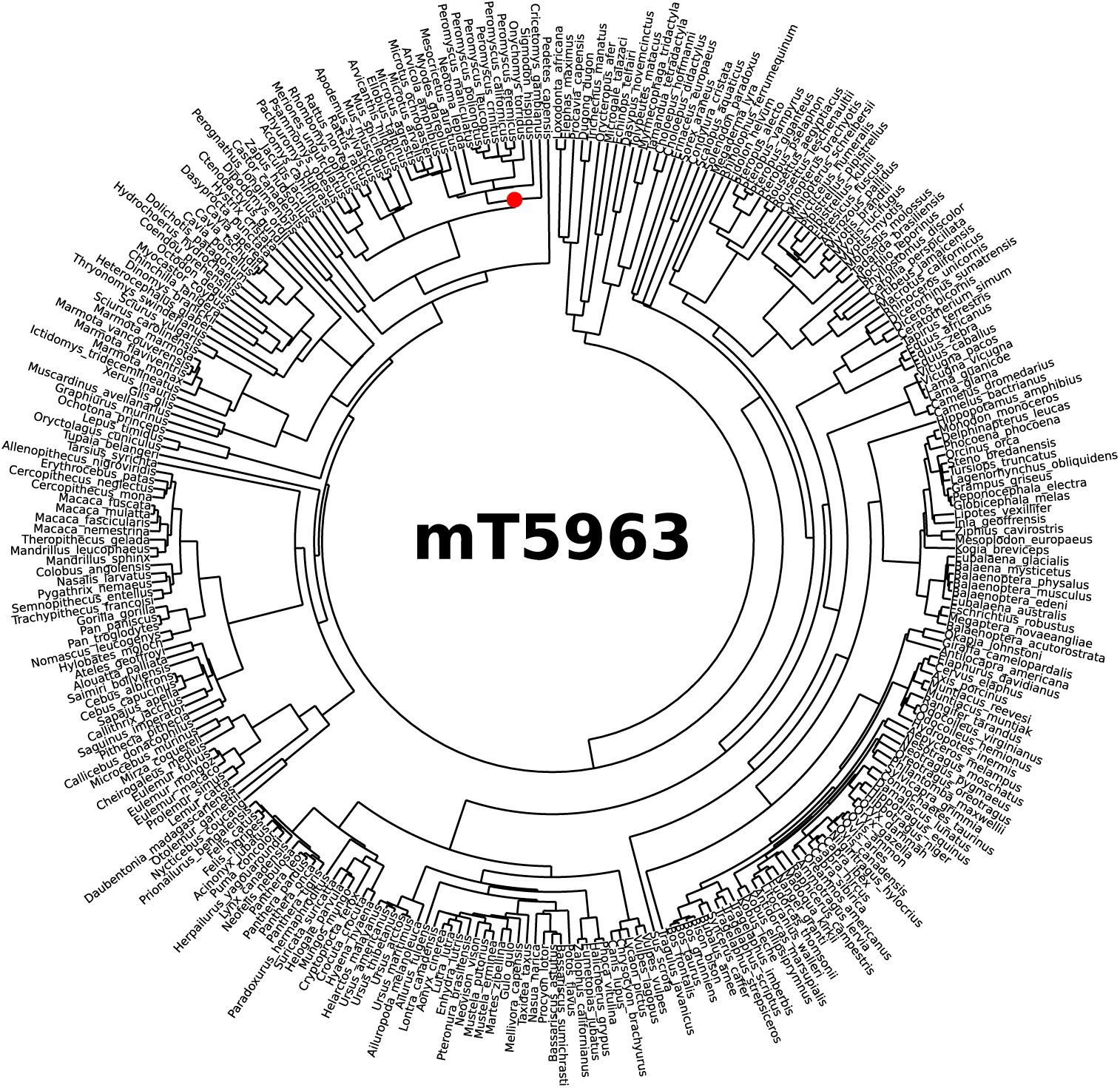
Mammal tip-dated tree 5963 (mT5963), with nodes involved in topology differences from the main tree highlighted in red.

**Figure S18:**
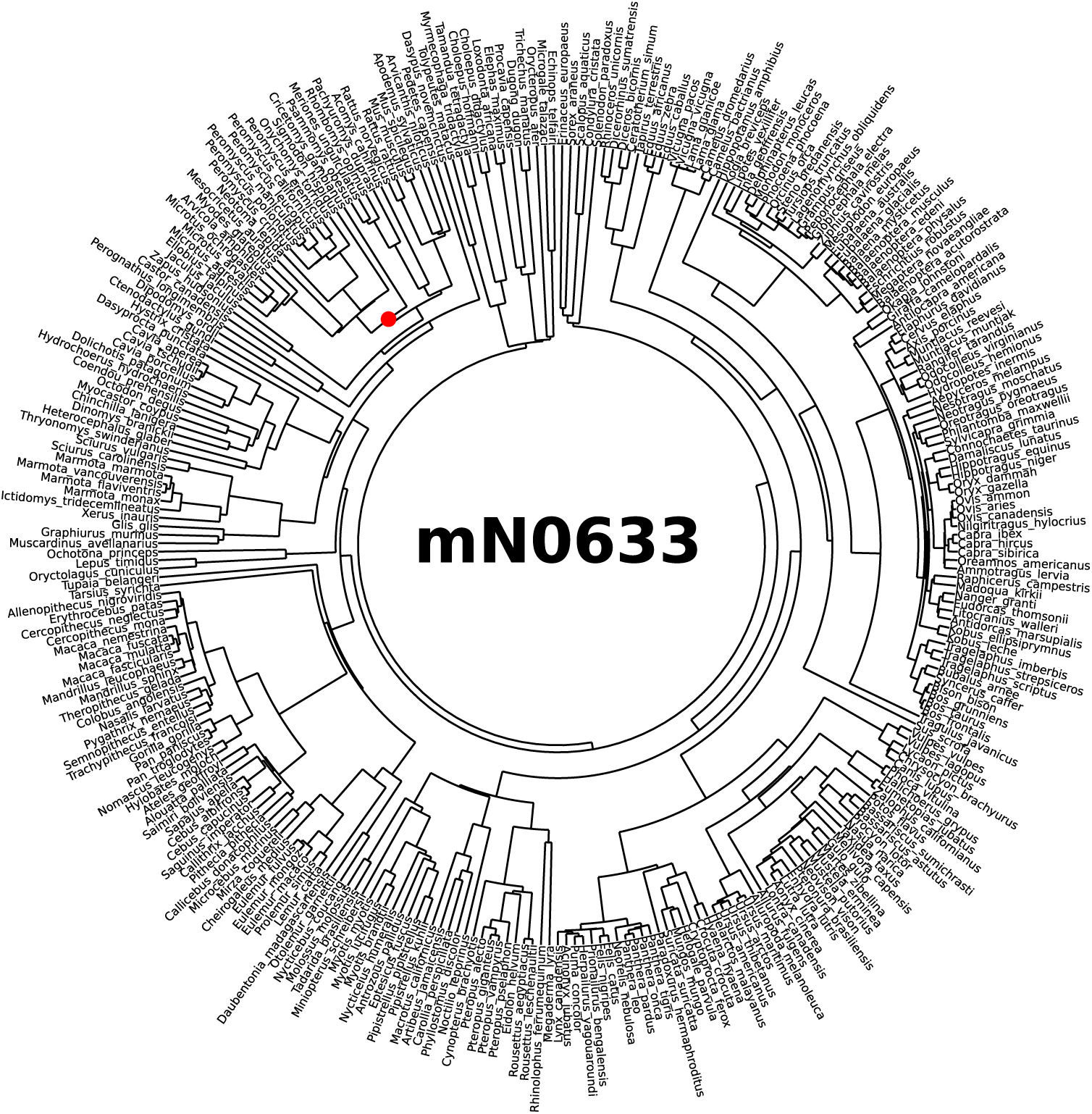
Mammal node-dated tree 0633 (mN0633), with nodes involved in topology differences from the main tree highlighted in red.

**Figure S19:**
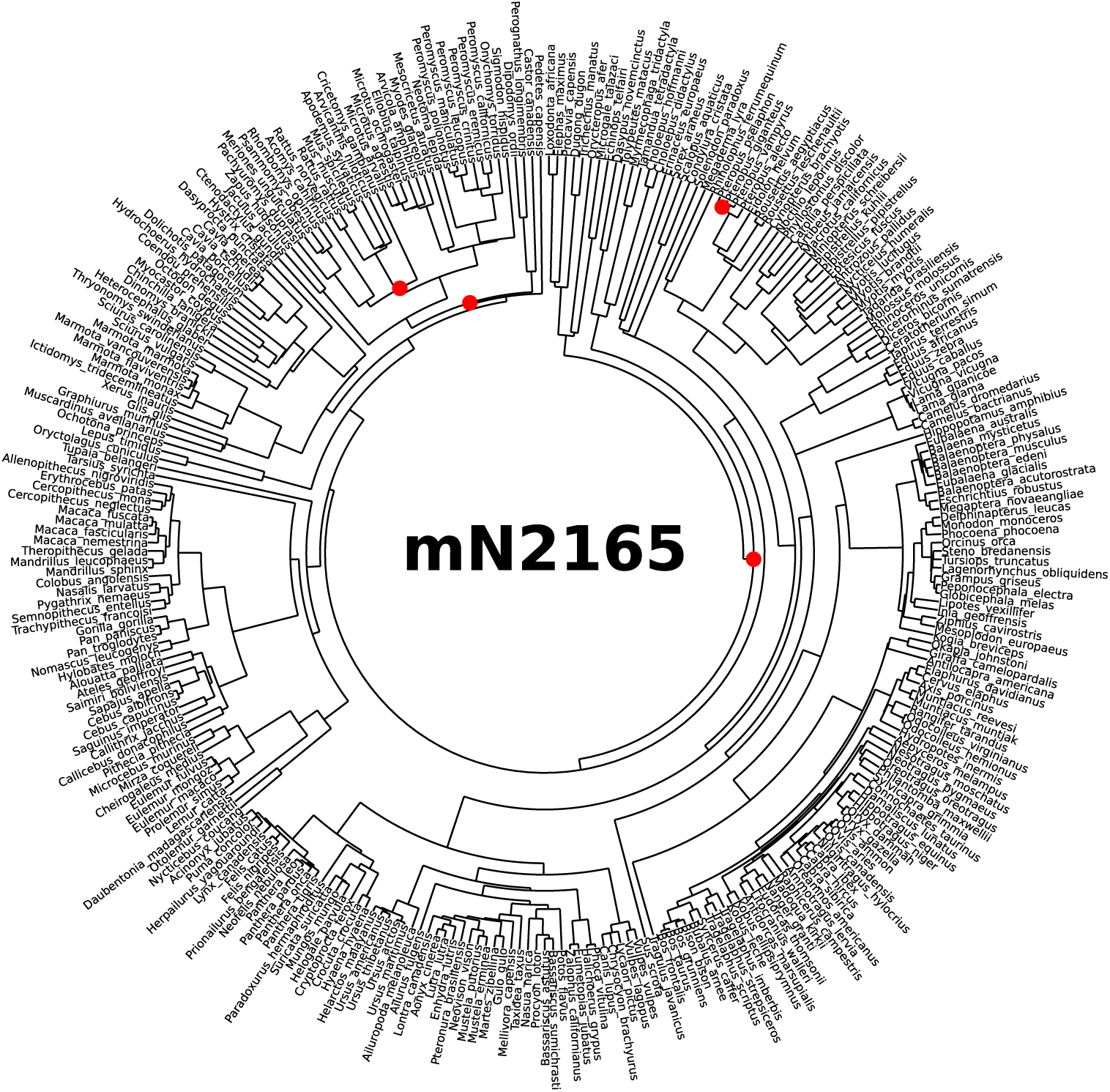
Mammal node-dated tree 2165 (mN2165), with nodes involved in topology differences from the main tree highlighted in red.

**Figure S20:**
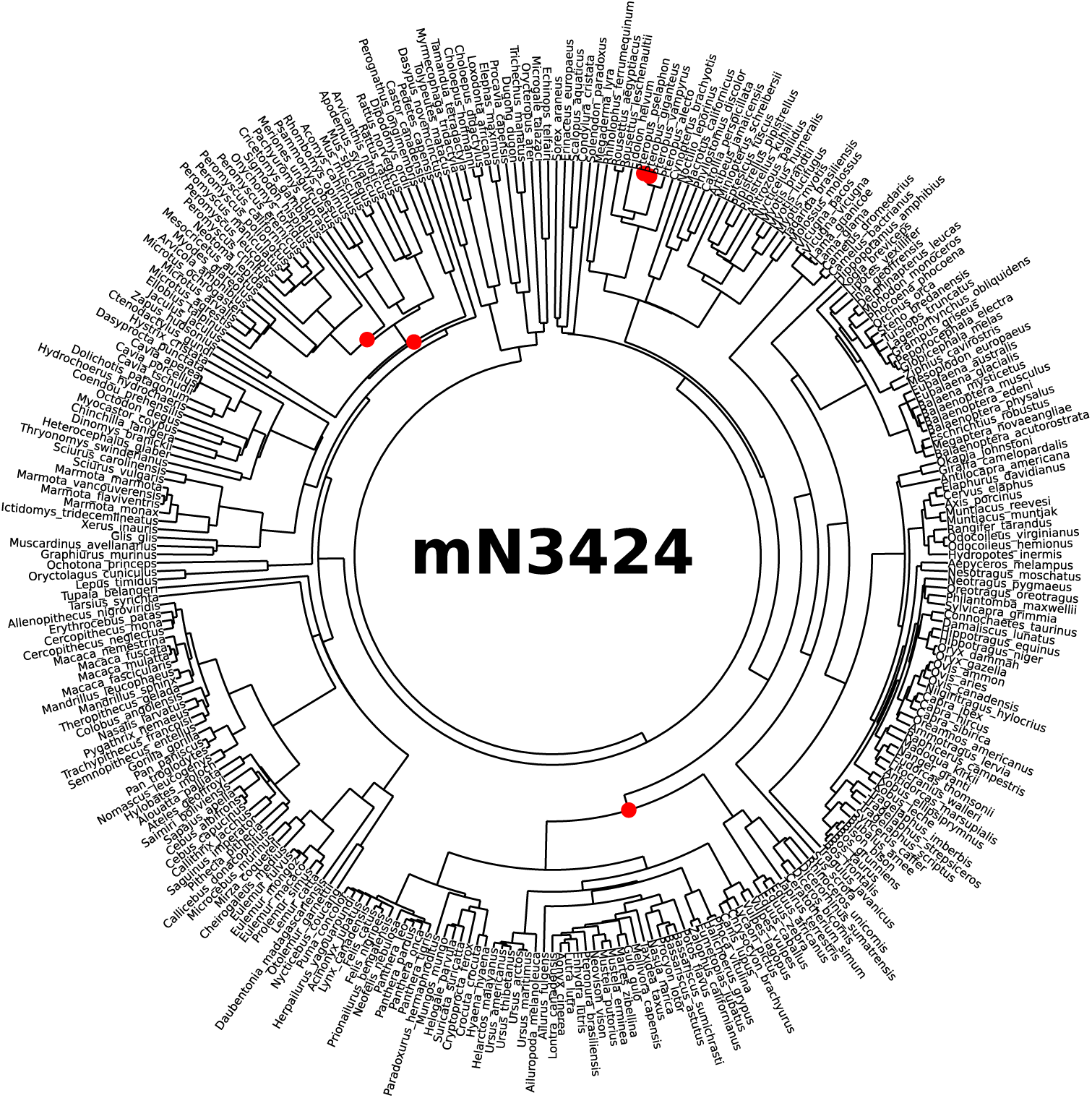
Mammal node-dated tree 3424 (mN3424), with nodes involved in topology differences from the main tree highlighted in red.

**Figure S21:**
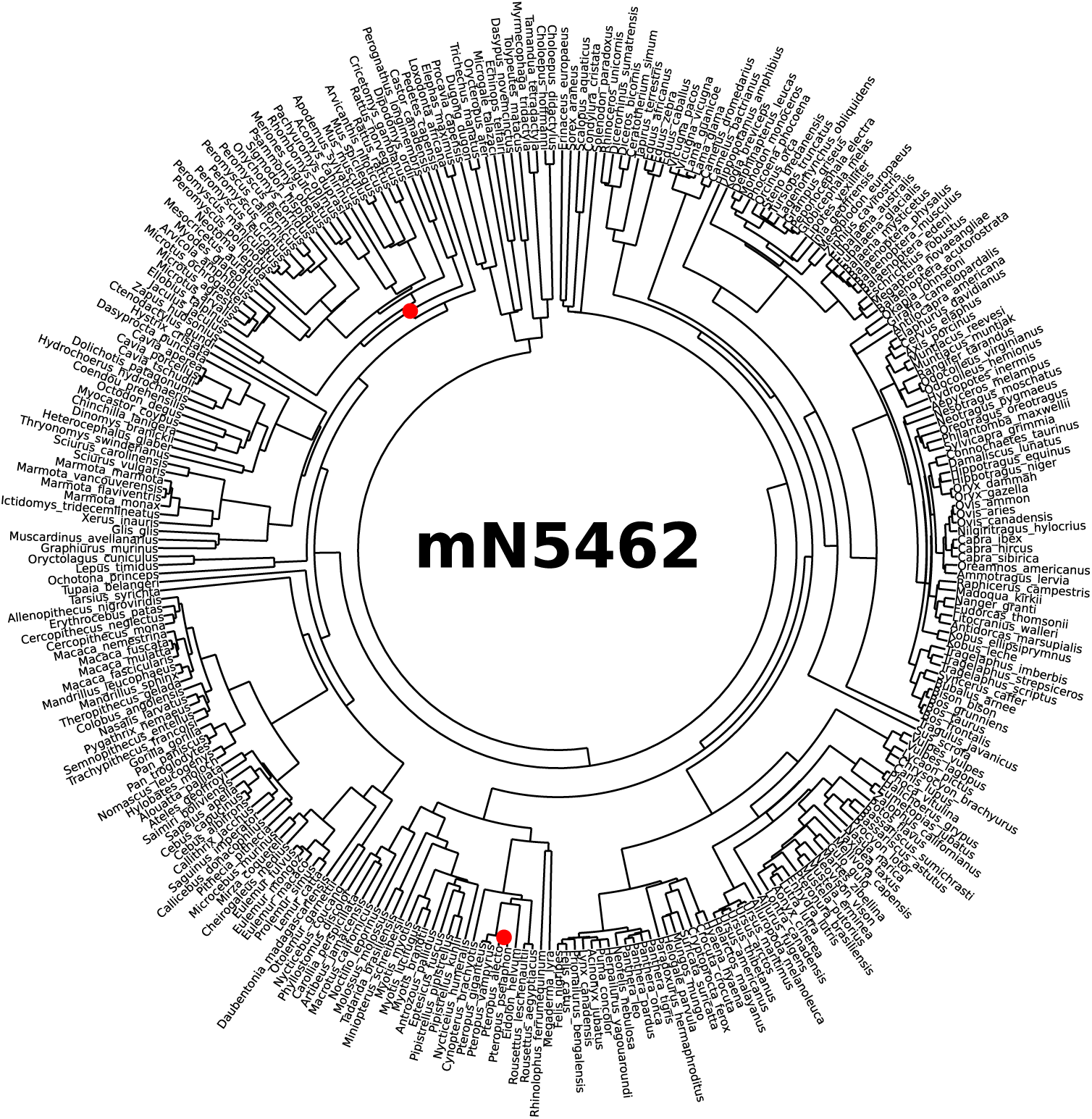
Mammal node-dated tree 5462 (mN5462), with nodes involved in topology differences from the main tree highlighted in red.

**Figure S22:**
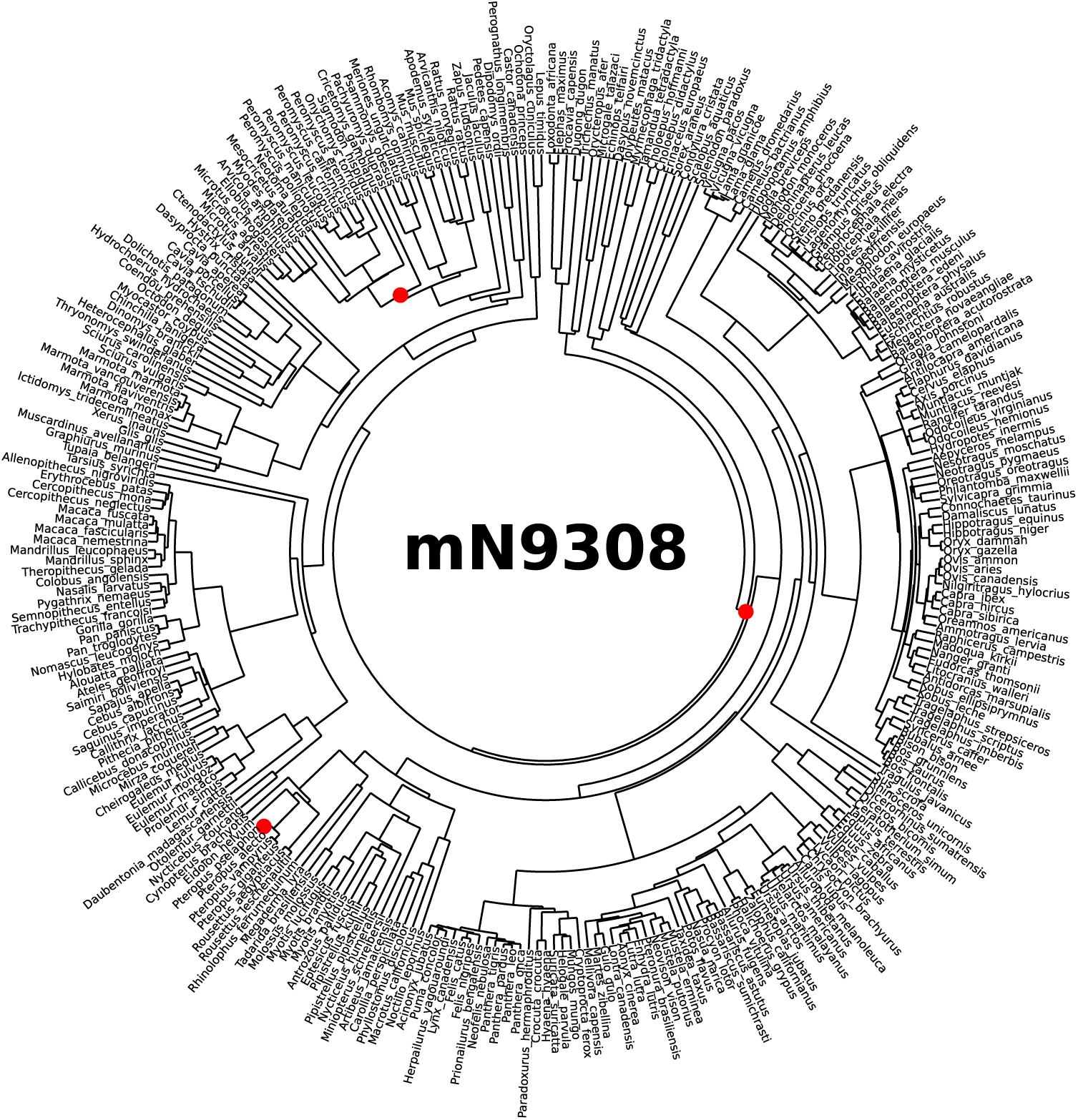
Mammal node-dated tree 9308 (mN9308), with nodes involved in topology differences from the main tree highlighted in red.

**Figure S23:**
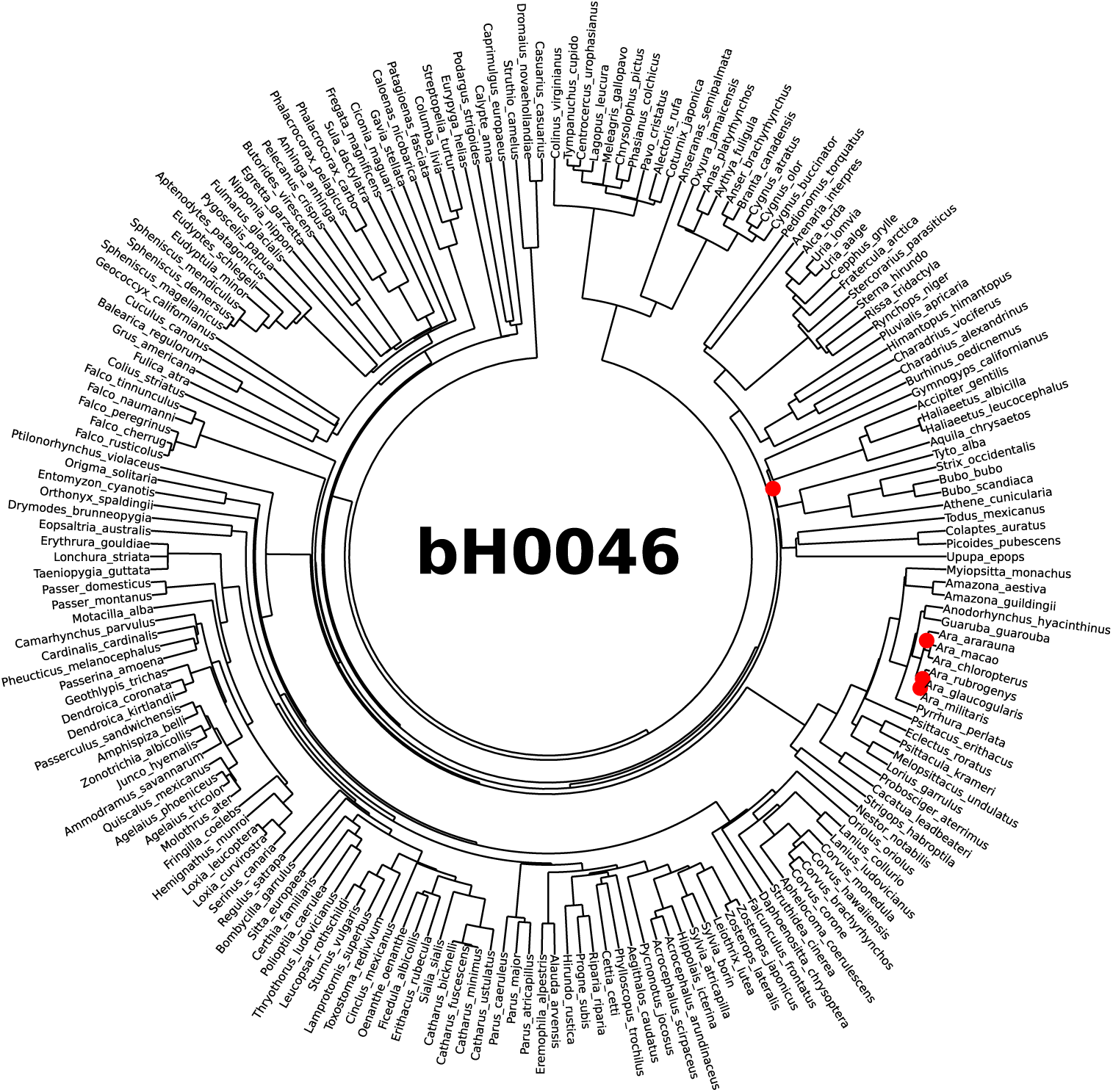
Bird Hackett backbone tree 0046 (bH0046), with nodes involved in topology differences from the main tree highlighted in red.

**Figure S24:**
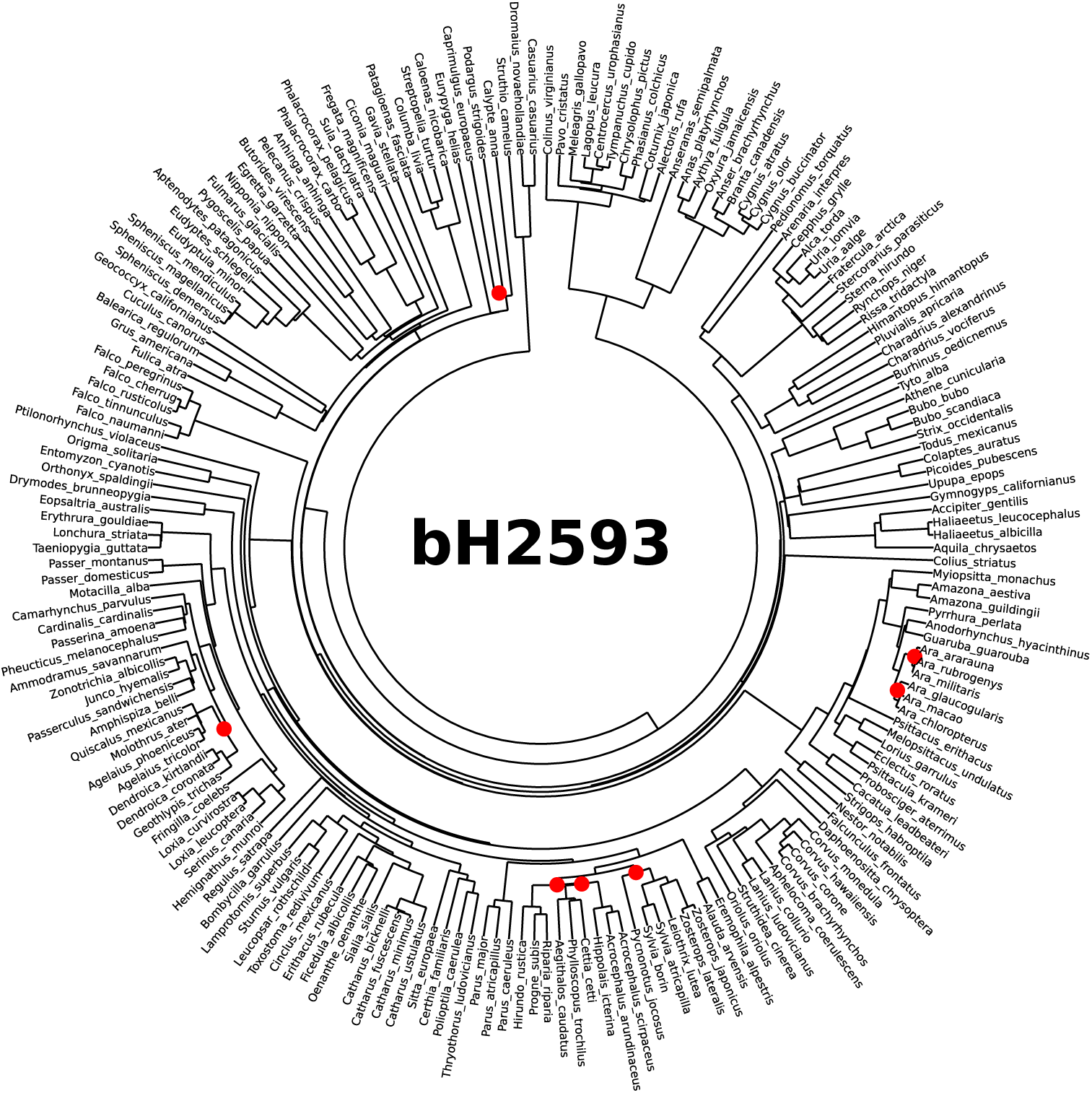
Bird Hackett backbone tree 2593 (bH2593), with nodes involved in topology differences from the main tree highlighted in red.

**Figure S25:**
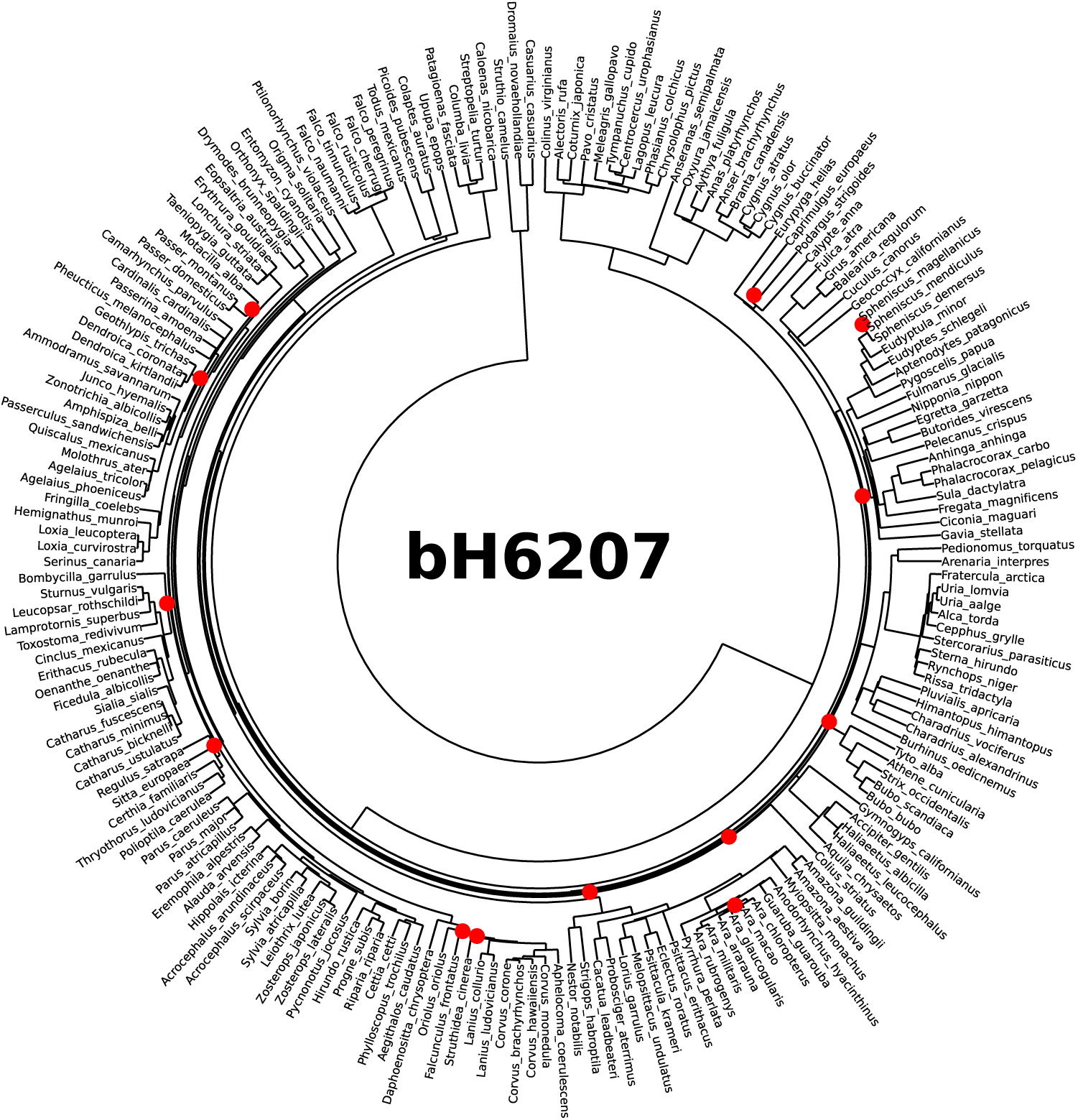
Bird Hackett backbone tree 6207 (bH6207), with nodes involved in topology differences from the main tree highlighted in red.

**Figure S26:**
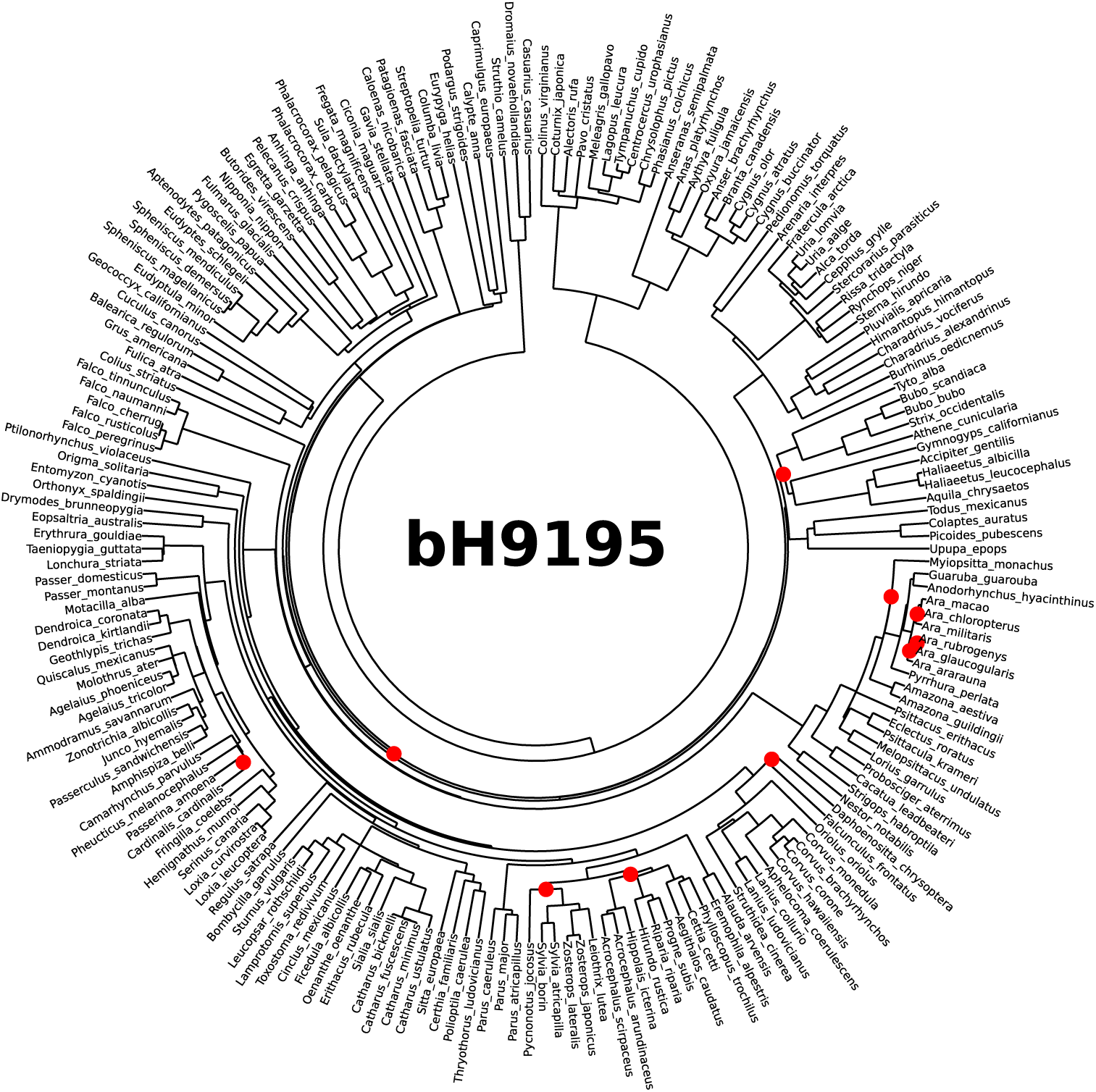
Bird Hackett backbone tree 9195 (bH9195), with nodes involved in topology differences from the main tree highlighted in red.

**Figure S27:**
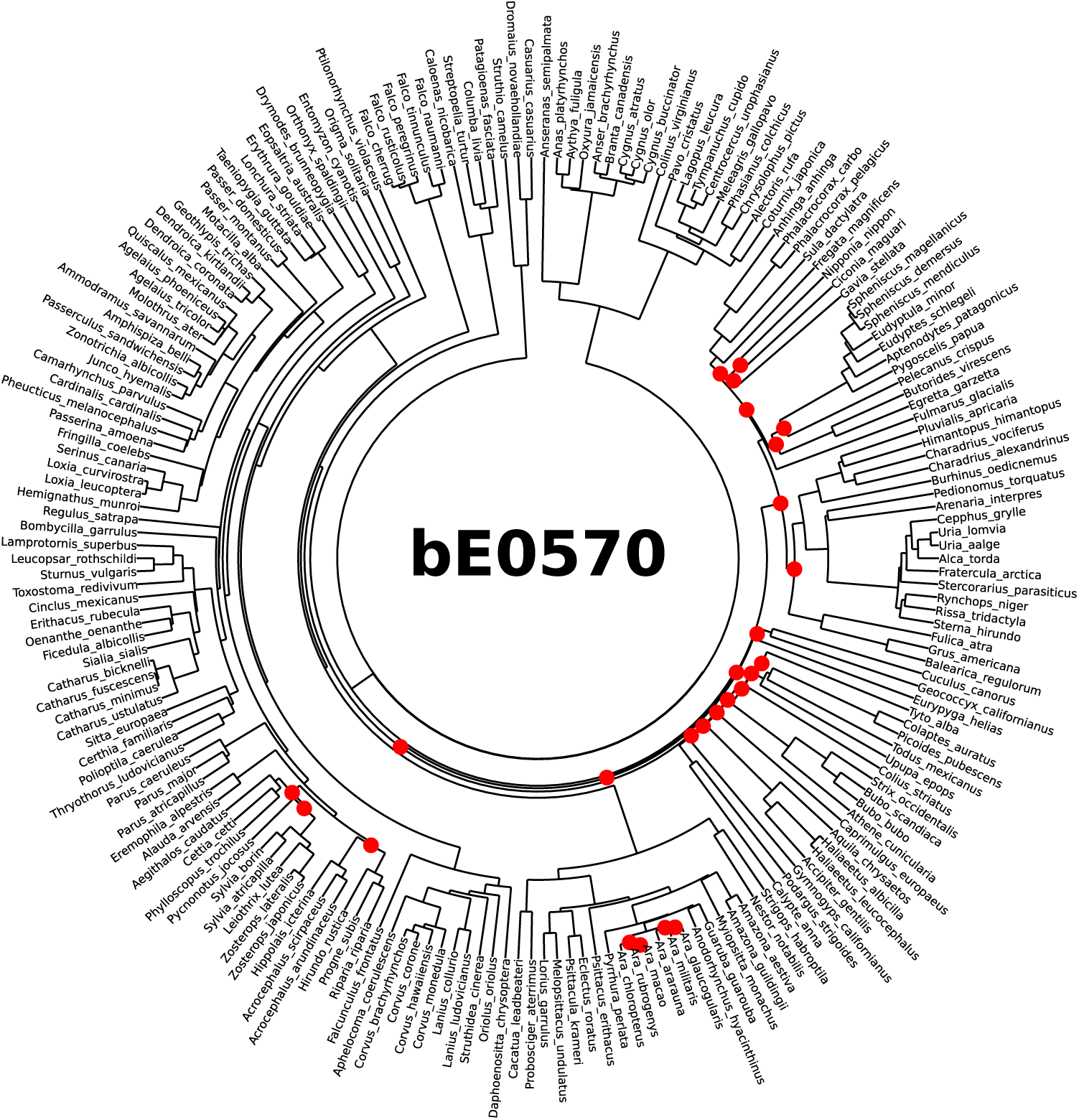
Bird Ericson backbone tree 0570 (bE0570), with nodes involved in topology differences from the main tree highlighted in red.

**Figure S28:**
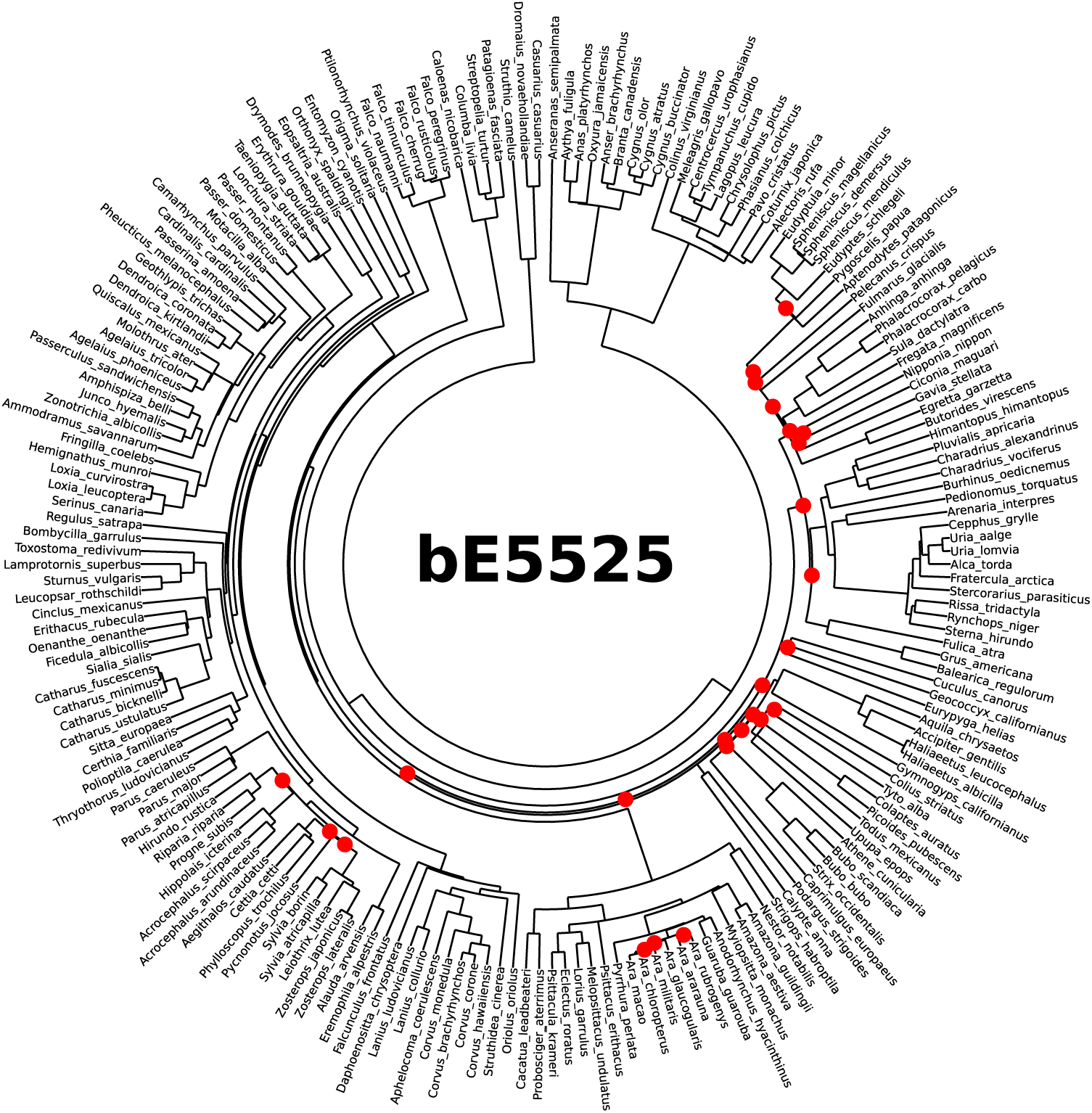
Bird Ericson backbone tree 5525 (bE5525), with nodes involved in topology differences from the main tree highlighted in red.

**Figure S29:**
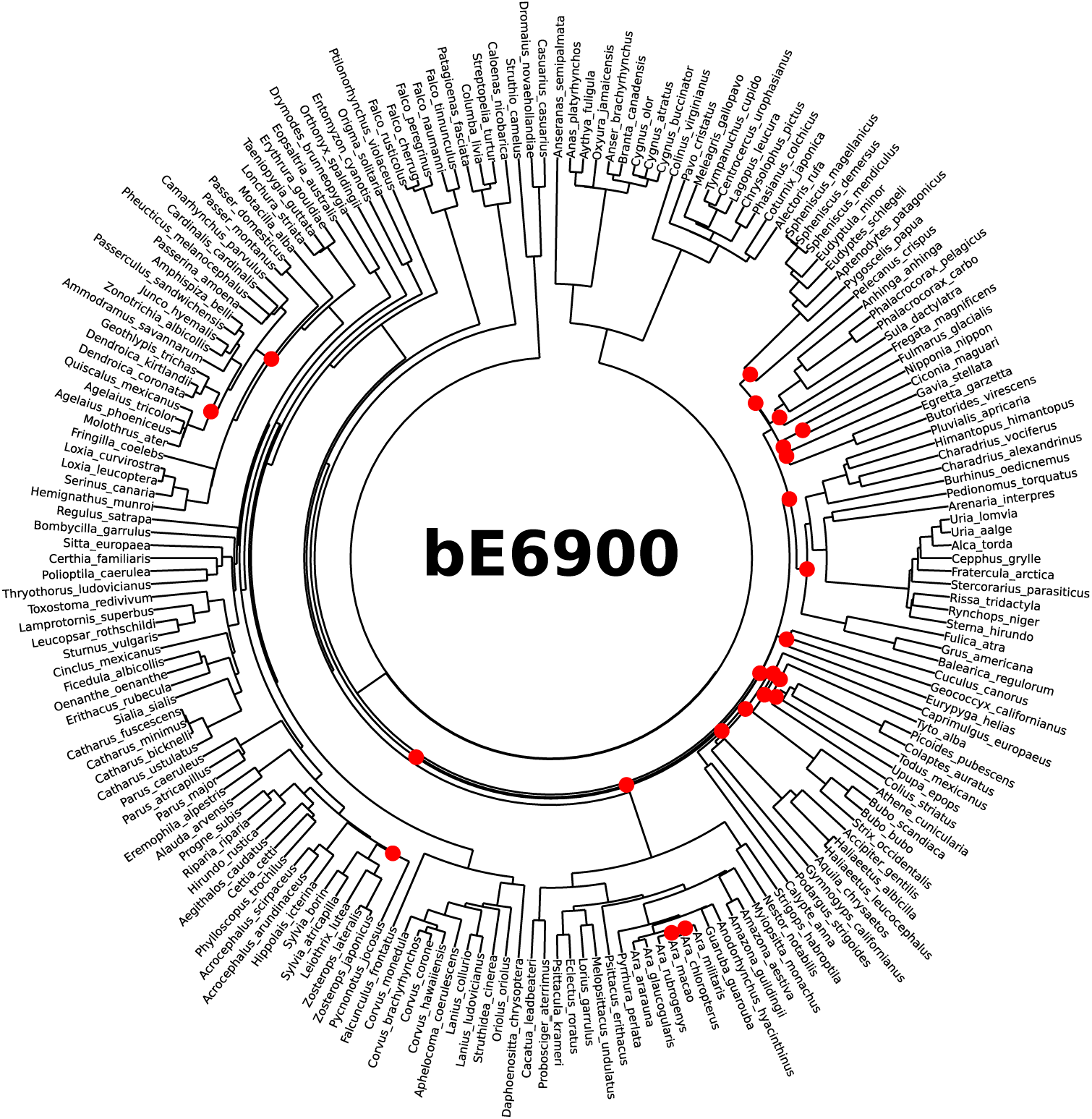
Bird Ericson backbone tree 6900 (bE6900), with nodes involved in topology differences from the main tree highlighted in red.

**Figure S30:**
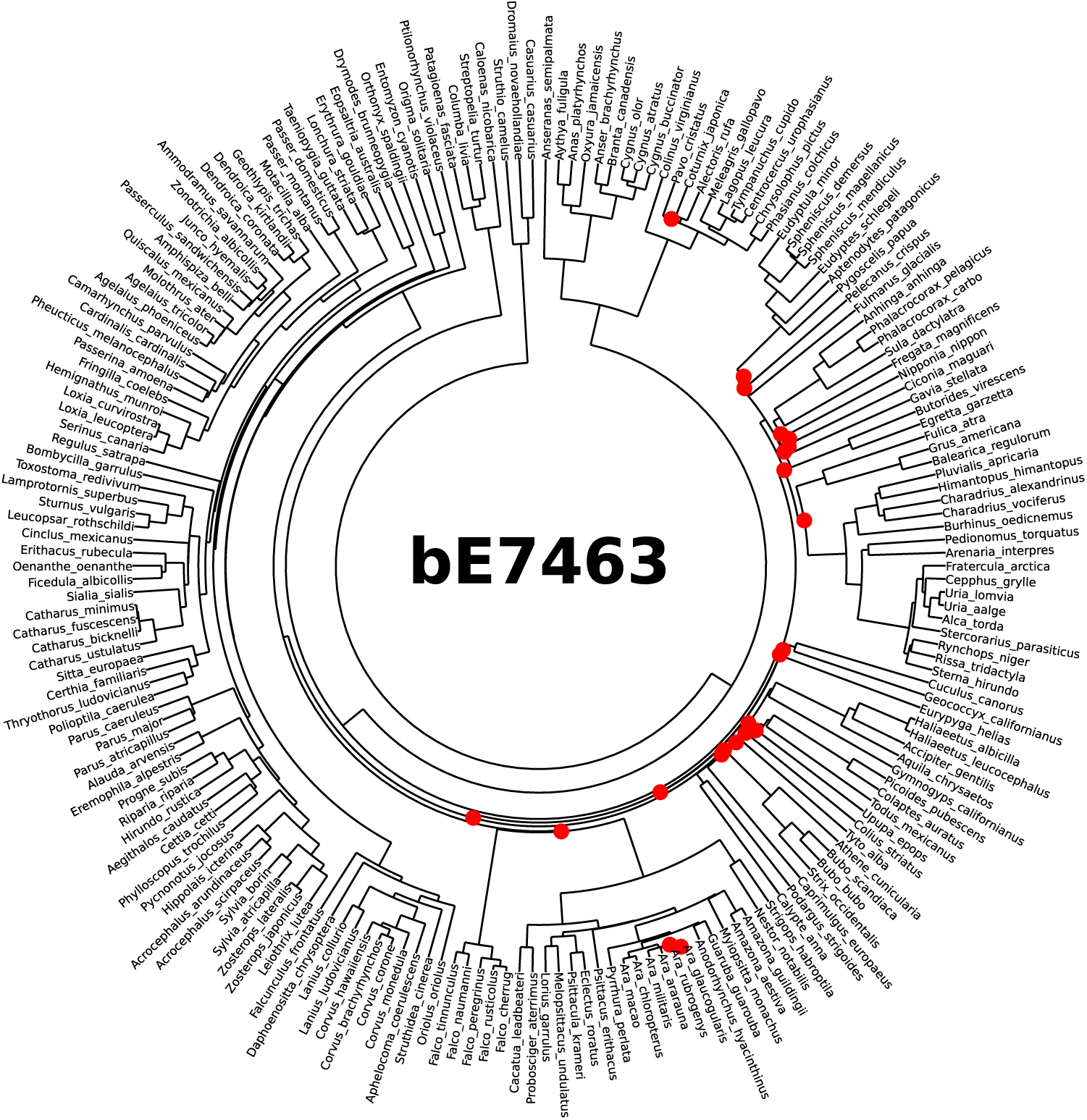
Bird Ericson backbone tree 7463 (bE7463), with nodes involved in topology differences from the main tree highlighted in red.

**Figure S31:**
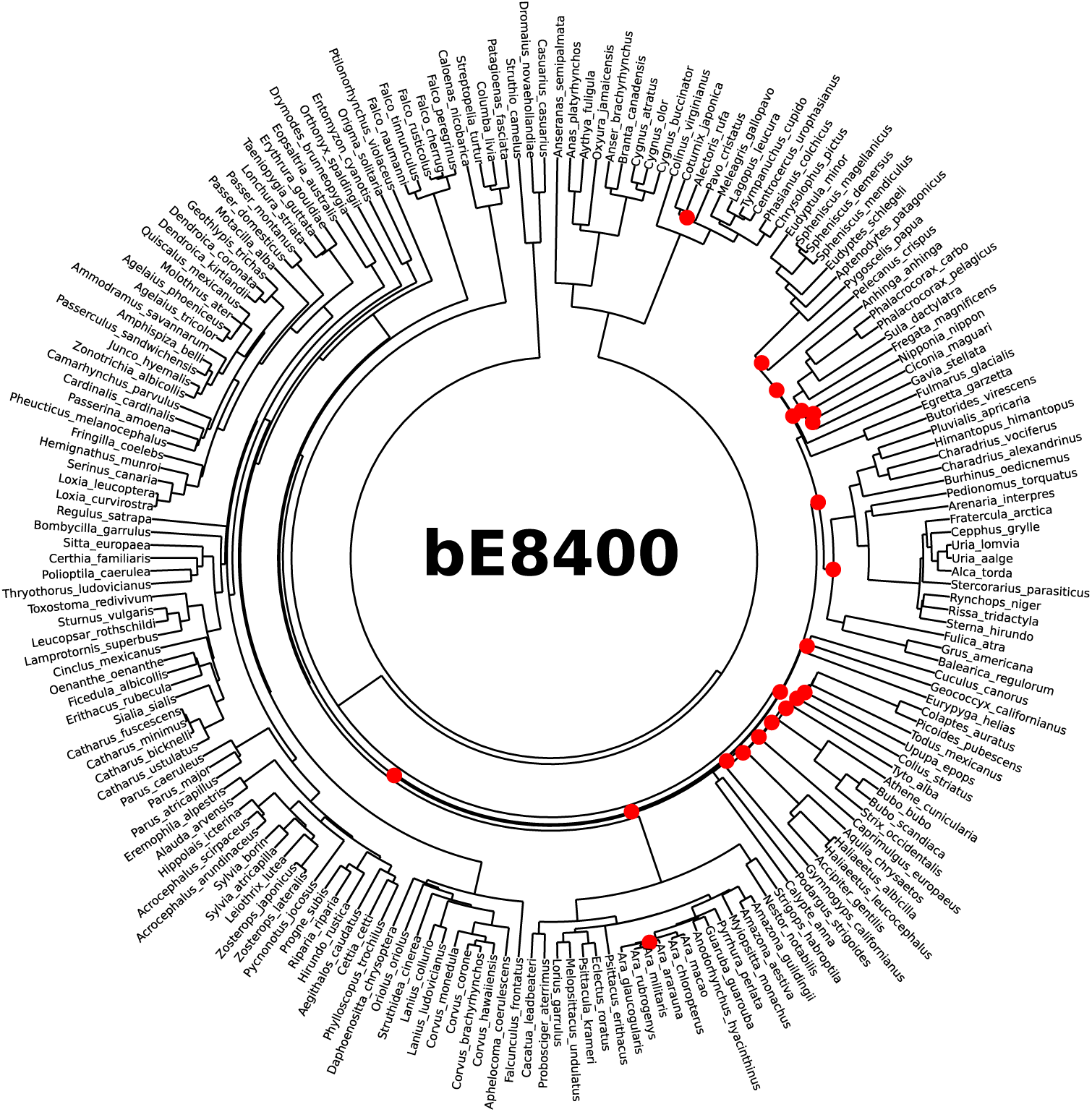
Bird Ericson backbone tree 8400 (bE8400), with nodes involved in topology differences from the main tree highlighted in red.

**Figure S32:**
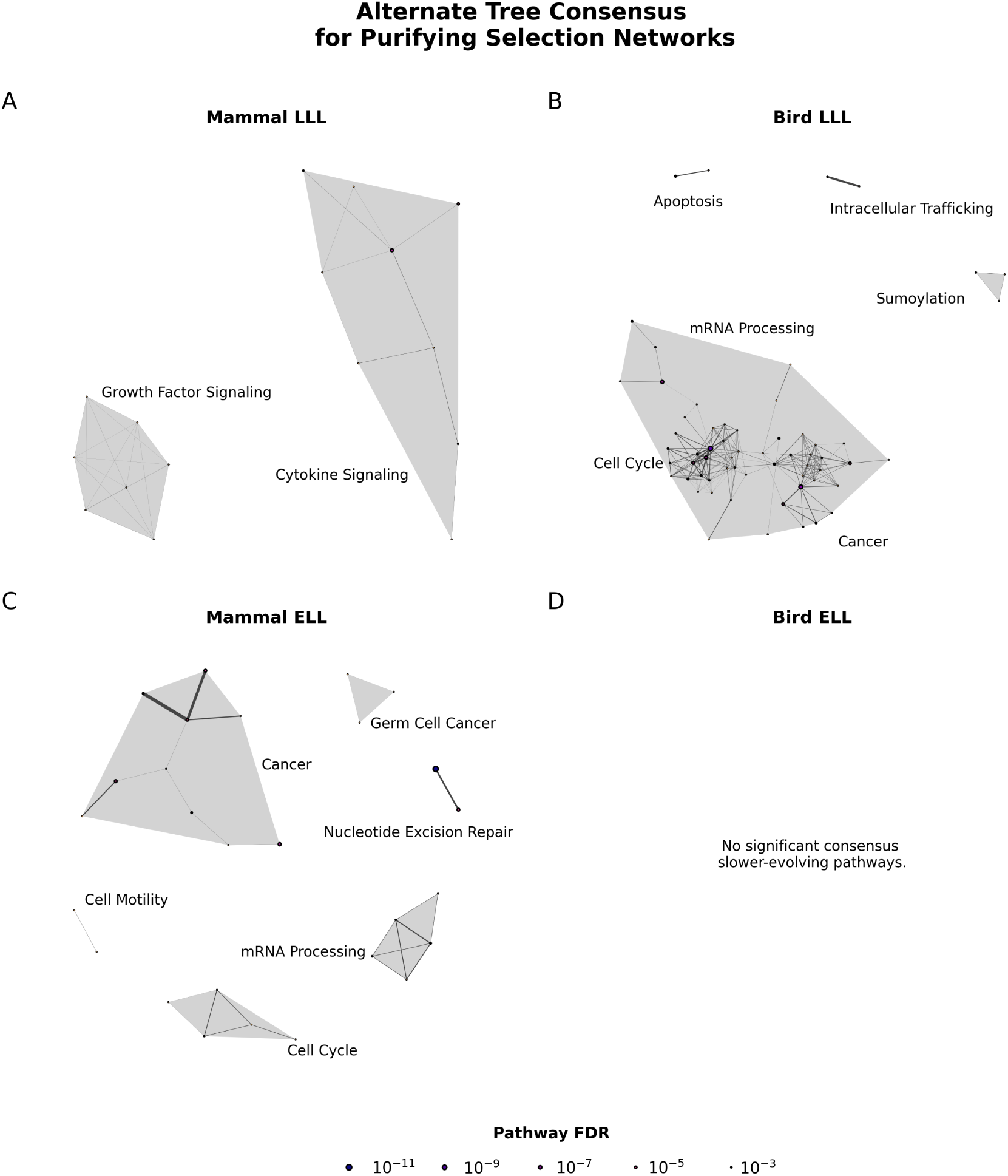
Consensus pathways under purifying selection in long-lived species in all trees analyzed. This figure includes only connected pathways with FDR < 0.05 *in all 9 alternate trees as well as the main tree, whose results are shown in* Figure 4*. The pathway “MEISSNER BRAIN HCP WITH H3K27ME3”, a significant and related (though unconnected) pathway in the main ELL birds analysis in Figure S37, is also significant in the alternate trees bE5525, bE7463, bE8400, bH9195*.

**Figure S33:**
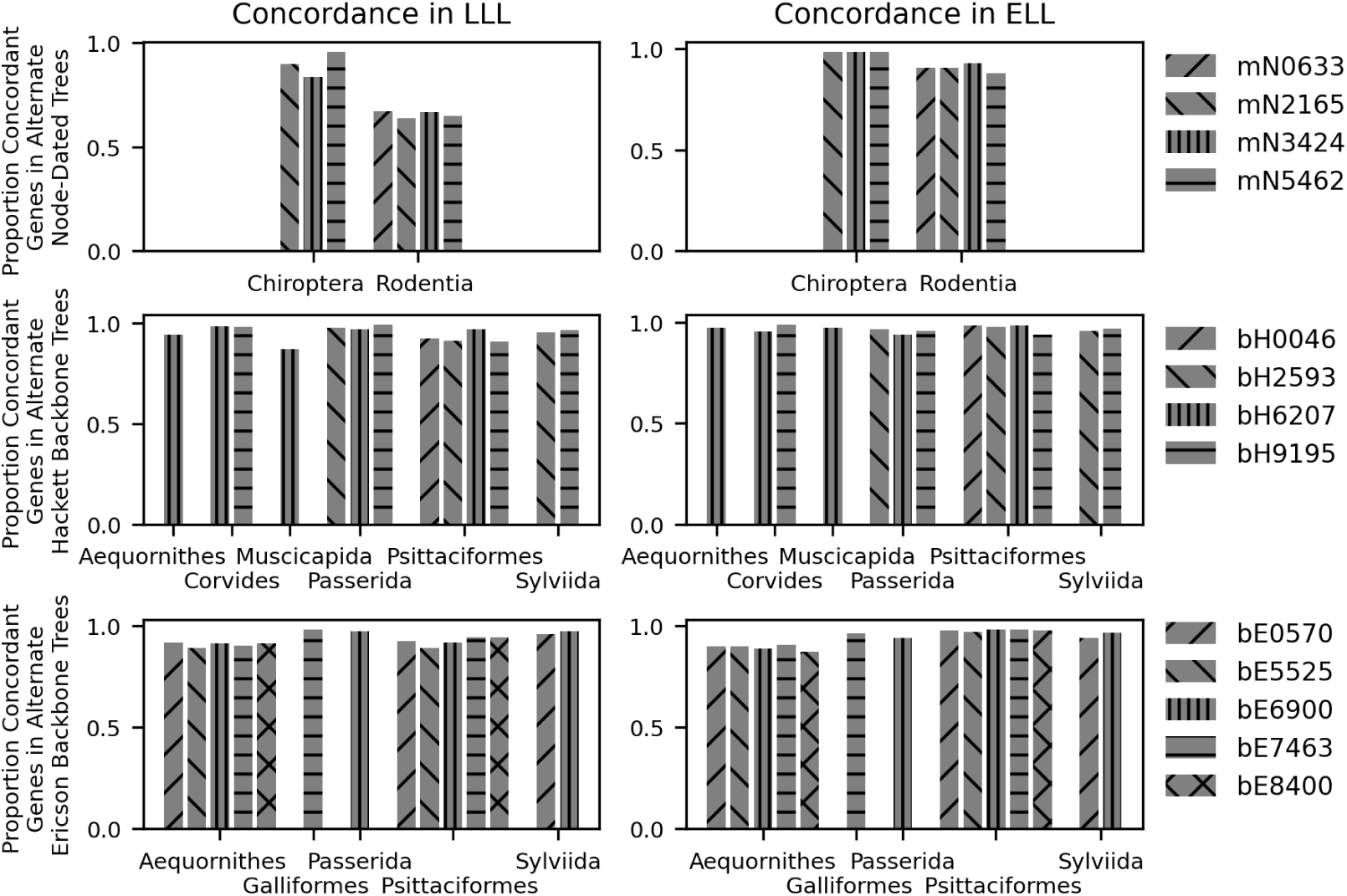
Concordance between positive selection results from our main mammal and bird trees and those derived from alternate topologies. *Note that alternate tree results for Rodentia LLL analyses, despite having lower concordance with the main analysis, have similar number of significant genes overall, with mN0633, mN2165, mN3424, and mN5462 having 213, 199, 206, and 208 significant genes (adjusted p <* 0.05 *by the Benjamini-Hochberg method), respectively, compared to 205 in the main tree analysis*.

**Table S26:**
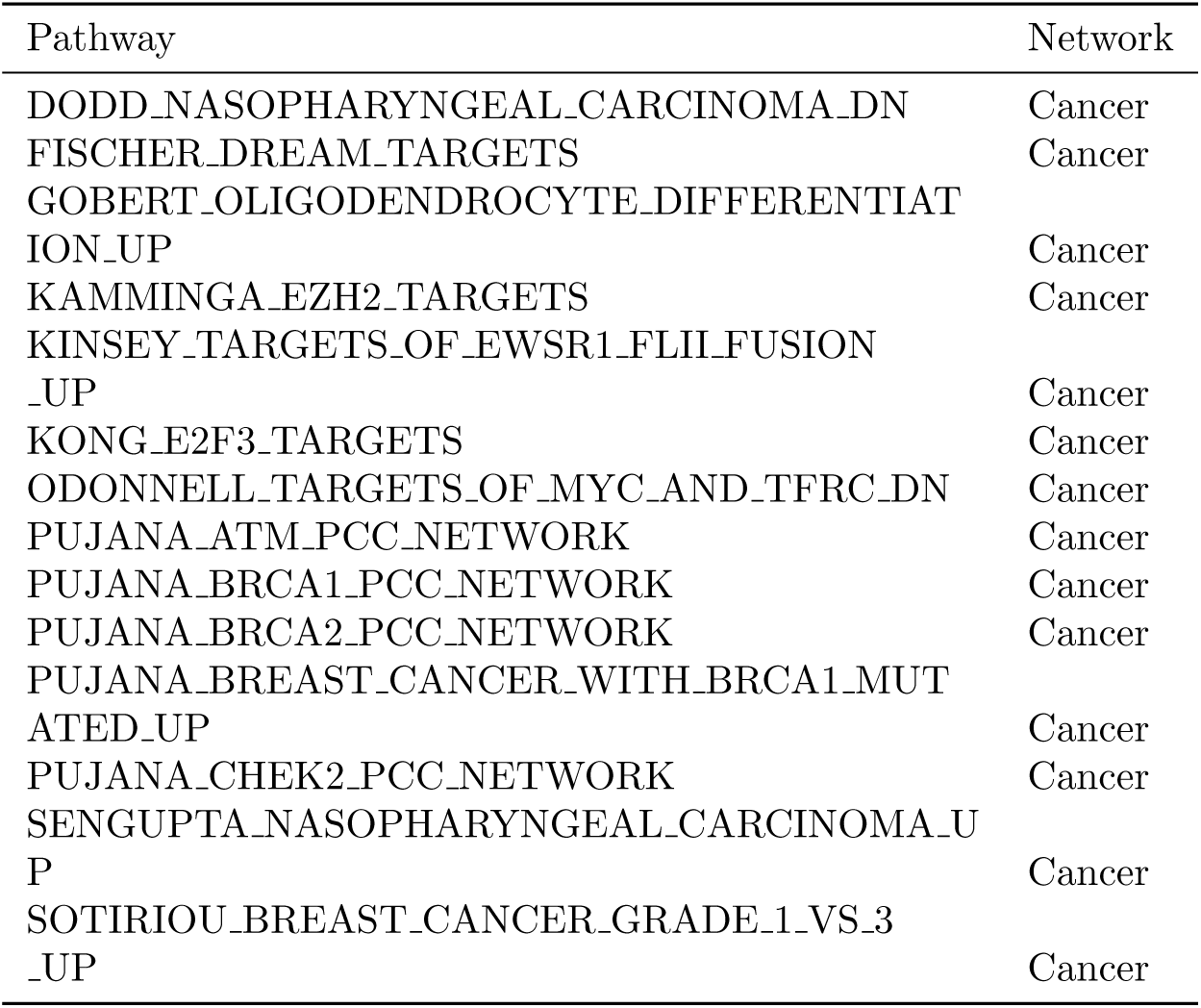
Connected pathways classified as belonging to the Cancer network.

**Table S27:**
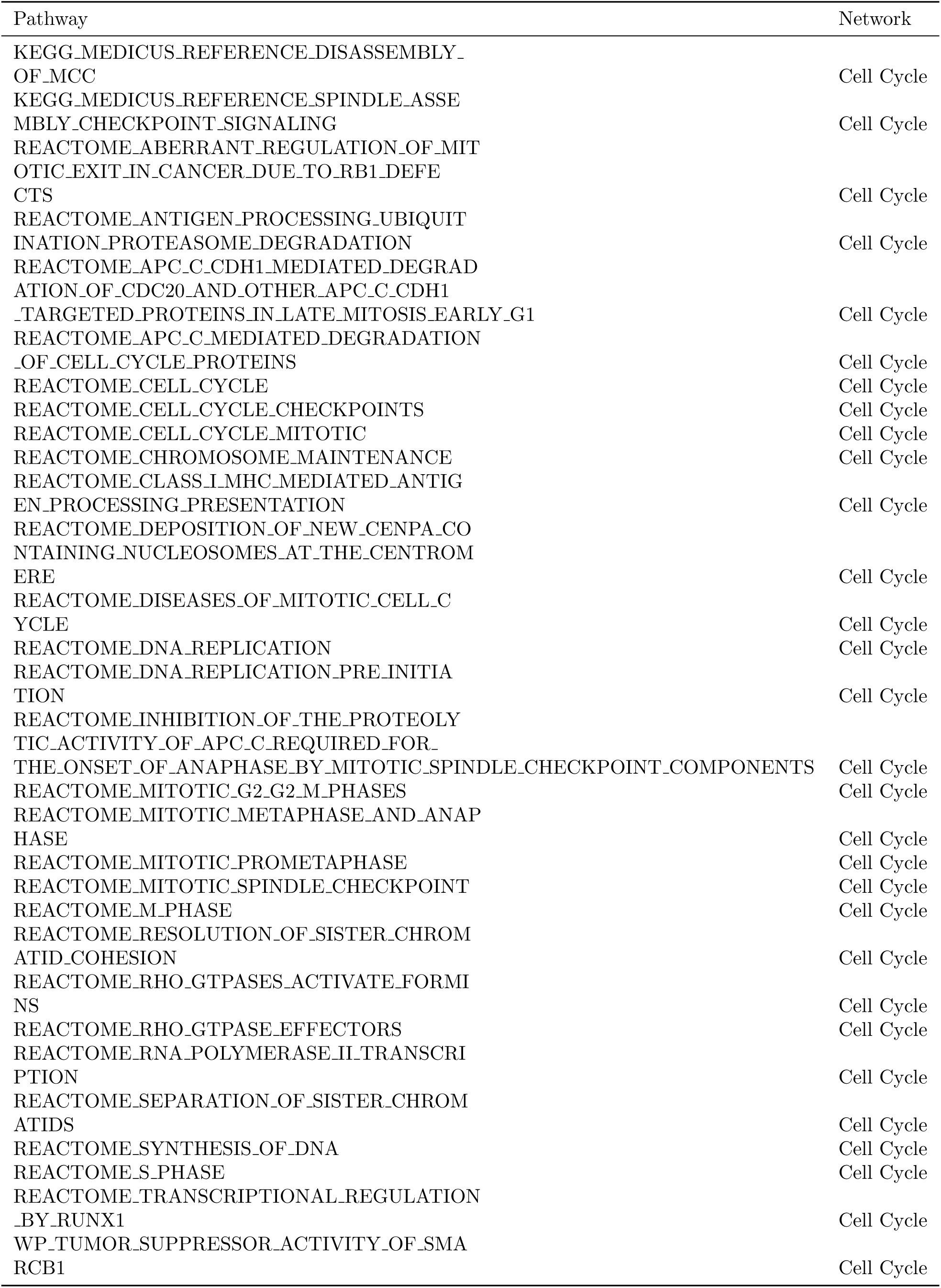
Connected pathways classified as belonging to the Cell Cycle network.

**Table S28:**
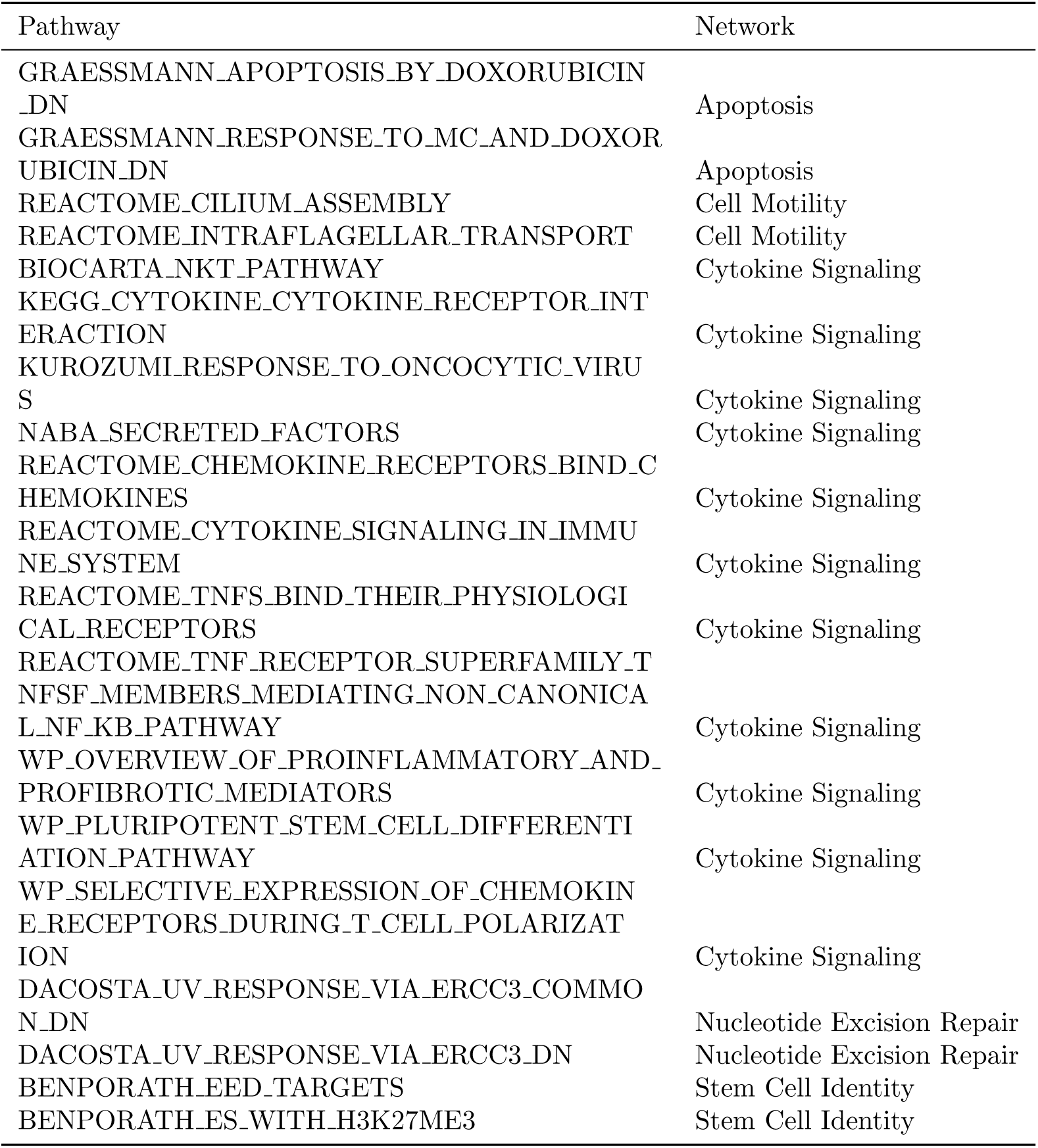
Connected pathways classified as belonging to the Apoptosis, Cell Motility, Cytokine Signaling, Nucleotide Excision Repair, or Stem Cell Identity networks.

**Table S29:**
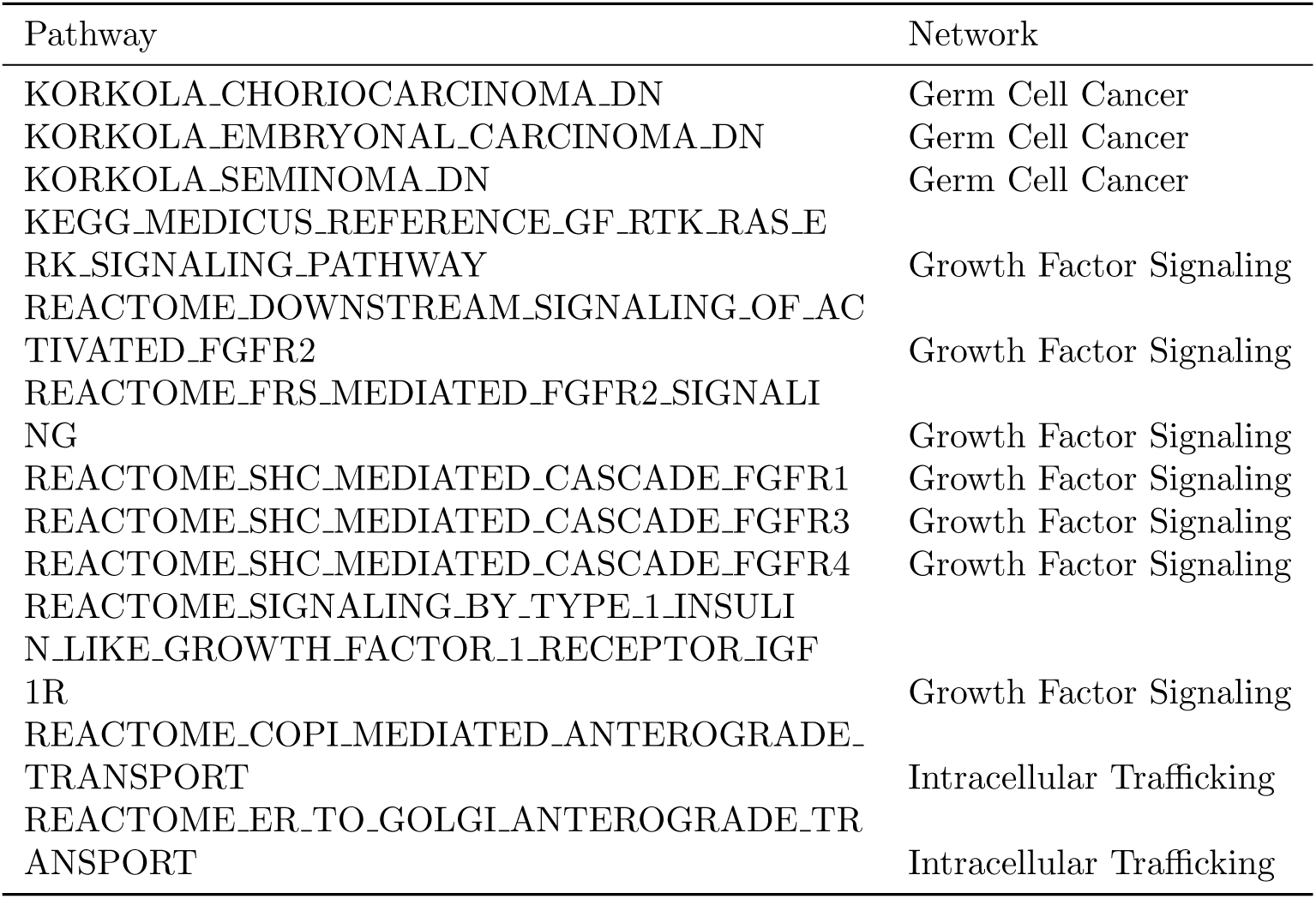
Connected pathways classified as belonging to the Germ Cell Cancer, Growth Factor Signaling, or Intra-cellular Trafficking networks.

**Table S30:** Connected pathways classified as belonging to the Infection Response, Sumoylation, or mRNA Processing networks.

| Pathway | Network |
| --- | --- |
| REACTOME_GENE_SILENCING_BY_RNA | Infection Response |
| REACTOME_HIV_INFECTION | Infection Response |
| REACTOME_HOST_INTERACTIONS_OF_HIV_F<br>ACTORS | Infection Response |
| REACTOME_INFLUENZA_INFECTION | Infection Response |
| REACTOME_PIWI_INTERACTING_RNA_PIRNA<br>_BIOGENESIS | Infection Response |
| REACTOME_TRANSCRIPTIONAL_REGULATION<br>_BY_SMALL_RNAS | Infection Response |
| REACTOME_SUMOYLATION | Sumoylation |
| REACTOME_SUMOYLATION_OF_CHROMATIN_O<br>RGANIZATION_PROTEINS | Sumoylation |
| REACTOME_SUMOYLATION_OF_DNA_DAMAGE_<br>RESPONSE_AND_REPAIR_PROTEINS | Sumoylation |
| CHNG_MULTIPLE_MYELOMA_HYPERPLOID_UP | mRNA Processing |
| KEGG_MEDICUS_REFERENCE_TRANSLATION_<br>INITIATION | mRNA Processing |
| KEGG_RIBOSOME | mRNA Processing |
| KEGG_SPLICEOSOME | mRNA Processing |
| LEE_METASTASIS_AND_RNA_PROCESSING_U<br>P | mRNA Processing |
| REACTOME_CELLULAR_RESPONSES_TO_STIM<br>ULI | mRNA Processing |
| REACTOME_CELLULAR_RESPONSE_TO_STARV<br>ATION | mRNA Processing |
| REACTOME_EUKARYOTIC_TRANSLATION_ELO<br>NGATION | mRNA Processing |
| REACTOME_EUKARYOTIC_TRANSLATION_INI<br>TIATION | mRNA Processing |
| REACTOME_METABOLISM_OF_RNA | mRNA Processing |
| REACTOME_MITOCHONDRIAL_TRANSLATION | mRNA Processing |
| REACTOME_MRNA_SPLICING | mRNA Processing |
| REACTOME_PROCESSING_OF_CAPPED_INTRO<br>N_CONTAINING_PRE_MRNA | mRNA Processing |
| REACTOME_RESPONSE_OF_EIF2AK4_GCN2_T<br>O_AMINO_ACID_DEFICIENCY | mRNA Processing |
| REACTOME_RRNA_PROCESSING | mRNA Processing |
| REACTOME_SELENOAMINO_ACID_METABOLIS<br>M | mRNA Processing |
| REACTOME_SRP_DEPENDENT_COTRANSLATIO<br>NAL_PROTEIN_TARGETING_TO_MEMBRANE | mRNA Processing |
| REACTOME_TRANSLATION | mRNA Processing |
| WP_MRNA_PROCESSING | mRNA Processing |

**Figure S34:**
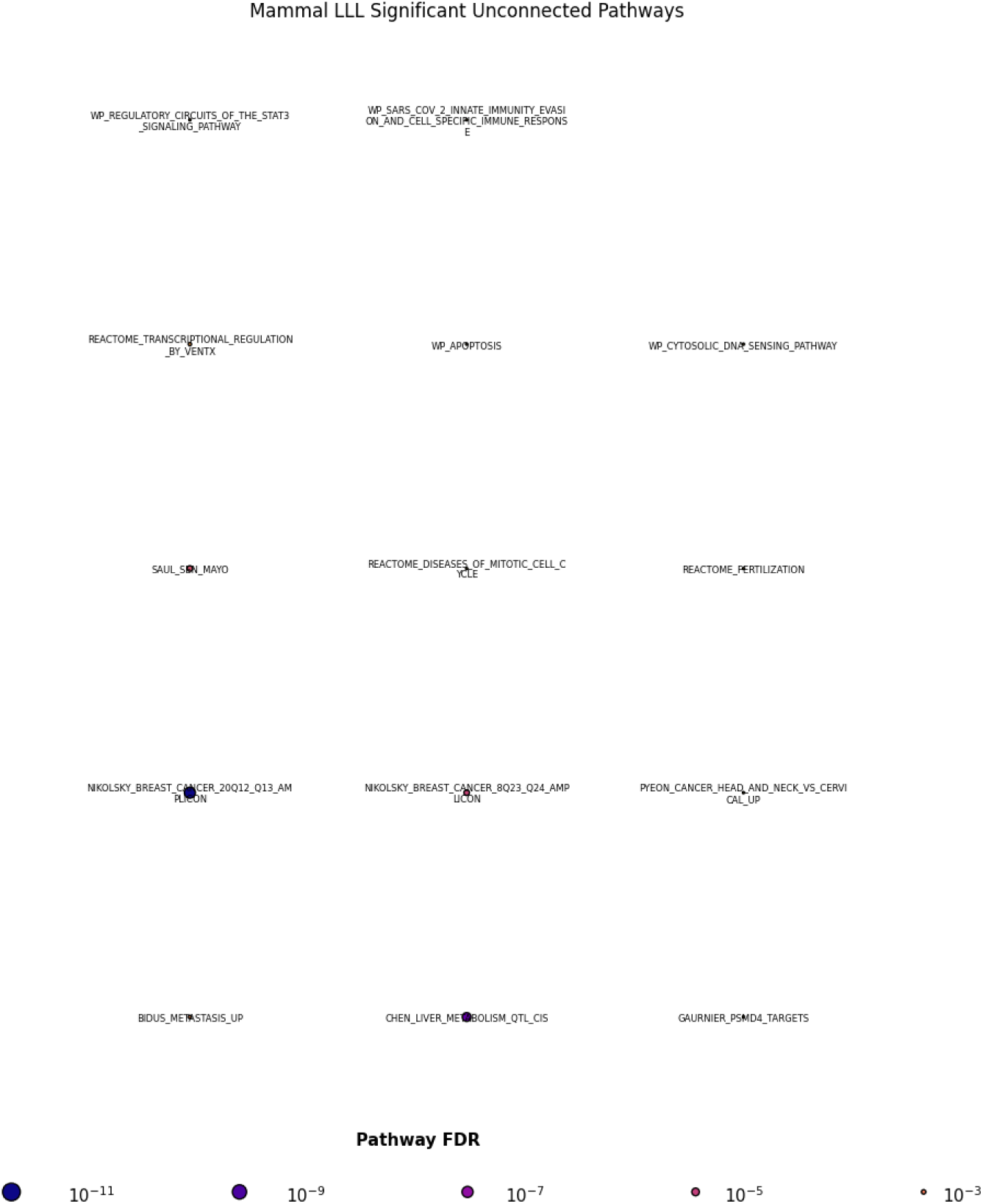
Unconnected pathways under significant constraint in LLL mammals. These pathways are themselves statistically significant at the FDR <0.05 level, but do not connect (shared genes >70% of smaller pathway’s gene set) to any other significant pathway.

**Figure S35:**
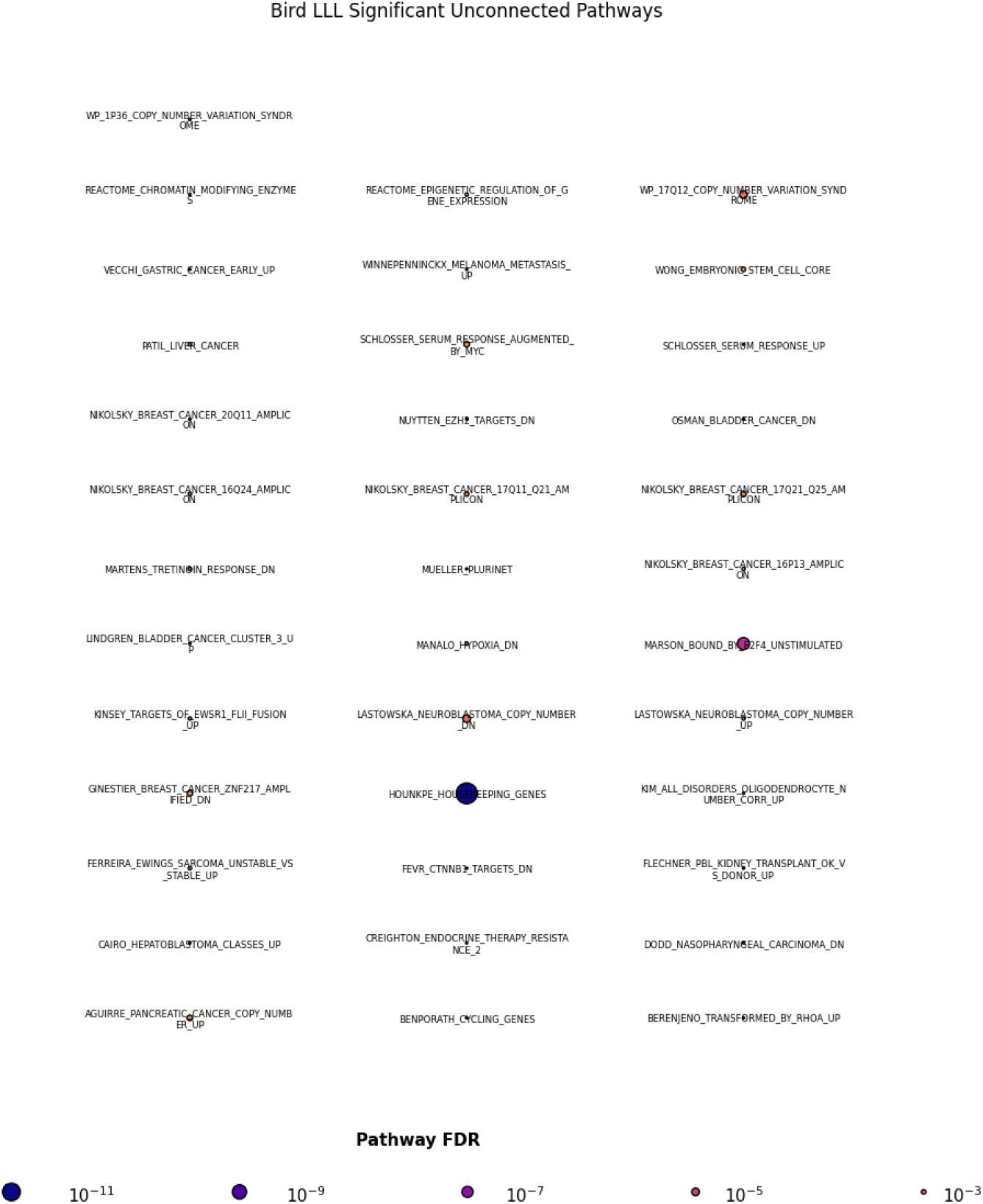
Unconnected pathways under significant constraint in ELL mammals. These pathways are themselves statistically significant at the FDR <0.05 level, but do not connect (shared genes >70% of smaller pathway’s gene set) to any other significant pathway.

**Figure S36:**
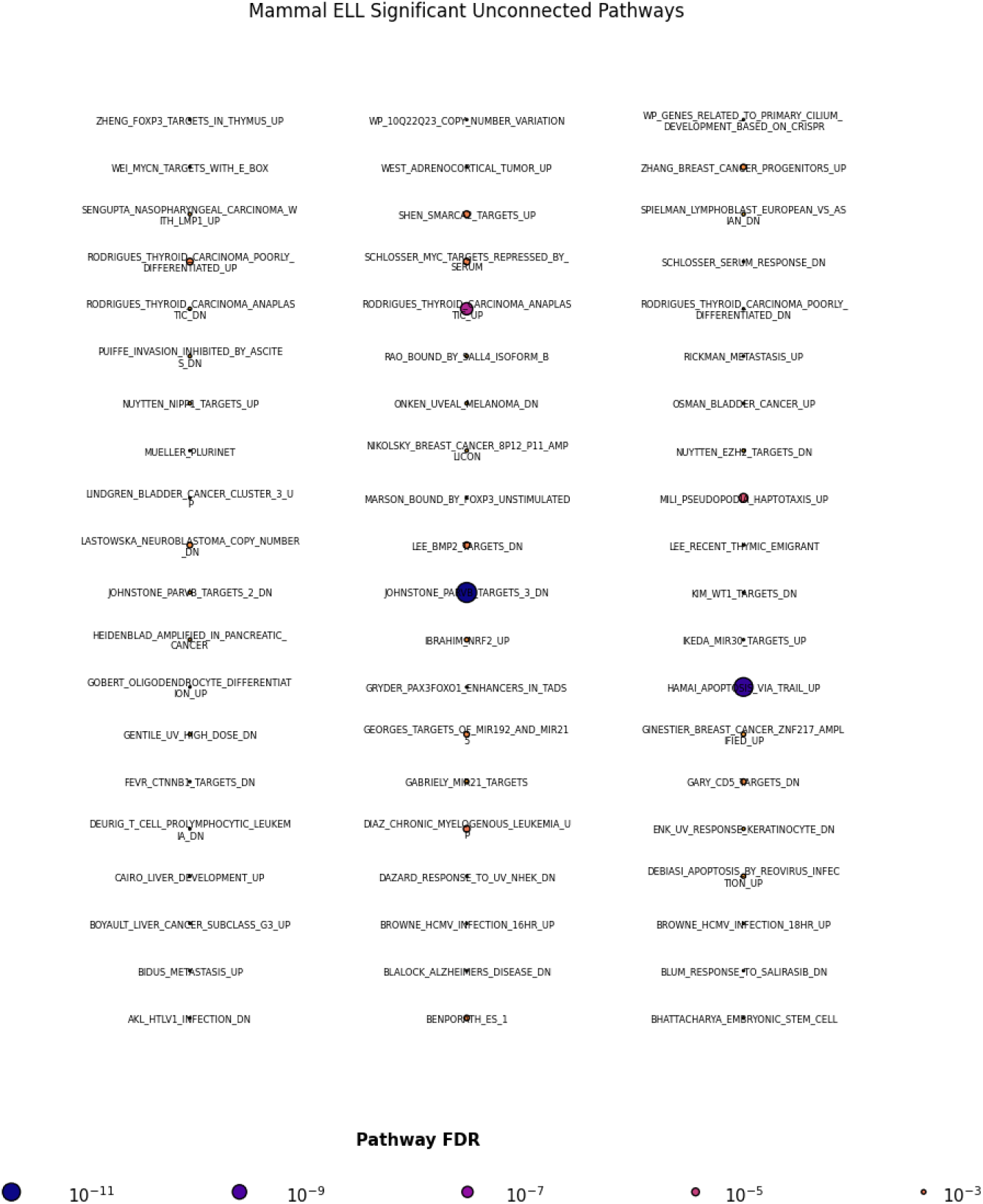
Unconnected pathways under significant constraint in LLL birds. These pathways are themselves statistically significant at the FDR <0.05 level, but do not connect (shared genes >70% of smaller pathway’s gene set) to any other significant pathway.

**Figure S37:**
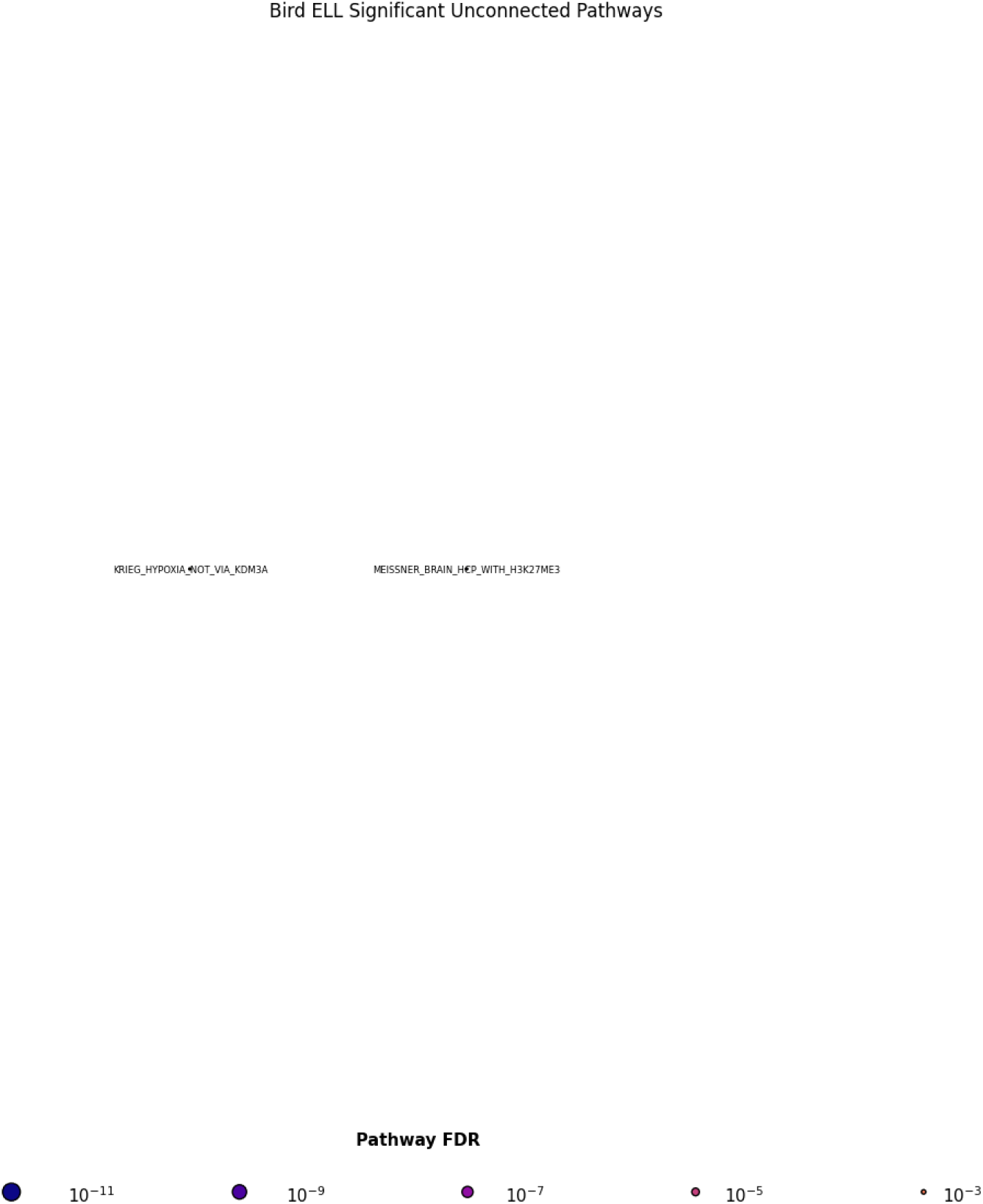
Unconnected pathways under significant constraint in LLL birds. These pathways are themselves statistically significant at the FDR <0.05 level, but do not connect (shared genes >70% of smaller pathway’s gene set) to any other significant pathway.

**Table S31:** Sources of silhouettes from PhyloPic. These were re-colored and used in Figure 6 to illustrate clades in which particular genes were positively selected in long-lived species. Most images used are in the public domain, with the exceptions of the representative icons for Chiroptera and Muscicapida (Attribu-tion 3.0 Unported: https://creativecommons.org/licenses/by/3.0/), and Corvides (Attribution 4.0 International: https://creativecommons.org/licenses/by/4.0/).

| Clade | Species Depicted | Artist |
| --- | --- | --- |
| Rodentia | *Marmota monax* | T. Michael Keesey |
| Ruminantia | *Bos bison* | Tracy Heath |
| Carnivora | *Panthera tigris altaica* | Steven Traver |
| Primates | *Gorilla gorilla* | Margot Michaud |
| Chiroptera | *Rousettus aegyptiacus* | Melissa Ingala |
| Cetacea | *Balaenoptera novaeangliae* | Guillaume Dera |
| Perissodactyla | *Rhinoceros unicornis* | Christophe Mallet |
| Aequornithes | *Pygoscelis papua* | Ferran Sayol |
| Corvides | *Corvus corone* | Stephen James McWilliam |
| Sylviida | *Alauda arvensis* | Pearson Scott Foresman |
| Muscicapida | *Sturnus vulgaris* | Maxime Dahirel |
| Passerida | *Agelaius phoeniceus* | Andy Wilson |
| Psittaciformes | *Anodorhynchus hyacinthinus* | Zitan Song |
| Charadriiformes | *Uria aalge* | Carbo |
| Galliformes | *Centrocercus urophasianus* | Richard Rich |
| Anseriformes | *Cygnus buccinator* | Xavier Giroux-Bougard |

